# Cells collectively reshape habitability of temperature by helping each other replicate

**DOI:** 10.1101/726463

**Authors:** Diederik S. Laman Trip, Hyun Youk

## Abstract

How the rising global temperatures affect organisms is a timely question. The conventional view is that high temperatures cause microbes to replicate slowly or die, both autonomously. Yet, microbes co-exist as a population, raising the underexplored question of whether they can cooperatively combat rising temperatures. Here we show that, at high temperatures, budding yeasts help each other and future generations of cells replicate by secreting and extracellularly accumulating glutathione - a ubiquitous heat-damage-reducing antioxidant. Yeasts thereby collectively delay and can halt population extinctions at high temperatures. As a surprising consequence, even for the same temperature, a yeast population can either exponentially grow, never grow, or grow after unpredictable durations (hours-to-days) of stasis, depending on its population density. Despite the conventional theory stating that heat-shocked cells autonomously die and cannot stop population extinctions, we found that non-growing yeast-populations at high temperatures - due to cells cooperatively accumulating extracellular glutathione - continuously decelerate and can eventually stop their approach to extinction, with higher population-densities stopping faster. We show that exporting glutathione, but not importing, is required for yeasts to survive high temperatures. Thus, cooperatively eliminating harmful extracellular agents – not glutathione’s actions inside individual cells – is both necessary and sufficient for surviving high temperatures. We developed a mathematical model - which is generally applicable to any cells that cooperatively replicate by secreting molecules - that recapitulates all these features. These results show how cells can cooperatively extend boundaries of life-permitting temperatures.

---

Many model organisms are typically studied at a particular temperature that is deemed “optimal” for that organism’s growth. For example, the budding yeast is often studied at 30 °C while *E. coli* and mammalian cells are usually studied at 37 °C. Yet, an organism can live and replicate at a range of “habitable temperatures” (*1–3*). Understanding why an organism cannot replicate and eventually dies for temperatures outside of the habitable range – at “unlivable temperatures” - may provide insights into living systems’ vulnerabilities and how temperature, a fundamental physical quantity, drives processes of life. Moreover, given the rising global temperatures, a timely and practical question is how organisms do or can combat high temperatures to avoid extinction. For microbes and mammalian cells, the conventional view is that as the temperature increases above some optimal value within the habitable temperature regime and approaches unlivable temperatures, cells need more time to replicate and that once the temperature enters the unlivable temperature regime, cells fail to replicate and eventually die (*1–4*) (Figs. 1a-b). This view states that at sufficiently high temperatures, crucial proteins unfold and other heat-induced damages occur (e.g., damages by reactive oxygen species) (*5–7*), all of which disrupt cell replications. Moreover, it states that whether a cell can replicate or not at a high temperature depends on its autonomous ability to repair heat-induced damages by using its heat-shock response system, which is conserved across species (*8*). It is thought that while cells can autonomously repair heat-induced damages at moderately high, still-habitable temperatures that are just below the unlivable temperatures, they fail to do so at temperatures that are too high (i.e., at unlivable temperatures). Thus, this view states that one cell’s ability to replicate and its lifespan are both independent of any other cell’s lifespan and ability to replicate (*4*) (Fig. 1c). Yet a cell rarely exists alone - it lives within a population and can cooperate with other cells. Indeed, microbes can use mechanisms such as quorum sensing to coordinate their behaviors (*9*), share food (*10–12*), and collectively tune their extracellular pH (*13*). These findings motivated us to ask whether microbes use collective strategies to combat rising temperatures and if so, what such strategies are (Fig. 1d). These questions remain unaddressed for many microbial species. We sought to address them for the budding yeast, *Saccharomyces cerevisiae*. Surprisingly, we discovered that budding yeasts use a collective strategy to help each other replicate, help future generations of cells replicate, and thereby delay and even prevent population extinctions at high temperatures that one would conventionally define as being unlivable for the budding yeast.

**Figure 1.**
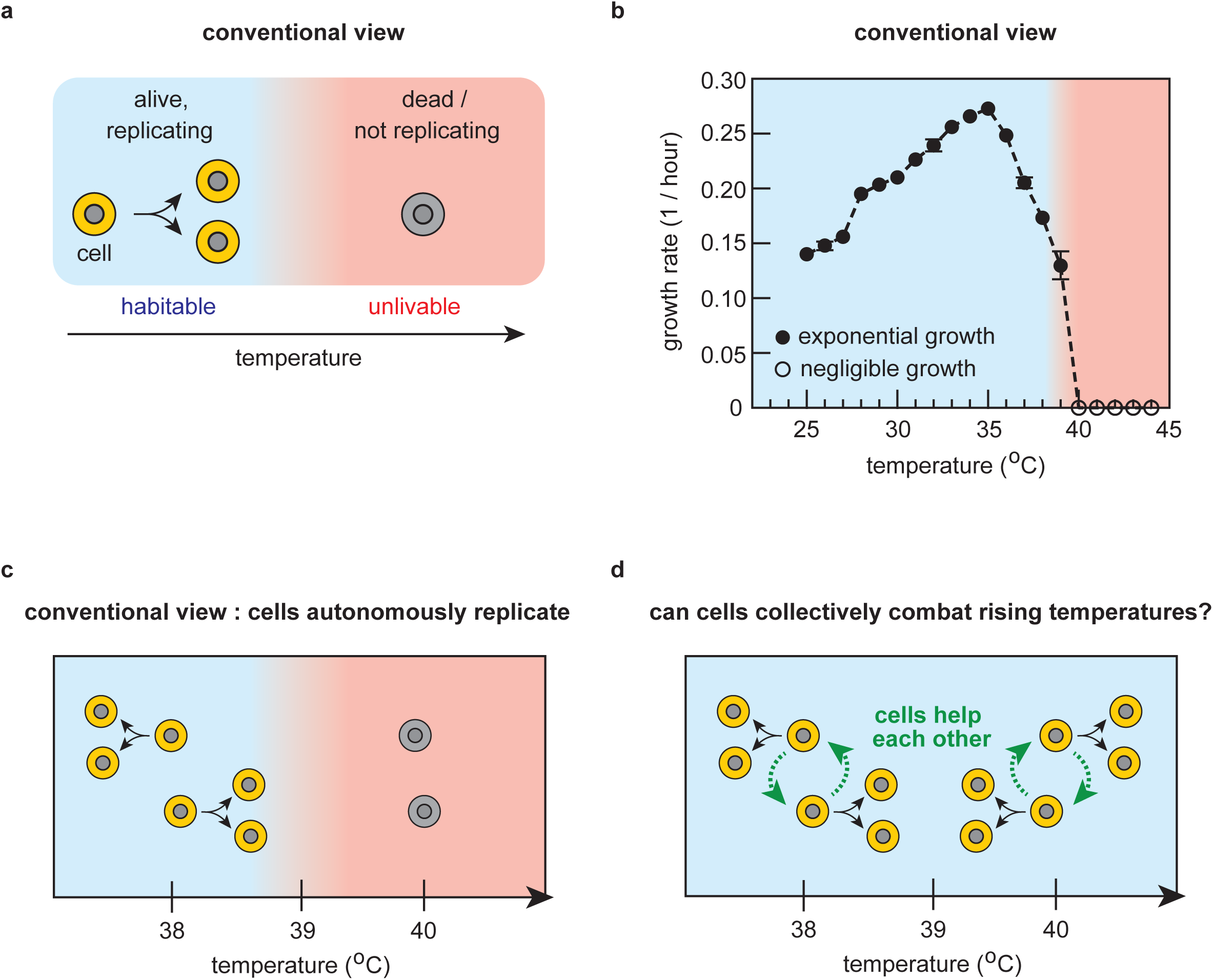
Conventional, cell-autonomous view of temperature-dependent cell-replications. (**a**) The conventional view states that cells autonomously replicate at “habitable temperatures” (blue region) and that at sufficiently high temperatures (i.e., “unlivable temperatures”), cells fail to replicate and can eventually die (red region). This view states that whether a cell can replicate or not at a given temperature depends on the cell’s autonomous ability to use its heat-shock response system to sufficiently deal with various heat-induced damages. (**b**) Growth rate as a function of temperature for populations of wild-type yeast cells. Black data points in the blue region are for populations with sustained, exponential growth over time and white data points in the red region are for populations without sustained exponential growth. 39 °C is near a boundary of blue and red regions (also see Supplementary Fig. 1). (**c**) The conventional view (explained in (a)) applied to budding yeast, based on the data in (b). (**d**) Question that we investigated in our study: can cells (budding yeasts) cooperatively combat rising temperatures so that they can turn a temperature that is unlivable for a few cells (e.g., 40 °C shown in (c)) into a habitable temperature if there are sufficiently many cells working together to fight off extinction?

As our starting point, we reproduced the well-known, textbook picture of how temperature affects the growth of yeast - and other microbes (*4,14–16*) - by using populations of a laboratory-standard (“wild-type”) strain of haploid budding yeast (Fig. 1b and Supplementary Fig. 1). According to this picture, the yeasts’ growth rate becomes zero - the population density negligibly changes - for temperatures of 40 °C and higher, which would thus be defined as “unlivable temperatures” for budding yeast (Fig. 1b - dark red region; Supplementary Fig. 1). Despite being evidently true - since we reproduced it here - we discovered that this cell-autonomous view of yeast replication (Fig. 1c) is misleading. For example, we discovered that yeast populations, depending on their population densities, can actually grow at “unlivable” temperatures (e.g., at 40 °C) and not grow at “habitable” temperatures (e.g., at 38 °C). As we will show, we discovered a picture that revises the cell-autonomous view: yeasts help each other and their future generations of cells replicate at high temperatures by secreting and extracellularly accumulating glutathione - a versatile antioxidant that is widely used by many species - which reduces damages caused by reactive oxygen species created by high temperatures. Thus, we found that yeasts collectively set the habitability of each temperature (Fig. 1d). A surprising consequence of this is that a temperature which is unlivable for relatively few yeasts becomes habitable if there are enough yeasts in a population that can help each other replicate. As another surprising consequence, we found that a yeast population - due to its cells gradually accumulating extracellular glutathione - decelerates and can eventually stop its approach to extinction at high temperatures. In fact, we found that a population of a higher density can more rapidly decelerate and halt its approach to extinction due to more cells cooperating through their secreted glutathione. This revises the prevalent theory which states that cells autonomously die at high temperatures. Intriguingly, we discovered that yeast populations at one particular temperature - a special temperature that places yeasts at a cusp of being able to replicate and unable to replicate - exhibit behaviors that are akin to those of biomolecular (*17,18*) and physical systems (*19*) that undergo phase transitions. We developed a mathematical model that recapitulates all the main results of our experiments. The model also quantitatively explains the origin of the phase-transition-like features by using, as the mechanism, cells cooperatively accumulating extracellular glutathione which promotes cell replications. Our paper ends by showing that glutathione’s extracellular action as an antioxidant, not its intracellular action, is solely responsible for promoting cell replications, thus highlighting an underappreciated extracellular role of glutathione for yeast. Glutathione, antioxidants, and heat-induced reactive oxygen species are common to many organisms, including humans. Hence the cooperative mechanism for yeast that we uncovered here - involving glutathione’s previously underappreciated role as a secreted factor at high temperatures – raises the possibility that cells of other species do or can also collectively combat rising temperatures in a similar manner. Taken together, our work shows how the habitability of a temperature emerges as a community-level property for a specie - determined by intraspecies interactions - rather than determined solely by the specie’s autonomous features.

## RESULTS

### Population density determines replicability of cells and habitability of each temperature

We re-examined the conventional, cell-autonomous picture by incubating populations of wild-type yeasts at a conventionally-defined habitable temperature (∼38 °C), unlivable temperature (∼40 °C), and a transition temperature in between the two (∼39 °C). We first grew the yeasts at 30 °C in 5 mL of standard minimal-media. We then transferred some of these cells to a fresh minimal media and then incubated them at a desired temperature, just as we did to obtain the conventional picture (Fig. 1b). This time, however, we took care to transfer a precise number of cells to the fresh minimal media so that we could vary the initial population-density (# of cells/mL) over a wide range. This is in contrast to how one obtains the conventional picture: one usually ignores the precise number of cells that a population begins with and, regardless of the temperature, keeps the initial population-density within a typical range of values that a plate reader can convert to an accessible range of optical densities (typically ∼10^6^ cells/mL in our experiments) (Supplementary Fig. 1). With a flow cytometer, we counted the integer numbers of cells per volume to determine the population density at each time point. For each temperature, we incubated multiple liquid cultures - these all started with the same population density – with cells that all came from a single liquid culture of cells that exponentially grew at 30 °C. These experiments revealed surprising behaviors that deviated from the conventional picture. Specifically, at the supposedly-habitable temperature of ∼38 °C, none of the populations that started with a relatively low population density (200 cells/mL) grew at all during ∼12 days of incubation except for a small, transient growth that occurred for a few hours right after the transfer from 30 °C (Fig. 2a - red curves). At the same temperature (∼38 °C), setting the initial population density to be just five times larger (1,000 cells/mL) than these non-growing populations yielded a population whose behavior was completely unpredictable: it could either grow until it reached the carrying capacity (i.e., ∼10^7^ cells/mL) or not grow at all after the initial transient-growth (Fig. 2a - green curves). When the population did grow, it could wait four days or eight days or some other, unpredictable time before starting to grow (Fig. 2a - multiple green curves). Still, at the same temperature (∼38 °C), setting the initial population density to be just five times larger (5,000 cells/mL) than these randomly-growing populations yielded populations that always grew exponentially over time, all in the same way, until they reached the carrying capacity (Fig. 2a - blue curves). Thus, among the three initial population densities at 38 °C, only the largest one led to the deterministic, population-level growth that the conventional picture states should be exhibited by every initial population-density at habitable temperatures (*1,14–16*).

**Figure 2.**
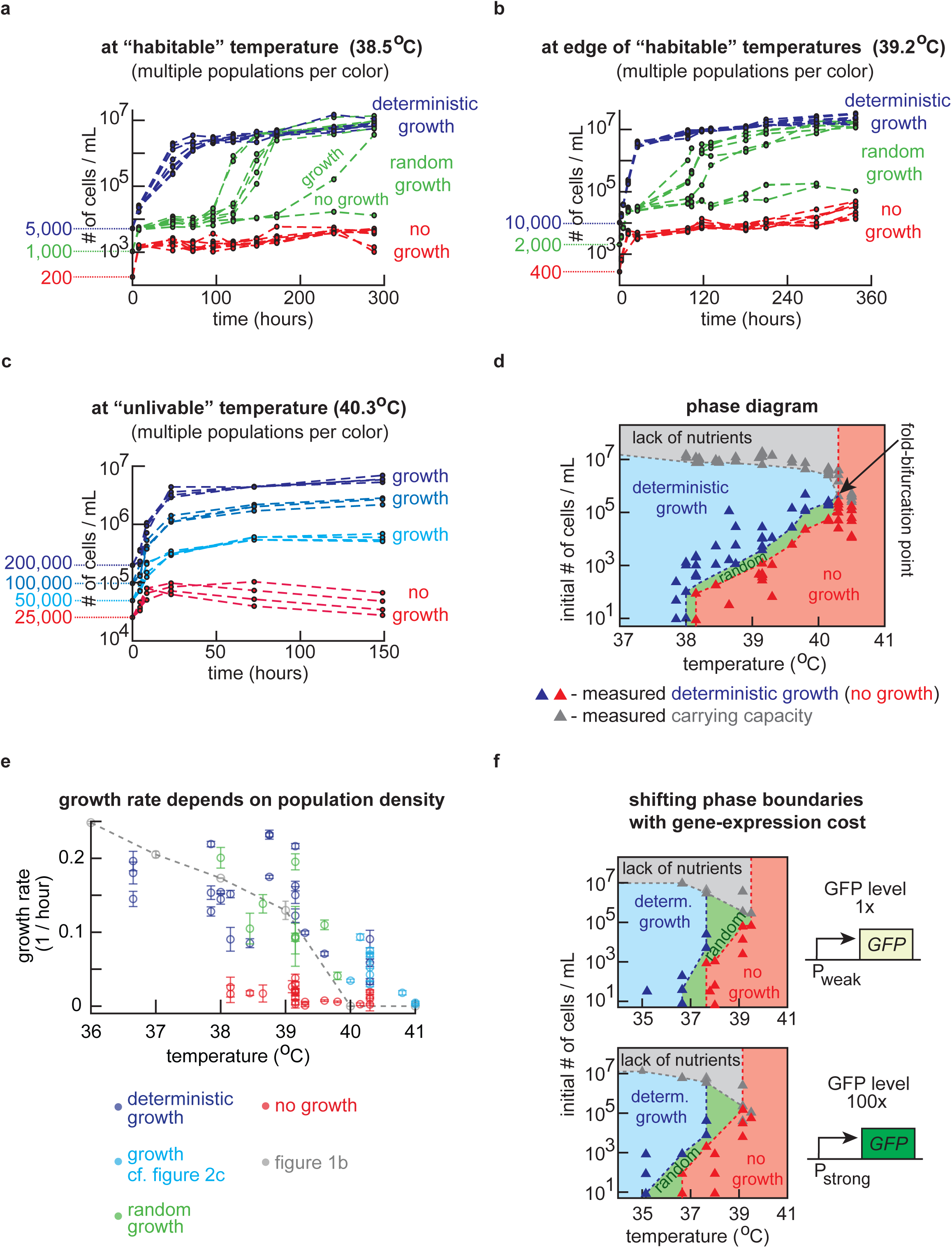
Population density determines replicability of cells and habitability of each temperature. (**a-c**) Population density (number of cells / mL) measured over time with a flow cytometer for populations of wild-type yeast of differing initial population-densities at (**a**) a conventionally-defined habitable temperature (38.4 °C), (**b**) near the edge of conventionally-defined habitable and unlivable temperatures (39.2 °C), (**c**) and at a conventionally-defined unlivable temperature (40.3 °C). Figure 1b sets the conventional definition of temperature’s habitability. For (a-b): Each color shows eight populations that start with the same density. Red curves show no growths beyond initial transient growths (i.e., “no growth”). Green curves show unpredictable growths (i.e., “random growth”). Blue curves show deterministic, exponential growths whereby all populations identically grow (i.e., “deterministic growth”). For (c): Each color shows four populations with the same initial population-density. All colors except the red show growths by ∼10-fold. (**d**) Phase diagram constructed from measurements. Colors represent the behaviors mentioned in (b) - blue region marks deterministic growth, green region marks random-growth, red region marks no-growth, and grey region marks populations not growing as they have more cells than the carrying capacity. Each triangle represents an experiment of the type shown in (a-c), performed at a specific initial population-density and temperature (Supplementary Figs. 3-4). A triangle’s color represents the phase exhibited by populations. See caption of Figure S4 for a detailed description of how we deduced the phase boundaries from the measurements. (**e**) Growth rates of populations in the no-growth phase (red), random-growth phase (green) and deterministic-growth phase (blue) as a function of temperature (error bars are s.e.m., *n* = 6 or more for temperatures below 40 °C, and *n* = 3 for temperatures above 40 °C). Grey data are from Fig. 1b. (**f**) Phase diagrams constructed for engineered yeast strains that constitutively express *GFP* at the indicated levels (1x and 100x - see Supplementary Fig. 5). Triangles indicate experimental data (samples of growth data represented by these triangles are in Supplementary Fig. 5). Since the wild-type strain’s genetic background is slightly different from that of GFP-expressing strains, compare the 1x-GFP strain with the 100x-GFP strain only.

We also observed the same three population-level behaviors - no growth, random growth, and deterministic growth - as a function of the initial population-density for a temperature (∼39 °C) that is near the boundary of the “habitable” and “unlivable” regimes (Fig. 2b). Moreover, at a temperature that is supposedly unlivable (∼40 °C), according to the conventional picture (Fig. 1c), we found that a population with an initial density of at least 50,000 cells/mL did, in fact, grow by ∼10-fold and then stably remained at a high density for several days without reaching the carrying capacity (Fig. 2c - non-red curves). In contrast, if the initial population-density was just half of this value (25,000 cells/mL), then the population density, instead of plateauing, continuously decreased over several days after an initial, transient growth (Fig. 2c - red curves). These results show that just a two-fold difference in the initial population density can determine whether there is a net cell-death or net cell-growth at ∼40 °C, which the conventional picture of autonomous cell-replications cannot explain.

### Phase diagram shows allowed population-level behaviors across temperatures

Above results (Figs. 2a-c) show that in order to determine whether a population grows or not, one must know *both* the temperature and the initial population-density. We can summarize this in a “phase diagram” that we constructed by performing the above growth experiments at multiple, additional temperatures with multiple, initial population densities (Fig. 2d and Supplementary Figs. 3-4). The phase diagram consists of four phases - deterministic growth, random growth, no growth, and no-growth due to insufficient nutrients (i.e., the initial population-density is beyond the carrying capacity for a given temperature) - as a function of the initial population density and temperature. It shows that the conventional picture (Figs. 1b-c) mistakenly arises because one typically ignores the initial population-density or sets it to be within some narrow range when studying population growth across temperatures. This leads to, for example, the growth rate *appearing* to decrease as the temperature increases within a given range (e.g., 36.5 °C ∼ 39 °C) (Fig. 1b). But in fact, for the same temperature range, we found that the populations’ growth rates - when they grew in the deterministic-growth and random-growth phases - were poorly correlated with temperature and could highly vary among populations even for the same temperature if we widely varied the initial population-density (Fig. 2e). In constructing the phase diagram, we determined at least how many cells per volume were necessary to guarantee that a population grew at each temperature (i.e., minimum number of cells per unit volume required for a deterministic growth at each temperature). A curve that represents how this value varies as a function of temperature forms the boundary between the deterministic-growth and random-growth phases in the phase diagram (Fig. 2d). We also determined at most how many cells per volume were necessary to guarantee that a population never grew at each temperature. A curve that represents how this value varies as a function of temperature forms the boundary between the random-growth and no-growth phases in the phase diagram (Fig. 2d). We found that both of these values - the minimum starting population-density required to guarantee that a population grows at a particular temperature and the maximum starting population-density required to guarantee that a population does not grow at a particular temperature - are extremely sensitive (i.e., “ultra-sensitive” (*20, 21*)) to temperature. For example, the phase diagram revealed that each of these two values change by ∼100-fold as the temperature increases from 39 °C to 40 °C (Fig. 2d). The random-growth phase, in the phase diagram, lies between the no-growth and deterministic-growth phases - one might see it as a hybrid of the growth and no-growth phases - and is thin along the axis that represents the initial population-density. Its thinness reflects our previous observation that a small change (5-fold or less) in the initial population-density can transform either a no-growth or a deterministic growth into a random growth (Fig. 2a) (i.e., populations are ultra-sensitive to their initial densities). Intriguingly, the phase boundaries - the borders between the four different phases - all converge at a single point, a “fold-bifurcation point”, whose coordinate is (40.3 °C, 1 x 10^5^ cells/mL) in the phase diagram (Fig. 2d). This convergence of all four phases at the fold-bifurcation point leaves only the no-growth phase for temperatures higher than 40.3 °C. Thus, according to the phase diagram, a population cannot grow to reach the carrying capacity regardless of its initial density for temperatures higher than 40.3 °C. More precisely, at temperatures higher than 40.3 °C, a population can grow but would eventually stop growing before reaching the carrying capacity, with the final population-density depending on the initial population-density. The term, “fold-bifurcation point”, comes from dynamical systems theory and is appropriate here because, according to this theory, this is the point in the phase diagram where a stable fixed point (carrying capacity) - this point “attracts” growing populations - merges with an unstable fixed point (the upper boundary of the no-growth phase) - this point “repels” populations towards extinction or the carrying capacity. The fold-bifurcation point is special for another reason which we will later turn to - one that is reminiscent of critical points in phase diagrams of physical systems in which temperature is one of the main parameters.

### Tuning the cost of expressing a single superfluous gene reshapes habitability of temperature

To explore possible ways of manipulating the phase diagram (i.e., where the phase boundaries are), we genetically engineered the wild-type yeast so that it constitutively expressed the Green Fluorescent Protein (GFP), which serves no function for cell growth. We constructed two such strains - one expressing GFP at a relatively low level (defined as 1x) and another at a relatively high level (∼100x) (Supplementary Fig. 5). We repeated the growth experiments with these two strains, for multiple temperatures and initial population-densities, and used these results to construct a phase diagram for each strain (Fig. 2f and Supplementary Fig. 5). Comparing the phase diagram of the 1x-GFP-expressing strain with that of the 100x-GFP-expressing strain revealed that increasing the GFP expression shifts the phase boundaries towards lower temperatures in such a way that a population of 100x-GFP-expressing strain need to have a higher initial population-density, compared to the 1x-GFP-expressing strain, in order to grow at a given temperature. Specifically, this means that the random-growth and no-growth behaviors are possible for the 100x-GFP-expressing strain at lower temperatures than for the 1x-GFP-expressing strain (Fig. 2f). For example, at ∼36 °C, a population with high GFP-levels (100x) can be in the random-growth and no-growth phases whereas the yeasts with the low GFP-levels (1x) can only deterministically grow regardless of how few cells there are. Therefore, a population that could grow at a given temperature can no longer grow at that same temperature because its cells express GFP. One needs to either increase the initial population-density or decrease the temperature to observe its growth. These results show that the cost of expressing superfluous genes can markedly alter the phase boundaries’ locations and shapes. In particular, this means that reducing the cost of expressing a single, unnecessary gene can increase the life-permitting temperature by several Celsius degrees. In light of previous studies (*22,23*), this may be due to the fact that overexpressing a superfluous gene by a sufficiently large amount shifts the intracellular resources away from performing roles that are for cell growth.

### Single-cell measurements reveal that a few “pioneer” cells initiate replications in randomly growing populations and sustenance of transiently replicating sub-populations in non-growing populations

To gain further insights, we turned to single-cell-level measurements. Unlike the GFP-expressing strains, the wild-type strain lacks a functional *ADE2* gene for synthesizing adenine. Since we incubated yeasts in the minimal media with all the essential amino acids and nitrogenous sources - including adenine that represses their adenine-biosynthesis - the wild-type cells were still capable of growing. But, as is well-known, having a defective *ADE2* gene turns yeasts red if they have not divided for some time because they have accumulated red pigments - these are by-products of the not-fully-repressed and defective adenine-biosynthesis (*24*). The cells can only dilute away the red pigments through cell divisions. Defective *ADE2* gene cannot be a reason for any of the population-density-dependent growth behaviors that we observed because the GFP-expressing strains do have the functional *ADE2* gene and yet exhibit the same, surprising behaviors as the wild-type cells (Fig. 2f). But the red pigments were useful because, for the wild-type populations, we could use our flow cytometer’s red-fluorescence detector to count how many non-replicators (red cells) and how many replicators (non-red, “white cells”) co-existed in a population at each time point (Supplementary Fig. 6). As a result, we discovered that, in every population and at every temperature, nearly all cells were red at the start of each growth experiment due to the cells having just been transferred from cultures that grew to saturation in 30 °C to the higher temperature. Subsequently, the number of replicators increased to about 10-30% of the population during the initial, transient growth in which the cells from 30 °C adjust to the higher temperature (Supplementary Fig. 6). Afterwards, the number of replicators varied depending on which phase the population has. For a deterministically growing population, the number of replicators kept increasing over time until it reached the carrying capacity (Supplementary Fig. 6) whereas it typically decreased until very few cells (∼1-5 % of population) remained as replicators for random-growth and no-growth populations (Supplementary Fig. 6). Subsequently, two behaviors were possible depending on whether the population was in the no-growth or random-growth phase. For populations in the random-growth phase, the number of replicators, after unpredictable hours or days, suddenly started to increase by orders of magnitude until the population reached the carrying capacity (Supplementary Fig. 6). On the other hand, for populations in the no-growth phase, the number of replicators sustainably remained low (∼1-5% of the population) and fluctuated up and down by a few-fold during the incubation (∼300 hours in our experiments) (Supplementary Fig. 6). These fluctuations were too small to noticeably change the total population density over time. Yet, this result revealed that there is always a small sub-population of transiently replicating cells and that these transient replications are such that the fraction of replicators in the population could stably remain in low numbers (e.g., ∼1% of total population). We will later return to these single-cell data when we introduce a mathematical model that recapitulates them.

### Cells collectively delay and prevent population extinctions at high temperatures

We next asked whether cell deaths, like cell replication, also depend on the initial population density. At several temperatures, we measured how the number of surviving cells changed over time for a population that we kept in the no-growth phase (i.e., population with not enough cells to trigger its own random or deterministic growth). We counted the number of survivors by taking out an aliquot of cells from the population at different incubation times, spreading it onto an agar pad at 30 °C, and then counting how many colonies formed (Supplementary Fig. 7). Surprisingly, these measurements deviated qualitatively - not just quantitatively - from the conventional theory of cell deaths which states that deaths of heat-shocked cells are autonomous. Specifically, the conventional theory says that the number of survivors should exponentially decrease over time with a constant rate until the population becomes extinct due to every cell having the same, fixed probability of dying per unit time regardless of the population density (*4*) (Fig. 3a - brown line). Yet, we discovered that the rate at which cells die continuously decreases over time. This leads to the number of survivors decreasing as a heavy-tailed (power-law-like) function (Fig. 3a - blue curve), instead of as an exponential decay. Consequently, the number of survivors appears to decrease exponentially at a constant rate during the first day of incubation but then, after a few days, decreases exceptionally slowly as a heavy-tailed decay. The heavy-tailed decay means that the population continuously decelerates and then eventually ceases its approach to extinction. This causes the number of survivors to plateau after some time. For example, the number of survivors at 41 °C did not noticeably decrease after three days of incubation and deviated by ∼10^7^-fold from the number of survivors that the theory states one should have (Fig. 3a - last time point; Supplementary Figs. 7-8). Moreover, we discovered that the rate at which cells die during incubation at a high temperature depends on the initial population-density (Fig. 3b & Supplementary Fig. 7). After a transient growth of about one day of incubation at a high temperature - this arises from the cells being transferred from 30 °C to the desired high temperature - the number of survivors seems to exponentially decrease over time (during the first day of incubation at a high temperature) before it noticeably enters a heavy-tailed decay regime on later days (Fig. 3b). Hence, we can assign a constant rate of decay to each population in order to describe how the number of survivors initially decreases (e.g., during the first day of incubation at a high temperature). We found that this rate - which we will call the “initial death-rate” - depends on the initial population-density. Namely, we discovered that as the initial population-density increases, the initial death-rate decreases, meaning that number of survivors decreases more slowly during the first day for higher initial population-densities (Fig. 3b - three dashed lines with differing slopes). For example, after one day of incubation at 41.0 °C, a population that started with ∼92,000 cells/mL had about 100-fold less survivors than a population that started with about three times more cells (Figure 3b - compare blue and purple lines). Yet, the prevalent theory states that it should be 3-fold less, not 100-fold less. Taken together, these results establish that a population that starts with a higher density has a larger fraction of its cells remaining as survivors after one day and, due to the heavy-tailed decay, has vastly more survivors than a population that started with a lower density (Fig. 3b - last, faded data points for each color). This population-density dependent effect - a small initial difference in population density amplifying to a large, nonlinear change in the final population density - suggests a highly nonlinear cooperative effect that cells have on each other’s survival.

**Figure 3.**
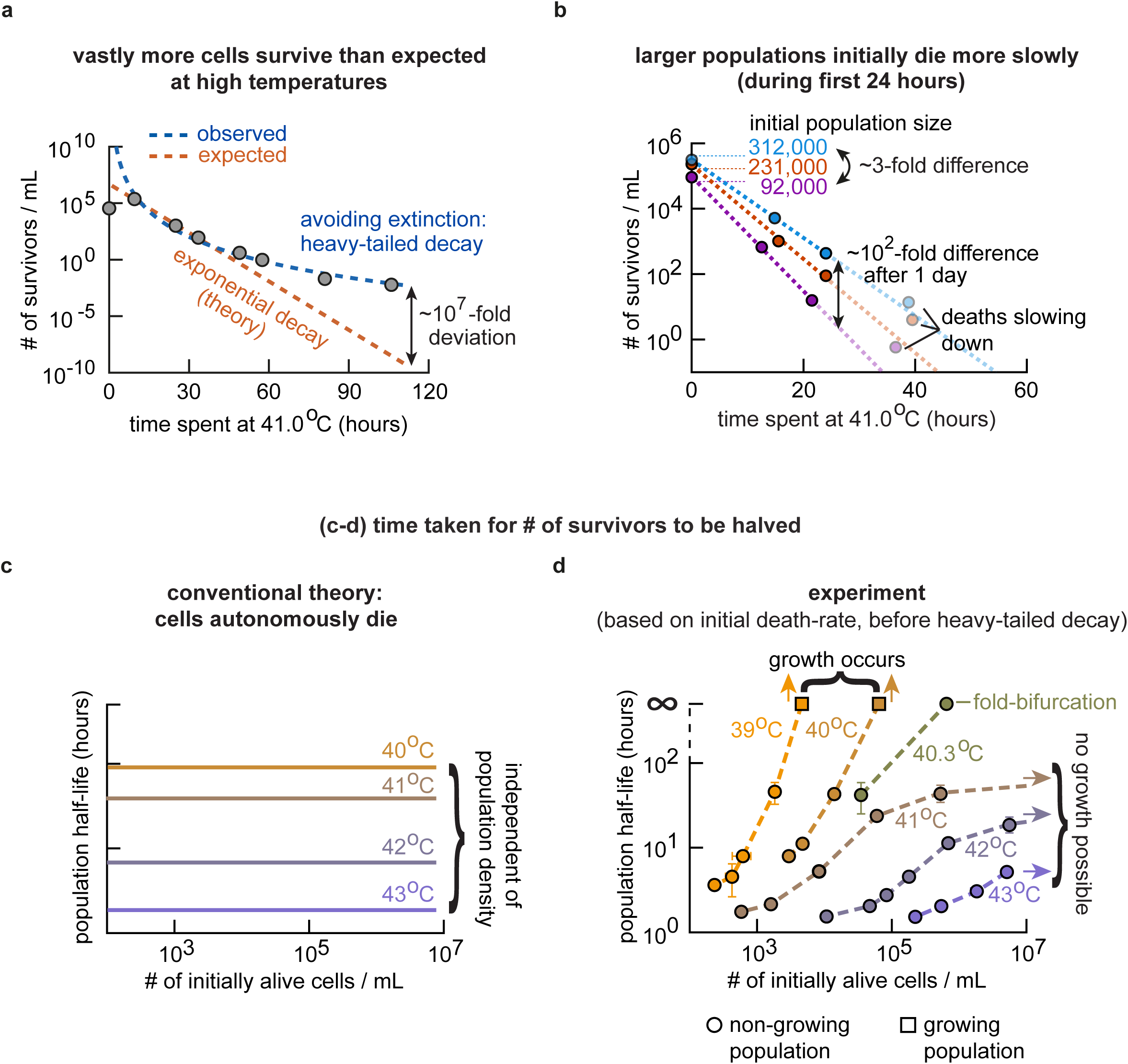
Cell do not autonomously die and their deaths depend on initial population-density at high temperatures: populations of higher initial densities decelerate and halt their approach to extinction more rapidly. (**a**) We determined the number of survivors per mL over time in a non-growing, no-growth-phase population at 41.0 °C by transferring an aliquot of the liquid culture to an agar pad at different time points, incubating the agar pad at 30 °C, and then counting the number of colony forming units (“# of survivors/mL”) (see Supplementary Fig. 7). Circles represent measured numbers of survivors per mL. Brown line is an exponentially decaying function fitted to the first three data points (between 10 and 40 hours). Blue curve is a power-law function fitted to all the data points. Also see Supplementary Fig. 7. (**b**) Number of survivors per mL for three populations of differing initial population-densities – all of them in the no-growth phase at 41.0 °C - measured as in (a). Initial population-densities, after transient growths, are 92,000 cells / mL (purple), 231,000 cells / mL (orange), and 312,000 cells / mL (blue). For each color, a dashed line is an exponentially decreasing function that we fitted to the first three timepoints (i.e., deaths that occur during the first day of incubation after a transient growth). Also see Supplementary Fig. 7. (**c**) Cartoon illustrating the conventional view which states that cells autonomously die and that every cell has the same probability of dying per unit time. This means that the population half-life (i.e., time taken for a population density to be halved) is independent of the initial population-density for every temperature. Different colors represent different temperatures as indicated. (**d**) Population half-life, plotted here as a function of initial population-density, is defined as the time taken for a population density to be halved, based on fitting an exponentially decreasing function to the number of survivors / mL measured during the first 24 hours of incubation, which in turn occurs after a ∼20-hours of transient growth that is due to the cells coming from 30 °C and adjusting to the new temperature. Hence, we measured the number of survivors at approximately 20 hrs, 28 hrs, 36 hrs, and 44 hrs after we incubated the populations at the desired high temperature (these times are measured after the ∼20 hours of transient growths ended). Usually, the 44-hr data point already deviated from exponential decay and thus we usually omitted them from the exponential fits. Shown here are the half-lives of populations at 39.2 °C, 40 °C, 40.3 °C, 40.8 °C, 42 °C, and 43 °C, each in a different color as indicated. Error bars: s.e.m. (*n* = 3 for all data points). Circles represent populations in the no-growth phase. The two squares (at 39.2 °C and 40 °C) represent populations that grew due to having sufficient densities to trigger their own growths, in accordance with the phase diagram (Fig. 2d).

### A special temperature that defines the fold-bifurcation point separates two extinction-avoidance regimes

The conventional theory of cell death - which imposes a constant, exponential death rate - says that the time taken for the number of survivors to be halved decreases with increasing temperature and that it is independent of the initial population density (Fig. 3c). Yet, our experiments revealed a starkly different story (Fig. 3d). We can see this by extracting the initial half-life, which we define to be the amount of time taken for the number of survivors to be halved based on the initial death-rate (Fig. 3d). The initial half-life is inversely proportional to the initial death-rate. We discovered that while increasing the initial population density always increases the population’s initial half-life, it is the temperature that determines how sensitively the initial half-life depends on the initial population-density. In particular, we found that a population’s initial half-life has two regimes of sensitivities, depending on whether the temperature is below or above 40.3 °C. Intriguingly, 40.3 °C is the temperature where the fold-bifurcation point lies in the phase diagram (Fig. 2d). For temperatures below 40.3 °C, we found that the initial half-life can increase from a few hours to a few days when the initial population density changes by just a few fold (e.g., 3-fold) (Fig. 3d - yellow and brown curves for 39 °C and 40 °C respectively). Moreover, as we keep increasing the initial population density, the initial half-life keeps increasing and eventually reaches infinity, due to the fact that a sufficiently high-density population would grow at these temperatures (Fig. 2d). For temperatures above 40.3 °C, we discovered an opposite trend: increasing the initial population-density above some value hardly changes the initial half-life, leading to the half-life eventually plateauing at a finite value as we keep increasing the initial population density (Figure 3d - purple curves for 41 °C ∼ 43 °C). Thus, after some amount, increasing the initial population density does not yield much gain in the initial half-life. This occurs because a population can never grow regardless of its density at these temperatures (Fig. 2d). Exactly at 40.3 °C, we found that a population whose initial density matches that of the fold-bifurcation point (∼1 x 10^5^ cells/mL) neither replicates nor dies. It is unable to grow yet can indefinitely maintain its number of survivors at a nearly constant value (i.e., its initial half-life is infinite because its initial death-rate is zero). The fold-bifurcation point is the only combination of temperature (40.3 °C) and population density (∼1 x 10^5^ cells/mL) for which an infinite initial half-life is possible without the population growing. This makes the fold-bifurcation point special: a population at the fold-bifurcation point initially can - on average and subject to population-density fluctuations - maintain its cell numbers at a constant value by having the number of births balanced by the number of deaths per unit time. Hence, a population at the fold-bifurcation point can live for a very long time.

Taken together, the conventional theory of autonomous cell deaths (*4*) cannot explain the features of cell deaths that we uncovered (Figures 3a-d). Likewise, mechanisms that are similar to those known for yielding long-lived populations under other stresses - such as persistence to antibiotics (*25*) - also cannot explain our data on cell deaths (Supplementary Fig. 8). There are two reasons for their inability to do so. First, how fast cells die over time depends on the initial population-density at high temperatures (rather than being independent of the initial population-density as in the case of antibiotic persistence). Secondly, the rate at which cells die changes over time at high temperatures (rather than remaining constant over time as in the conventional theory of autonomous cell deaths). Thus, having “persistor-like” cells as a subpopulation that is more tolerant of a high temperature than the rest of the population is inconsistent with above two features in our data (see Fig. S8 for more in-depth explanations). Moreover, having more heat-tolerant mutants forming a subpopulation - which may have formed during or before the first few days of incubation at a high temperature – also cannot explain our data on cell deaths (Figs. 3a-d and Supplementary Fig. 8). We verified this by taking the last remaining survivors of a high temperature environment (i.e., cells at the tail-end of the heavy-tailed decay), growing them to a high density at 30 °C, and then subjecting the resulting population to the same, original high temperature with the same initial population-density as their ancestral (the first) population. We observed that this population approached extinction over time by following the same heavy-tailed decay function as its ancestral population (Supplementary Fig. 8), which is inconsistent with having more heat-tolerant mutants forming a subpopulation before or during a heat-shock – such a subpopulation should lose viable cells over time as a slow, exponentially decaying curve or not decay at all at the high temperature rather than as the same heavy-tailed function as the ancestral population (Supplementary Fig. 8).

### Extracellular factor dictates cell replications at high temperatures

Having established that replications and deaths of cells depend on the initial population-density, we sought to uncover the underlying mechanisms. We first focused on finding a mechanism that is responsible for cell replication. As we will show, this mechanism also explains the cell deaths. Broadly, we can consider two classes of mechanisms. One is that the “factors” that dictate a cell’s replication, at high temperatures, resides purely within that cell (i.e., purely intracellular factors dictate cell replications). In this case, cell replication would be a purely cell-autonomous process and the reason that it depends on the initial population-density (Figs. 2a-c) would be that if we have more cells, then it is more likely that at least one cell would manage to replicate. The other possibility is that an extracellular factor dictates cell replications. In this case, a cell’s replication would depend on elements outside of that cell which may include the other cells in the population. To distinguish these two classes of mechanisms, we performed experiments in which we physically separated the cells from their extracellular environment. Specifically, in one experiment, we took cells that were exponentially growing at a particular temperature and then, at that same temperature, transferred them to a fresh medium that never harboured any cells before. This experiment would reveal whether the cells can keep on growing which, if true, would mean that purely intracellular factors dictate cell replications (Fig. 4a - bottom left tube). In another experiment, we took away the growth medium from cells that were growing at a particular temperature and then, at that same temperature, transplanted into the medium fresh, non-growing cells whose initial population-density was too low for growth according to the phase diagram. This experiment would reveal whether the transferred medium, taken from growing cells, can induce growth of a population that should not grow which, if true, would mean that extracellular factors dictate cell replications (Fig. 4a - bottom right tube). As an example of an experiment in which we transferred the growing cells instead of their growth medium, we took some wild-type cells from a population that was exponentially growing at a particular temperature (∼39 °C; Fig. 4b - blue curves) and then transplanted them to a fresh medium at that same temperature so that this newly created population started with a population density (400 cells/mL) that should not permit growth according to the phase diagram (Fig. 2d). We found that these cells stopped growing in the fresh medium soon after the transfer (Fig. 4b - green curves; also see Supplementary Fig. 9). This strongly suggests that extracellular factors dictate cell replications (i.e., a cell does not have an intrinsic ability to replicate that is independent of its extracellular environment).

**Figure 4.**
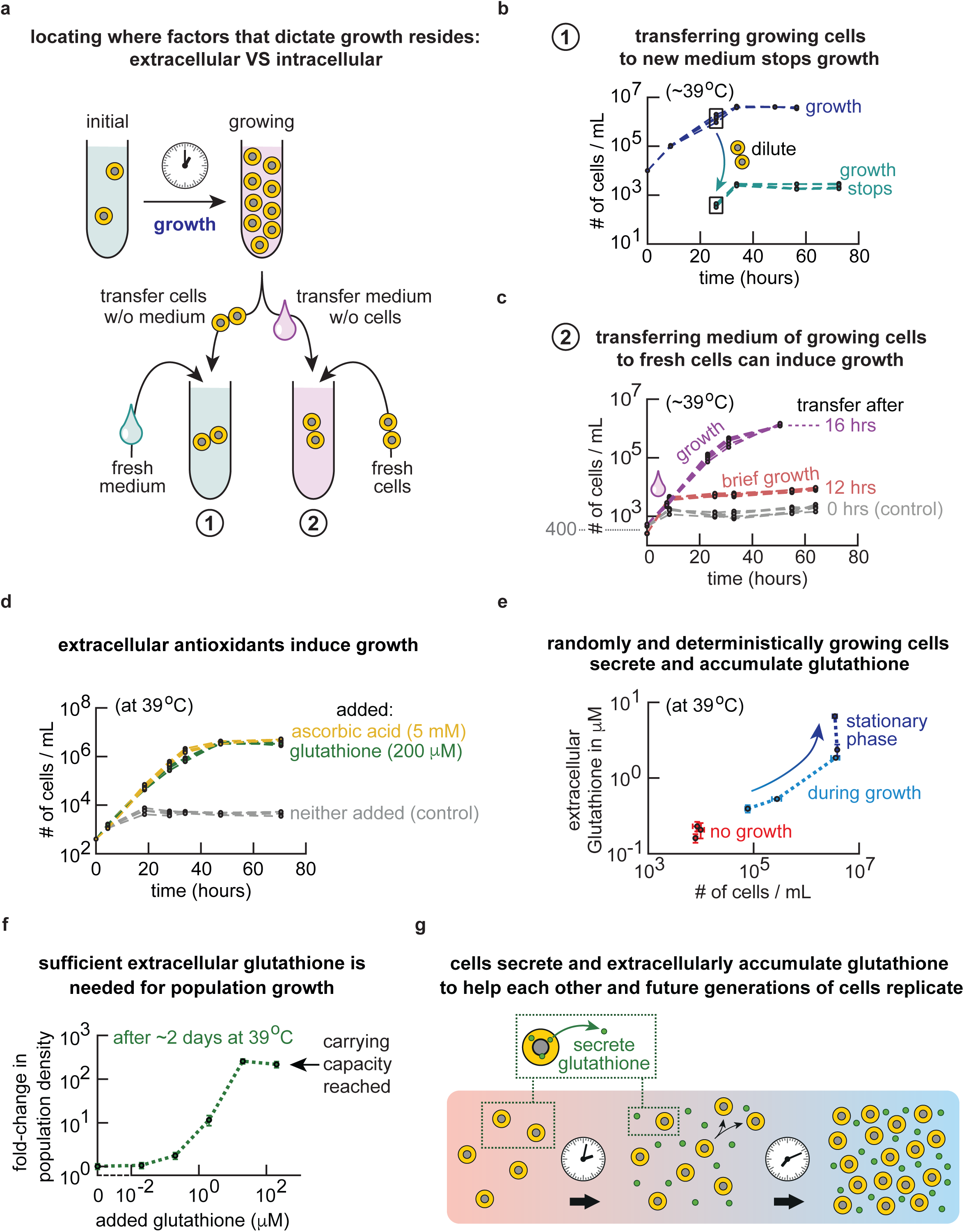
Cells secrete and extracellularly accumulate glutathione to help each other and future generations of cells replicate at high temperatures. (**a**) Schematic description of experiments in (b) and (c) to distinguish two classes of mechanisms. Labelled “1’: transfer growing cells into a fresh medium (to determine if intracellular factors dictate population growth). Labelled “2”: take out only the liquid medium of a growing population and then incubate fresh cells in it (to determine if extracellular factors dictate population growth). (**b**) Experiment labelled “1” in (a) at 39.2 °C. Blue curves show populations of wild-type yeasts (initially ∼10.000 cells/mL) that start to exponentially grow. Cells of these populations in exponential growth, marked by the boxed blue data point, were transferred without their growth medium into a fresh medium. This resulted in ∼400 cells/mL in the new culture, represented by the green curves (four replicate populations). Also see Supplementary Fig. 9. (**c**) Experiment labelled “2” in (a) at 39.2 °C. Wild-type population’s density over time for cells that were incubated in a growth medium that previously harbored a population of exponentially growing wild-type cells at ∼39 °C for 0 hours (grey curves), 12 hours (red curves), and 16 hours (purple curves). Each color shows at least 6 replicate populations. Also see Supplementary Fig. 10. (**d**) At 39.2 °C. All wild-type populations start at 400 cells/mL, which is too low of a density to permit growths. Adding either ascorbic acid (5 mM - yellow) or glutathione (200 μM - green) to the growth media induces population growths. Without adding either one, populations do not grow (grey). Each color shows three replicate populations. (**e**) At 39.2 °C. Shown here are the concentrations of extracellular glutathione for wild-type populations in the no-growth phase (initially 400 cells/mL - red data), random-growth phase (initially 2,000 cells/mL), and deterministic-growth phase (initially 10,000 cells/mL). Three biological replicates are shown for each phase (*n* = 3; error bars are s.e.m.). We measured three timepoints for each phase. Plotted here are the concentrations of extracellular glutathione as a function of the population density over time. The arrow shows both the population density and concentration of extracellular glutathione increasing together over time (see also Supplementary Fig. 13). (**f**) Sensitivity of no-growth wild-type populations (initially ∼400 cells/mL) at 39.2 °C to glutathione that we added into the growth medium. Shown here is, as a function of glutathione concentration, the fold-change in the population density after two days of incubation with the added glutathione. Last two data points show ∼200-fold change, which represents the populations having fully grown to reach the carrying capacity (see also Supplementary Fig. 13). (**g**) Cartoon illustrating the mechanism deduced in (a-f). Yeasts secrete glutathione at high temperatures. The extracellular glutathione thus accumulates. When its concentration reaches at least a threshold amount (∼0.3 μM from (f)), the population replicates at high temperatures by noticeable amounts.

To further test the idea that extracellular factors dictate cell replications, we transferred the growth medium from cells that were growing at a particular temperature (∼39 °C) to fresh, non-growing cells and tested whether these cells could also grow at that same temperature. Strikingly, we found that these fresh cells could grow after being transplanted into the medium (Fig. 4c - purple curves), even though they initially had a low population-density (400 cells/mL) that - without the transplanted medium - prohibits growth according to the phase diagram. Crucially, whether the transferred medium induced growth or not depended, in a highly sensitive manner, on the amount of time that the medium harboured exponentially growing cells before the transfer (Fig. 4c and Supplementary Fig. 10). For example, at ∼39 °C, if cells exponentially grew for 12 hours or less before we transferred their growth medium to fresh cells, then the fresh cells did not grow in the transferred medium at that same temperature (Fig. 4c - grey and red curves and Supplementary Fig. 10). However, if cells had grown for 16 hours or more before we transferred their growth medium to fresh cells, then the fresh cells grew in the transferred medium (Fig. 4c - purple curves and Supplementary Fig. 10). These results support the following idea: for a population to grow by an appreciable amount at high temperatures, the concentration(s) of growth-inducing extracellular factor(s) must be above some threshold value(s). The next question then is whether these extracellular factors must be secreted or depleted by cells – above or below some threshold value(s) respectively - to induce population growths.

To test whether it is depletions of some extracellular factors that induce population growths, we performed several experiments (Supplementary Figs. 10-11). In one experiment, we took a fresh growth media that contained all the essential amino acids, diluted them with water by different amounts, added a saturating level (2%) of glucose to them, and then incubated cells in them at high temperatures (Supplementary Fig. 11). We found that a population could grow in any of these diluted media only if their initial population-densities matched the values that, according to the phase diagram (Fig. 2d), permit growth. The same was true if we decreased only the glucose in fresh media by various amounts - without diluting other factors such as amino acids - and then incubated cells in them (Supplementary Fig. 11). Thus, population growths are not dictated by depletions of any of the resources that were already in the media before cells were incubated in them. Instead, these results strongly suggest that a secretion of factor(s) induces population growths.

As a complementary experiment, we took growth media that were harbouring growing cells at 30 °C for various amounts of time, from a few hours to 12 hours. We then transferred these media, after flowing them through a filter membrane to eliminate any cells from them, to a fresh cell-population and then incubating the resulting liquid culture a high temperature (39 °C). These newly created populations all had the same initial population-density that were too low for growth at 39 °C on their own (i.e., 400 cells/mL) (Supplementary Fig. 10). We found that the transferred media taken from the log-phase cultures at 30 °C did not cause the fresh cell-population to grow at 39 °C. This result further supports the idea that a secretion, not depletions, of factor(s) induces population growths. Intriguingly, when we took a medium from a population that was in a stationary phase at 30 °C – this population underwent a diauxic shift after a log-phase growth – and then transplanted a fresh cell-population into at 39 °C, we found that this population grew at 39 °C (Supplementary Fig. 10). Hence certain factor(s) that yeasts secrete at 30 °C when they are in a stationary phase (following a diauxic shift) can induce population growths at high temperatures. We reasoned that these secreted factor(s) may also be the same factor(s) that cells growing in log-phase at high temperatures also secrete to induce population growths.

### Extracellular antioxidants - glutathione and ascorbic acid - enable yeasts to replicate at high temperatures

To help us identify the secreted factor(s) responsible for cell replications at high temperatures, we performed a transcriptome analysis (RNA-seq) on wild-type yeast at different parts of its phase diagram (Supplementary Fig. 12). We found that deterministically growing cells at high temperatures, compared to those growing at 30 °C, downregulate genes involved in the central carbon metabolism and majority of other genes in general. But we also found that they had, compared to those growing at 30 °C, upregulate genes that are associated with DNA damage response, translation initiation, and biogenesis and assembly of cell membranes. These changes in gene expressions are similar to those seen for yeasts experiencing environmental stresses in general (*26,27*). We hypothesized that a cell upregulates these genes to repair damages to various cellular components. Specifically, we hypothesized that reactive oxygen species may cause these damages at high temperatures. Reactive oxygen species are known to damage nucleic acids (*28*), proteins (*29*), and lipids in the cell membrane (*30*). Indeed, our transcriptome analysis revealed that many genes associated with the mitochondria were downregulated in replicating yeasts at high temperatures compared to yeasts at 30 °C. Since respiration through mitochondria creates reactive oxygen species, we reasoned that downregulating respiratory mitochondrial genes may decrease the amount of reactive oxygen species that form and, as a result, allow the yeast to replicate at high temperatures (*31,32*). Cells can combat reactive oxygen species by producing antioxidants. Antioxidants reduce oxidative-stress-related damages by capturing and then deactivating reactive oxygen species through redox reactions. We hypothesized that budding yeasts may be secrete antioxidants at high temperatures. Moreover, we reasoned that – as suggested by our media-transfer experiments (Fig. 4c and Supplementary Fig. 10) – the secreted antioxidants must be present in high enough concentrations, meaning that a population density must be high enough, in order for a population to grow at high temperatures. Strengthening this hypothesis is that heat shocks are known to cause budding yeasts to produce and maintain an intracellular – but not extracellular - pool of glutathione (*33*). As an antioxidant, glutathione is widely used by many species, including humans. It is budding yeast’s primary, intracellular antioxidant (*32,34*). Also strengthening our hypothesis is the fact that after shifting from a log-phase growth to a stationary phase at 30 °C, budding yeasts are known to produce glutathione – though most of the glutathione is known to be intracellularly kept and only low amounts of it have been detected in their extracellular medium (*35*). Of particular interest to us is the fact that in a few known instances, budding yeasts can secrete glutathione – they are known to do so as an extracellular defense against arsenite that harms them (*36*). But for the budding yeast, glutathione’s extracellular role still remains underexplored and it is mainly viewed as an intracellular agent with varied roles – for example, as a regulator of iron metabolism (*37*) or an antioxidant (*38*). We reasoned that the very low concentration of extracellular glutathione, if it indeed exists in the growth medium of the stationary-phase population at 30 °C, may have induced the growth of the fresh cell-population at 39 °C that we previously observed (Supplementary Fig. 10). It is unclear, however, whether yeasts at high temperatures indeed secrete glutathione.

To first test whether antioxidants can even induce population growths at high temperatures, we added either glutathione or ascorbic acid - two prominent antioxidants (*34*) - or trehalose at various concentrations to the growth medium that harboured a low density of wild-type cells. Strikingly, we found that both glutathione and ascorbic acid can induce population growths as long as we add either one at high concentrations (e.g., glutathione at 200 μM and ascorbic acid at 5 mM) (Fig. 4d). Furthermore, we determined that trehalose cannot induce population growths at high temperatures (Supplementary Fig. 11). These results establish that at high temperatures, glutathione and ascorbic acid are sufficient for inducing growths of low-density populations that would not have grown on their own (e.g., 400 cells/mL, Fig. 2d). Moreover, these results suggest that reactive oxygen species causing damages is the primary reason that yeasts fail to replicate at high temperatures.

### Cells secrete glutathione to help each other replicate at high temperatures

Although ascorbic acid, known as “erythroascorbate” for the budding yeast, acts as an antioxidant in eukaryotes, its role in budding yeast remains unclear as researchers have not detected appreciable amounts of it in yeast (*33*). Therefore, we reasoned that glutathione is a more likely candidate as the secreted antioxidant. To test if yeast populations secrete glutathione at high temperatures, we measured the glutathione concentration over time in the growth medium for deterministically growing, randomly growing, and no-growth-phase populations at high temperatures (Fig. 4e and Supplementary Fig. 13). For populations in the no-growth phase at high temperatures (e.g., 39 °C), glutathione concentration barely increased during ∼50 hours of incubation. The population did not grow during this time. In contrast, for populations in the random-growth and the deterministic-growth phases at high temperatures, we detected a continuous increase in the extracellular glutathione concentration over the ∼50 hours of incubation, leading to ∼10-fold increases in the extracellular glutathione concentration. After these populations reached their carrying capacities and stopped growing, their extracellular glutathione concentration still continued to increase (Fig. 4e). These results establish that both log-phase and stationary-phase yeasts at high temperatures secrete glutathione. On the other hand, at multiple temperatures below 36.7 °C, we found that only stationary-phase yeasts after a diauxic shift, but not during a log-phase growth, secrete glutathione (Supplementary Fig. 13). Thus, in summary, budding yeasts secrete glutathione while in log-phase growth and stationary phase at high temperatures (above 36.7 °C) but they do not secrete glutathione during log-phase growth at lower temperatures. Consistent with glutathione being responsible for the population-density-dependent growths, the temperatures at which we detect secreted glutathione from log-phase growing yeasts – above ∼36 °C - coincide with the temperatures at which growth depends on the population density (Fig. 2d and Supplementary Fig. 4).

### Extracellular glutathione concentration must be above a threshold concentration to induce population growths

We next addressed how sensitive yeasts are to extracellular glutathione at high temperatures. At high temperatures (e.g., 39 °C), we added various concentrations of glutathione to growth media and then incubated a low density, no-growth-phase population in them. By measuring the fold-change in the population density after two days of incubation, we obtained a highly non-linear relationship between glutathione concentration and the fold-change in the resulting population density (Fig. 4f). Namely, when the extracellular glutathione concentration was just above 0.3 μM, the population grew by ∼10-fold. But when the extracellular glutathione concentration was either just below or much lower than 0.3 μM, the population hardly grew even after two days. Moreover, when the extracellular glutathione concentrations were much higher than 0.3 μM (e.g., ∼10 μM), populations deterministically grew and reached their carrying capacities (i.e., ∼200-fold growth). These results establish that a population must accumulate enough glutathione in the extracellular medium - above some threshold concentration (∼0.3 μM) - in order to grow (Fig. 4g). Consistent with glutathione being responsible for inducing population growths at high temperatures, this threshold concentration of glutathione matches the secreted and accumulated glutathione concentration that we measured in the growth media of populations that were able to grow at high temperatures (Fig. 4d).

### Minimal mathematical model recapitulates all the experimental observations

To test if glutathione secretion can tie together and quantitatively explain all our experimental data, we developed a mathematical model. In this model, cells secrete glutathione at a constant rate - this is the simplest possible scenario and we found that not assuming a constant secretion rate does not qualitatively alter the model’s outcomes (see Supplemental text). Moreover, aside from always secreting glutathione, a cell in the model takes one of three actions: replicate, die, or stay alive without replicating (Fig. 5a). A probability for each action determines what the cell does next (i.e., a cell “rolls” a three-sided, loaded dice to determine its next action). Specifically, we assume that the probability that a cell dies in the next time step to be fixed by temperature and thus remain constant over time (Fig. 5b - left panel). Moreover, we assume that the probability of dying to linearly increase with temperature. But we found that relaxing this assumption - having it non-linearly increase with temperature - does not change the model’s outcomes (see Supplementary text). We let the probability of a cell replicating in the next time step to non-linearly (sigmoidally) increase with glutathione concentration (Fig. 5b - right panel). This reflects our experimental observation that glutathione concentration must be above some threshold value to appreciably induce population growths (Fig. 4f). The probability of staying alive without dividing is then fixed by the probabilities of dying and of replicating. This model contains four parameters, three of which are rigidly fixed by (directly read-off from) the experimental data without any possibility of us adjusting their values: (1) the maximum growth rate that a population can have (∼0.25 / hour from our experiments – Fig. 2e), (2) the temperature at which heat-induced deaths begin to be non-negligible (i.e., temperature above which no-growth phase starts to exist - this is ∼38 °C according to the wild-type’s phase diagram (Fig. 2d) and it is also the temperature at which the red line in Figure 5b starts to increase above zero), and (3) the temperature at which heat-induced deaths are always dominant over cell replications (i.e., temperature above which only the no-growth phase exists - this is ∼40.3 °C according to the wild-type’s phase diagram (Fig. 2d) and it is also the temperature at which the highest possible probability of replicating matches the probability of dying (Fig. 5b - grey line)). The only free parameter that we can flexibly fit to our data (i.e., fit to combinations of data rather than directly reading-off from a single measurement) is the glutathione concentration at which the probability of replicating is half its maximum (Fig. 5b - blue curve).

**Figure 5.**
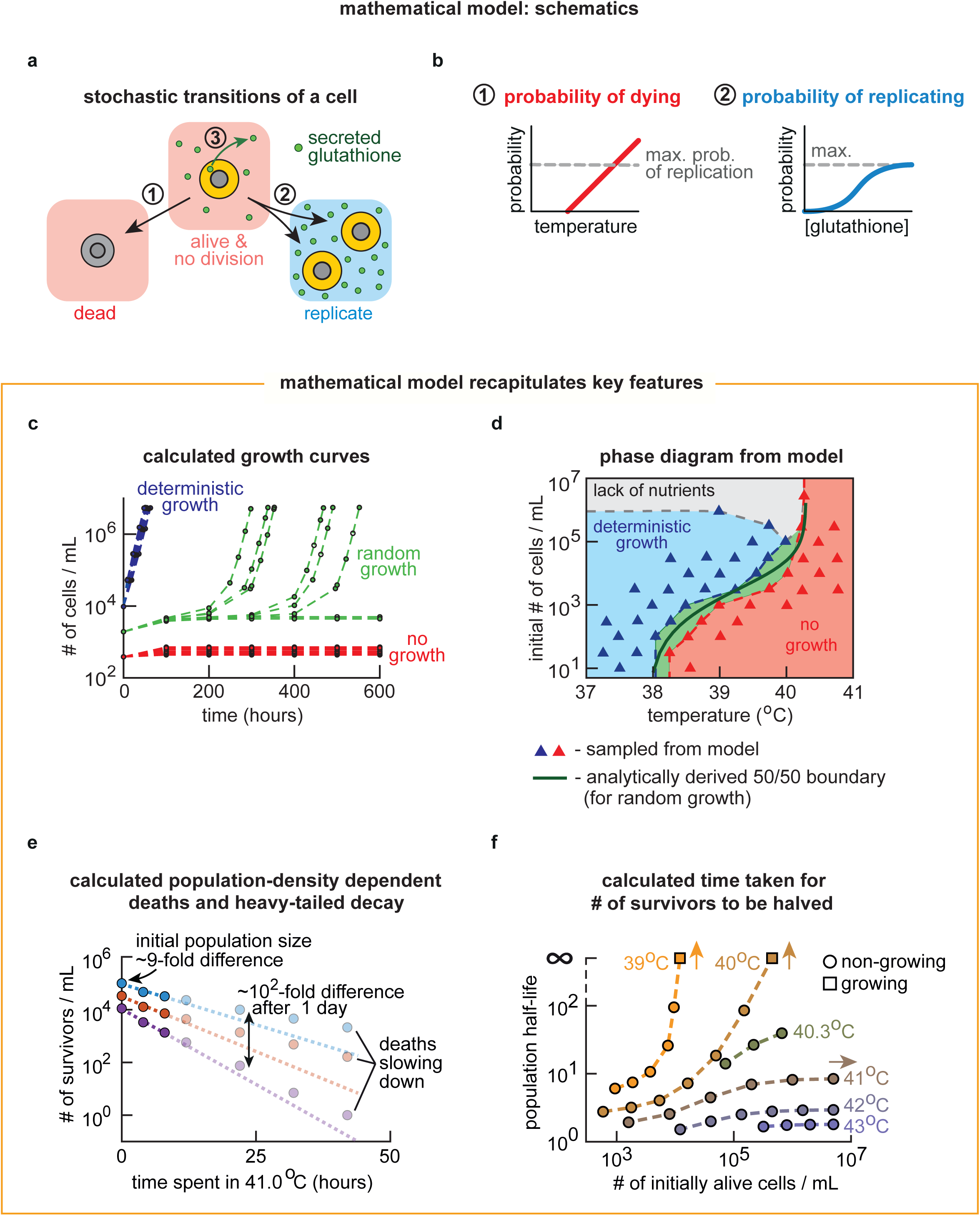
Mathematical model with one free parameter recapitulates all the main experimental data. (**a-b**) Description of the mathematical model (see full description in the Supplemental text). (**a**) A cell (yellow circle) can be in three states. In each time step, any alive cell either stays alive without replicating, replicates, or dies. Alive cells constantly secrete glutathione (green circle). (**b**) Schematic description of the probabilities that describe each of the transitions between states shown in (a). Left panel: probability of a cell dying (red line) is fixed by the temperature and does not change over time. It linearly increases with temperature and, beyond some temperature, it exceeds the maximum allowed value for the probability of a cell replicating (grey line). Right panel: probability of a cell replicating (blue curve) nonlinearly increases with the concentration of the extracellular glutathione. (**c-f**) Results generated by the model described in (a-b) with a single fixed set of parameters for all the panels. Model recapitulates: (**c**) the population-growth curves (compare with Fig. 2a), (**d**) the phase diagram (compare with Fig. 2d), (**e**) population-density dependent deaths (compare with Fig. 3b), (**f**) population half-life (based on cell deaths during the first day of incubation - compare with Fig. 3d), the number of survivors decaying over time as a heavy-tailed function (see Supplementary Fig. 14), and single-cell-level data on growths (compare Supplementary Fig. 14 with Supplementary Fig. 6). Also see Supplementary Fig. 14.

Strikingly, after using our data for the wild-type strain to fit the one free parameter, our highly constrained model qualitatively - and quantitatively - recapitulated all the main experimental data (Figs. 5c-f and Supplementary Fig. 14). Specifically, the model recapitulates the population-level growth kinetics with the distinct growth phases (Fig. 5c, compare with Fig. 2a), the single-cell-level growth kinetics for each growth phase (Supplementary Fig. 14, compare with Supplementary Fig. 6), the phase diagram (Fig. 5d, compare with Fig. 2d), the population-density dependent deaths (including the number of survivors decreasing over time in a heavy-tailed manner) (Fig. 5e, compare with Fig. 3b), and the relationship between temperature and the population-density-dependent deaths (Fig. 5f, compare with Fig. 3d). Moreover, we used a mathematical argument to establish that our model is the simplest possible class of model that can explain our data (see Supplementary text). To intuitively see how the model reproduces all the experimental data, note that the probability that a cell replicates is initially zero when the population first enters a high-temperature environment, since the glutathione concentration is initially zero. The probability increases over time as the glutathione concentration increases due to the cells constantly secreting glutathione (Fig. 5b and Supplementary Fig. 14). However, the probability for a cell to die starts at a nonzero value when the population first enters a high-temperature environment, since it is set only by temperature. This probability remains constant over time since its value is independent of the extracellular glutathione concentration (Fig. 5b and Supplementary Fig. 14). Therefore, there exists a specific glutathione-concentration - a threshold concentration - above which the probability of replicating exceeds the probability of dying, thereby resulting in a population growth. This sets up a “race” in which a population of cells, starting without any extracellular glutathione, must realize the threshold concentration before going extinct. There are initially more cell deaths than cell replications and thus there is a “ticking time bomb” until extinction. As a result, if the initial population-density is sufficiently high, then population growth wins the race - this occurs in the deterministic-growth phase (Supplementary Fig. 14). If the initial population-density is sufficiently low, then population extinction wins the race - this occurs in the no-growth phase (Supplementary Fig. 14). For a population that starts with an intermediate density, the glutathione concentration gets close to the threshold concentration by the time there are very few alive cells remaining. At this point in time, any subsequent, small changes in the number of alive cells determines whether the probability of a cell replicating exceeds the probability of a cell dying (Supplementary Fig. 14). Hence, when a population would grow and if it can even grow are both completely unpredictable because the population-level behavior here is highly sensitive to the stochastic behaviors of the few alive cells (*40*) - this occurs in the random-growth phase. In the model, at sufficiently high temperatures (i.e., above ∼40.3 °C), the probability of dying exceeds the highest probability that a cell can have for replicating (Fig. 5b - grey dashed line). This occurs precisely at the fold-bifurcation point in the phase diagram. In the no-growth phase, the model also reproduces the number of survivors decreasing over time in a heavy-tailed manner (Fig. 5e and Supplementary Fig. 14). The mechanism here is that cells keep dying as time passes - due to the probability of dying remaining constant over time - but, for the still-alive cells, the probability of replicating increases over time due to each surviving cell continuously secreting glutathione into the environment, making replication more likely for each alive cell as time passes. The competition between the two - a constant probability of dying and an initially lower probability of replicating that gradually approaches the probability of dying - results in a population whose approach to extinction continuously slows down over time, leading to the number of survivors decreasing over time in a heavy-tailed manner that we experimentally observed (see Supplemental text). Taken together, these results show that our relatively simple model, with the secreted and accumulated glutathione helping the current and future generations of cells replicate as the primary ingredient, recapitulates all the main features of cell replications and deaths that we experimentally observed (Figs. 5c-f).

### Extracellular glutathione is both necessary and sufficient for cooperative cell-replication at high temperatures

Our mathematical model did not require knowing exactly how secreting glutathione helps cells replicate at high temperatures. We next sought to gain molecular insights into extracellular glutathione promoting cell replications at high temperatures. To recap, we have shown so far that, at high temperatures, yeasts secrete glutathione (Fig. 4e) and that a sufficient concentration of extracellular glutathione is required to induce a population growth (Fig. 4f). These two results show that extracellular glutathione is sufficient for the cells to help each other replicate at high temperatures. Next, we sought to address if extracellular glutathione is also necessary - in addition to being sufficient - for yeasts to help each other replicate. To test this, we inactivated (“removed”) all extracellular glutathione by adding a chemical that masked glutathione in the liquid media. This “masking reagent” (called 1-Methyl-2-vinylpyridinium (M2VP)) rapidly scavenges all extracellular glutathione, without interfering with the intracellular glutathione and any intracellular processes (Supplementary Fig. 15) (*41,42*). As an example, at ∼39 °C, we added the masking agent to the liquid media of deterministically growing populations (Fig. 6a). We found that these populations stopped growing after we added the masking agent. Hence, we can conclude that glutathione is not only sufficient (Fig. 4f) but is also necessary for cell replications at high temperatures. In other words, glutathione is the only molecule that determines whether a population grows or not at high temperatures. To see this, note that if any other molecules were necessary for growth at high temperatures, then masking the glutathione should not have prevented the population growths. Conversely, note that if any other secreted molecules besides glutathione were sufficient for inducing population growth at high temperatures, then removing glutathione alone should not have stopped the population growth. Finally, we found that adding the masking agent to the liquid media of a growing population at 30 °C did not affect the population growth - the population continued to grow in the presence of the masking agent at 30 °C (Supplementary Fig. 15). Taken together, these results establish that extracellular glutathione is both necessary and sufficient for inducing cell replications at high temperatures (above ∼36.7 °C) but that it is not necessary at temperatures below ∼36.7 °C.

**Figure 6.**
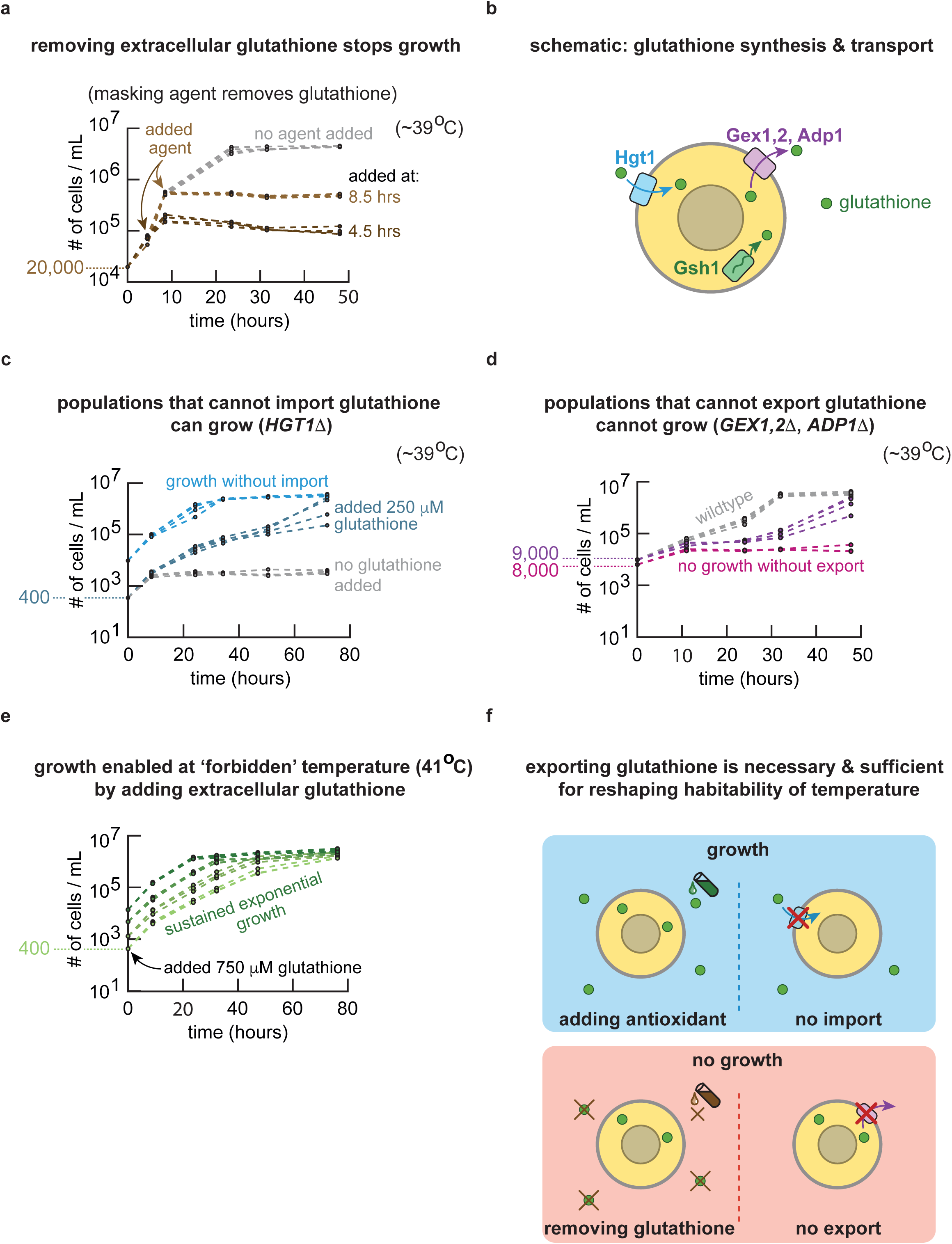
Budding yeast exports glutathione whose extracellular role – not intracellular roles - as an antioxidant enables yeasts to survive high temperatures. (**a**) At 39.2 °C. Curves show wild-type populations (all initially at ∼20,000 cells/mL) that should deterministically grow if left alone (as in grey curve). A glutathione masking agent (M2VP) – which inactivates extracellular glutathione - was added after 4.5 hours (dark brown) or 8.5 hours (light brown) of incubation (see Methods). Grey curve shows populations that did not receive the masking reagent. Each color shows four replicate populations. Also see Supplementary Fig. 15. (**b**) Schematic showing how the budding yeast synthesizes, imports, and exports glutathione. Glutathione is intracellularly synthesized via an enzyme (γ-glutamylcysteine synthetase) encoded by *GSH1*. Glutathione is imported by a proton-coupled glutathione-importer encoded by *HGT1*. Glutathione is exported by numerous exporters (not all shown here), including proton antiporters encoded by *GEX1,2* and an ATP-dependent exporter encoded by *ADP1*. (**c**) At 39.2 °C. Light blue curves show deterministically growing populations of a mutant strain (*HGT1Δ*-strain) that cannot import glutathione (initially at ∼10,000 cells/mL). Grey curves at the bottom show the mutant populations (initially ∼400 cells/mL) incubated without any glutathione added. Dark blue curves in the middle show the mutant populations (initially ∼400 cells/mL) incubated with 250μM of glutathione added to the media. Each color shows four replicate populations. (**d**) At 39.2 °C. Populations of a mutant strain that lacks some of the main glutathione exporters (*GEX1,2Δ*-*ADP1Δ*-strain) (initially ∼9,000 cells/mL (purple curves in the middle) or ∼8,000 cells/mL (pink curves at the bottom)). Grey curves at the top show, as a comparison, the wild-type populations (initially ∼9,000 cells/mL). Each color shows at least four replicate populations. (**e**) At 41 °C. Wild-type populations of various initial densities (from ∼400 cells/mL (lightest green curves) to ∼14,000 cells/mL (darkest green curves)) grown in media supplemented by 750μM of glutathione. Each color shows four replicate populations. (**f**) Cartoon illustrating mechanisms deduced in (a-e). Exporting glutathione is necessary and sufficient for budding yeasts to reshape the habitability of temperature. For yeasts to survive and grow at high temperatures, extracellular glutathione is (1) necessary since blocking glutathione or blocking glutathione-export stops yeast’s growths and is (2) sufficient since adding glutathione or blocking glutathione-import enables the yeasts to grow at high temperatures.

### Transport and function of glutathione at high temperatures

Now that we have identified extracellular glutathione to be the sole factor that dictates whether a population grows or not at high temperatures, we sought to gain molecular insights into how the extracellular glutathione can promote cell replications at high temperatures. Myriad proteins regulate the biosynthesis of glutathione and its transport in and out of the cell (Fig. 6b). The enzyme Gsh1 is required for synthesizing glutathione and thus a strain lacking the *GSH1* gene (i.e., *GSH1Δ*) cannot synthesize glutathione (*43*). The budding yeast requires the importer, Hgt1, to import glutathione from outside and thus a strain lacking the *HGT1* gene (i.e., *HGT1Δ*) is unable to import detectable amounts of glutathione (*44*). Finally, there are various glutathione exporters that each contribute to the overall secretion of glutathione (*36*). Of these, Gex1 and Gex2 are antiporters - they import protons while actively exporting glutathione - and knocking both the *GEX1* and *GEX2* genes cause a reduction of about 40% in the rate of glutathione export (*45*). The budding yeast also actively export glutathione via several ABC transporters such as Yor1 (*36*) and Adp1 (*46*). To genetically manipulate the synthesis, import, and export of glutathione at high temperatures, we constructed mutant strains that were either unable to synthesize glutathione (*GSH1Δ*-strain), or unable to import glutathione (i.e., *HGT1Δ*-strain), or had severely reduced ability to secrete glutathione (*GEX1,2Δ*-*ADP1Δ*-strain).

We first used the mutant strain that could not import glutathione (*HGT1Δ*-strain) to determine whether yeasts need to import their secreted glutathione in order to grow at high temperatures. We found that this mutant has the same population-density-dependent growths at high temperatures, just like the wild-type strain (Fig. 6c). Similar to the wild-type strain at sufficiently at low population densities, mutant populations cannot grow if they start with sufficiently low densities. But they can grow if we add enough glutathione to their liquid media (Fig. 6c). Together, these results establish that yeasts do not need to import secreted glutathione in order to replicate at high temperatures. This, in turn, means that glutathione’s extracellular action alone must be responsible for promoting cells replications at high temperatures.

Since extracellular glutathione, without being imported, determines a population growth at high temperatures, we reasoned that a mutant strain with a significantly reduced ability to secrete glutathione (*GEX1,2Δ*-*ADP1Δ*-strain) may have a reduced ability to replicate at high temperatures. Indeed, we found that while populations of this mutant can still grow at high temperatures (Fig. 6d), they have a reduced ability to replicate compared to the wild-type strain. Concretely, this means that while a wild-type population can deterministically grow at a given high temperature, a mutant population that starts with the same density as the wild-type population can at most randomly grow at the same temperature (i.e., the mutant achieves deterministic growth only with higher initial population-densities). Likewise, while a wild-type population can grow randomly at a given high temperature, a mutant population that starts with the same density cannot grow (i.e., the mutant strain needs to start with a higher population density to escape the no-growth phase). Thus, consistent with glutathione’s extracellular action – rather than intracellular action – promoting cell replications, these results together show that reducing the yeast’s ability to export glutathione reduces the ability of yeast populations to grow at high temperatures.

The mutant strain with a reduced glutathione-export (*GEX1,2Δ*-*ADP1Δ*-strain) still secreted measurable amounts of glutathione at high temperatures (Supplementary Fig. 16), which is expected as it has other glutathione exporters (e.g., Yor1p). We wondered how much of this secretion – and wild-type strain’s glutathione secretion - was because glutathione simply leaked out of the cells at high temperatures due to, for example, heat degrading the integrity of cell membranes. To test this, we used a standard test – staining by propidium iodide (*47*) - to check how many cells, in a mutant population (*GEX1,2Δ*-*ADP1Δ*-strain) that was growing at a high temperature, allowed propidium iodide to diffuse through their potentially leaky (compromised) membranes to stain their nucleic acids. We found that growing mutant populations at high temperatures actually have intact membranes as almost none of the cells in the populations were stained by propidium iodide (Supplementary Fig. 16). Thus, glutathione does not simply leak out of the cells at high temperatures – cells must use exporters to secrete glutathione at high temperatures.

Finally, we used the mutant strain that is unable to synthesize glutathione (*GSH1Δ*-strain) to double check that the wild-type cells secrete glutathione at high but not at low temperatures. We found this earlier by directly measuring the concentration of extracellular glutathione in growing populations. As is well known, the *GSH1Δ*-strain cannot survive or grow at any temperature, including at 30 °C, unless one supplements its growth medium with glutathione (Supplementary Fig. 17) (*43*). This is due to glutathione’s essential intracellular roles which are not related to heat-shock responses (*43*). As in our media-transfer experiments (Fig. 4b, Supplementary Fig. 10), we took the liquid media of wild-type populations that were growing at a high temperature (e.g., at ∼39 °C), then at 30 °C, we transplanted fresh populations of the *GSH1Δ*-cells that were starved of extracellular glutathione (Supplementary Fig. 17). We found that these mutant populations, which cannot synthesize but need glutathione to survive, grew in the transferred media. Thus, the media taken from the wild-type populations that were growing at high temperatures must contain glutathione (i.e., wild-type cells indeed secrete glutathione at high temperatures). Moreover, we found that the glutathione-starved *GSH1Δ*-cells did not grow in media that we took away from wild-type populations that were growing at 30 °C. This means that the wild-type cells indeed do not secrete glutathione at 30 °C. Taken together, these results again confirm our earlier conclusion: wild-type yeasts secrete glutathione only at high temperatures (above ∼36 °C)

### Yeasts can replicate at “unlivable” temperatures

Finally, since extracellular glutathione is both necessary and sufficient for population growths at high temperatures, we reasoned that wild-type populations neither randomly nor deterministically grow at temperatures above 40.3 °C - according to the wild-type strain’s phase diagram (Fig. 2d) - because they cannot accumulate sufficient concentrations of extracellular glutathione at temperatures beyond 40.3 °C. Note that 40.3 °C is where the fold-bifurcation point is located and that beyond 40.3 °C, only the no-growth phase exists in the wild-type strain’s phase diagram (Fig. 2d). We therefore wondered if we could help wild-type populations grow at temperatures higher than 40.3 °C by supplementing their media with glutathione. We tested this idea by adding a saturating amount of glutathione to the liquid media of wild-type populations at 41 °C (Fig. 6e). We found that these populations – nearly 1 °C above the supposedly maximum growth-permitting temperature - were now able to grow until they reached the carrying capacity. Thus, deterministic growths are possible at a temperature that is nearly one degree Celsius higher than the maximum temperature (40.3 °C) for the wild-type strain, if we help them accumulate extracellular glutathione.

Taken together, our experiments with the three mutant strains above establish that budding yeasts must secrete and extracellularly accumulate glutathione in order to replicate at high temperatures and that glutathione’s extracellular action alone – not its intracellular action - is responsible for promoting cell replications at high temperatures (Fig. 6f). The latter, coupled with the fact that adding another antioxidant (ascorbic acid) alone is sufficient for inducing population growths at high temperatures (Fig. 4d), lends support to the conclusion that extracellular glutathione promotes cell replications by reducing harmful extracellular agents such as reactive oxygen species at high temperatures.

## DISCUSSION

### Expanding the role of glutathione as mediator of cooperative cell-replication and extender of population lifespan at high temperatures

Glutathione is a well-known tri-peptide that is essential for many organisms, including humans (*34*). It is central to diverse processes such as combating cancer (*48*) and neurodegenerative diseases (*49*), slowing down aging (*50*), detoxifying the liver (*51*), and - relevant to our study - protecting cells from heat-induced reactive oxygen species that damage cells (*28–30, 52*). For the budding yeast, much of the focus on glutathione has been on its intracellular role. Intracellularly, glutathione indeed has essential roles in iron metabolism (*37*), thiol-redox regulations (*53*), protecting against reactive oxygen species as an antioxidant (*34,38*), detoxifying the environment poisoned by xenobiotics (*54*) and heavy metals such as cadmium (*55*), and responding to sulphur starvation (*56*) and nitrogen starvation (*57*). But glutathione’s extracellular role for budding yeast - both at high temperatures and in other contexts - remains poorly understood and has received relatively little attention compared to its intracellular roles. Among the studies that examined extracellular glutathione for yeasts are a recent study showing that extracellular glutathione can protect yeasts from toxic arsenite (*36*) and a work showing how extracellular glutathione aids in dealing with nutrient imbalance for certain mutant yeasts that have been evolved to secrete glutathione at 30 °C (*58*). Moreover, there have been some reports on how yeasts, during stationary-phase that follows a diauxic shift at 30 °C, uptake and secrete low amounts of glutathione (*35*). Indeed, we confirmed that stationary-phase populations, after many hours, can build up to ∼2 μM extracellular glutathione at 30 °C but that log-phase populations do not secrete glutathione at 30 °C. What was underappreciated before, which we have discovered here, is that yeasts secrete glutathione at high temperatures - during log-phase growth and stationary phases - and that this leads to their cooperative replications and an indefinite extension of a population’s lifespan (in the form of heavy-tailed population decay) at high temperatures.

Furthermore, we found that extracellular glutathione at high temperatures mirrors several known features of intracellular glutathione at 30 °C. Firstly, at 30 °C, researchers have found that yeasts maintain intracellular glutathione at high, millimolar concentrations and keep most of it in its reduced form by recycling it in a NADPH-dependent manner (*34, 59*). This is because glutathione in its reduced form, rather than its oxidized form, is essential for capturing reactive oxygen species (*34*). Analogously, we found that, at high temperatures, log-phase wild-type yeasts maintain ∼77% of their extracellular glutathione in the reduced form and the rest in the oxidized form (Supplementary Fig. 13). Secondly, we established that accumulating sufficiently large amounts of extracellular glutathione is crucial to yeasts’ survival at high temperatures, just as their survival depends on building up of intracellular glutathione. Specifically, high temperatures cause yeasts to respire, which in turn causes the mitochondria to produce reactive oxygen species (*33*). Thus, it makes sense that - as previous studies established - high temperatures cause yeasts to produce high amounts of intracellular glutathione to survive oxidative stress (*32*). Our work extends this finding by revealing that the budding yeast must export their intracellular glutathione in order to replicate and survive at high temperatures. Moreover, our work establishes that accumulating extracellular glutathione, even by the alive cells in the no-growth-phase populations, extends the population’s lifespan in a manner that depends on the population’s history - namely, how many cells the population started with and how the extracellular glutathione has been accumulating over time. In this way, glutathione extends population’s lifespan in a manner that depends on the population’s history. Even if the population currently has very few alive cells remaining, these cells benefit from the extracellular glutathione that the past generations of cells had secreted and accumulated before they died. Given glutathione’s utility and secretion in other organisms - for example, human lung epithelial cells secrete it when they are exposed to asbestos (*60*) - future studies may show that extracellular glutathione mediates cooperative survival in other organisms at high temperatures or for other stresses.

### Why cooperate instead of autonomously combatting heat-induced damages?

A natural question is why yeasts secrete glutathione at high temperatures to cooperatively survive instead of each yeast cell autonomously combating heat shocks by intracellularly keeping all the glutathione for itself. A major advantage of secreting glutathione is that glutathione can extracellularly accumulate, thereby allowing cells to help their future generations of cells to replicate, long after the current generation of cells have died off. Indeed, our model shows that it is the accumulation of extracellular glutathione that is responsible for a population to decelerate and eventually stop its approach to extinction.

### Relation to dynamical systems theory

The term, fold-bifurcation, comes from dynamical systems theory – a branch of mathematics that focuses on explaining all possible ways that a set of mathematical variables can change over time and the qualitative features that describe them, often by invoking geometric depictions such as phase diagrams. This theory can describe many biological systems, from the classical predator-prey systems (*61–63*) to – as recently verified experimentally - microbial populations that are on the verge of extinction due to environmental perturbations (*64–68*). In the phase diagram that we uncovered (Fig. 2d), the shape of the phase boundaries that converge at the fold-bifurcation point mathematically resembles the shape of phase boundaries that merge into a fold-bifurcation point in another phase diagram - one that describes a yeast population for which one simulates the rate of cell deaths by periodically flushing out the liquid culture of cells and for which the replicability of cells depends on the yeasts collectively breaking down extracellular sucrose into glucose that they can import (*64*). A yeast population mathematically behaves in similar ways in both systems. In particular, both systems exhibit “critical slowing down” (*64,69,70*) which, in our case, manifests as the yeast-population’s initial half-life being infinite at the fold-bifurcation point (Fig. 3d) - this is a direct consequence the phase diagram having a fold-bifurcation point (Fig. 2d) according to dynamical systems theory (*69*). Crucially, the resemblance between the two dynamical systems – yeasts at high temperatures and yeasts that collective breakdown sucrose – arises because cells cooperate in both systems. In such populations of cells whose survival depend on cooperating amongst them, one typically observes minimal population-density that is required for survival. By providing another example of cooperation among microbes, our work thus adds to the on-going efforts by researchers to experimentally map phase diagrams that depict how populations of cooperating microbes behave when they are on the verge of going extinct (*64–68*).

### Summary and outlook

With budding yeasts, we have shown that cells can avoid extinctions at high temperatures by helping each other and their future generations replicate. As a consequence of this cooperative behavior, a temperature which is unlivable for one population of cells can be habitable for another population of cells of the same species, depending on whether there are sufficiently many cells that are cooperating to keep the population alive. Specifically, we have shown that yeasts at high temperatures export glutathione and that the extracellularly accumulating glutathione, in turn, increases the probability of a cell replicating. By accumulating extracellular glutathione, yeasts also aid their future generations of cells to replicate, long after their own deaths. Conversely, yeast populations cannot grow if we remove extracellular glutathione. Thus, extracellular glutathione - in fact, exporting of glutathione - is both necessary and sufficient for yeasts to grow at high temperatures. Taken together, our experiments and mathematical model replaces the conventional picture in which whether a yeast survives a high temperature or not depends on whether it can autonomously use its heat-shock response system to combat heat-induced damages. A surprising consequence of our model, which recapitulates all the main features of the experimental data, is that yeasts can, in fact, replicate - albeit with a vanishingly low probability - at extremely high temperatures (e.g., 45 °C) for which the population cannot grow (due to only the no-growth phase existing in the population-level phase diagram) (Supplementary Fig. 14). Experimentally, we could indeed increase, by nearly one degree Celsius, the maximum temperature at which yeast populations can grow to a carrying capacity by supplementing the growth media with glutathione (Fig. 6e). Previous studies examined cell growth (*71–74*), gene regulations (*75,76*), and metabolite exchanges among microbes (*77–80*) for various species at their conventionally-defined habitable temperatures (i.e., habitability of temperature defined by a cell-autonomous view). Our work encourages re-examining these features at high temperatures that were previously dismissed as unlivable but which may, in fact, be habitable for sufficiently large populations due to the cells cooperating in ways that may have been overlooked.

A common explanation for why cells cannot replicate at high temperatures has been that a few, yet essential, proteins unfold at these temperatures and that the cells’ autonomous heat-shock response system fails to deal with these unfolded proteins. While crucial proteins do unfold at certain high temperatures in budding yeast (*6*), our work shows that the reason for yeasts failing to replicate at high temperatures is more intricate. For one, we find that if there is a sufficient amount of extracellular glutathione then budding yeasts can, in fact, replicate at extremely high temperatures (i.e., above 41 °C) for which the essential proteins for yeast have been reported to be unstable (Fig. 6e and Supplementary Fig. 14). Secondly, our work establishes that glutathione’s extracellular action as an antioxidant determines the replicability of yeasts and yeast population’s longevity at high temperatures. Simply put, yeasts fail to replicate at high temperatures because there are not enough of them working together to help each other replicate and not enough of them leaving behind extracellular glutathione to help their future generations of cells replicate.

From a standpoint of physics, we can interpret the random-growth phase in the phase diagram as a boundary formed by a co-existence of the deterministic growth and no-growth phases. A population that is at the endpoint of this boundary at the fold-bifurcation point can indefinitely maintain a steady number of cells without either growing or becoming extinct - its population density can fluctuate but does not change over time on average. The fold-bifurcation point and the number of surviving cells decreasing over time as a heavy-tailed function for populations that are near the fold-bifurcation point in the phase diagram, remind us of the power-law functions that often describe the behaviors of non-living systems at critical points which are associated with phase transitions (*17–19*). By exploiting these resemblances and using our data and model, a future study might further advance theories of non-equilibrium phase transitions (*81*) that pertain to biologically realistic, self-replicating systems that constantly drive and keep themselves out of equilibrium by using thermal energy. Relatedly, a chronologically ageing population of yeasts may also have the number of survivors decreasing over time as a heavy-tailed function (*82*), like the number of survivors in non-growing populations incubated at high temperatures. Thus, an improved understanding of how thermal energy reliably drives self-replication of cells may provide insights into the dynamics of cellular ageing as well. Furthermore, understanding how cells can work together to collectively combat extreme temperatures, as we have done here, may suggest intervention mechanisms to combat climate change as well as advance our understanding of how temperature and climate change can impact unicellular life and multicellular communities.

## Methods

### Growth media and strains

The “wild-type”, haploid yeast strain that we used is from Euroscarf with the official strain name “20000A”. It is isogenic to another laboratory-standard haploid yeast “W303a”, and has the following genotype: *MATa*; *his3-11_15; leu2-3_112; ura3-1; trp1Δ2; ade2-1; can1-100*. We built the two strains that constitutively expressed GFP by first using PCR to insert a functional *ADE2* gene into the locus of the defective *ADE2* gene in the wild-type strain, by a homologous recombination, so that the red pigments that would have accumulated without the *ADE2* insertion no longer existed (i.e., the strain can now synthesize adenine). We could thus detect their GFP fluorescence without interferences from the red pigments. After replacing the defective *ADE2* locus with a functional *ADE2*, we constructed the 1x-GFP and 100x-GFP strains (see GFP-expression levels in Fig. S4a) by integrating a single-copy of an appropriate, linearized yeast-integrating plasmid at the *HIS3* locus on the chromosome. Specifically, the 1x-GFP strain had its GFP expression controlled by the constitutive promoter of yeast’s *KEX2* gene (621 bases upstream of its ORF) which was on a yeast-integration plasmid (*83*) that constitutively expressed *HIS3* (from *C. glabrata*) and integrated into the non-functional *HIS3*-locus of the wild-type strain by a homologous recombination. The 100x-GFP strain had its GFP expression controlled by a strong constitutive promoter pGPD1 (*83*) which was on the same plasmid as the one for the 1x-GFP strain except that the *KEX2* promoter was swapped with the *GPD1* promoter. We cultured all yeasts in defined, minimal media that consisted of (all from Formedium): Yeast Nitrogen Base (YNB) media, Complete Supplement Mixture (CSM) that contained all the essential amino acids and vitamins, and glucose at a saturating concentration (2% = 2 g per 100 mL). The agar pads, which we used for growing yeast colonies, contained 2%-agar (VWR Chemicals), Yeast Extract and Peptone (YEP) (Melford Biolaboratories Ltd.), and a 2%-glucose.

### Growth experiments

In a typical growth experiment, we first picked a single yeast colony from an agar plate and then incubated it at 30 °C for ∼14 hours in 5-mL of minimal medium, which contained all the essential amino acids and a saturating concentration of glucose (2%). Afterwards, we took an aliquot of a defined volume from the 5-mL culture (typically 20 μL), and then flowed it through a flow cytometer (BD FACSCelesta with a High-Throughput Sampler) to determine the 5-mL culture’s population density (# of cells/mL). We then serially diluted the culture into fresh minimal media to a desired initial population-density for a growth experiment at various temperatures. Specifically, we distributed 5-mL of diluted cells to individual wells in a “brick” with twenty-four 10-mL-wells (Whatman: “24-well x 10mL assay collection & analysis microplate”). This ensured that we had 8 identical replicate cultures for each initial population-density (e.g., in Figure 2a-c). We sealed each brick with a breathable film (Diversified Biotech: Breathe-Easy), covered it with a custom-made Styrofoam-cap for insulation, and incubated it in a compressor-cooled, high-precision thermostatic incubators (Memmert ICP260) that stably maintained their target temperature throughout the course of our growth-experiments, with a typical standard deviation of 0.017 °C over time (deviation measured over several days - see Supplementary Fig. 2). Throughout the incubation, the cultures in the brick were constantly shaken at 400 rpm on a plate shaker (Eppendorf MixMate) that we kept in the incubator. To measure their population densities, we took a small aliquot (typically 50 μL) from each well, diluted it with PBS (Fisher Bioreagents) into a 96-well plate (Sarstedt, Cat. #9020411), and then flowed it through the flow cytometer which gave us the # of cells/mL. We determined the growth rates by measuring the maximum slope of the log-population density after their initial, transient growths.

### Flow cytometry

The flow cytometer that we used was a BD FACSCelesta with a High-Throughput Sampler and lasers with the following wave lengths: 405 nm (violet), 488 nm (blue), and 561 nm (yellow/green). We calibrated the FSC and SSC gates to detect only yeast cells (FSC-PMT=681V, SSC-PMT=264V, GFP-PMT=485V, mCherry-PMT=498V. As a control, flowing PBS yielded no detected events). The number of cells/mL that we plotted in our growth experiments is proportional to the number of events (yeast cells) that the flow cytometer measured in an aliquot of cells with a defined volume. We measured the GFP fluorescence with a FIT-C channel and the “red cells” (Figure S6) with a mCherry channel. We analysed the flow cytometer data with a custom MATLAB script (MathWorks).

### Measuring number of surviving cells

For Figures 3a-b and Figures S7-S8, we prepared 250-mL cultures of wild-type cells in 500-mL Erlenmeyer flasks. We placed a constantly spinning magnetic stir-bar at the bottom of the flasks and placed each flask on top of spinning magnets (Labnet Accuplate - at 220 rpm) inside the thermostatic incubators (Memmert ICP260) that we set at desired high-temperatures. For Figure 3d, we prepared a brick with cultures as described in the “Growth Experiments” section in order to have multiple replicate populations and to compare the different population densities. For every time point in Fig. 3 and Supplementary Figs. 7-8, we ensured that these populations were not growing (i.e., all populations were in the no-growth phase after a transient growth) by using the flow cytometer to measure their population-densities over time to verify that their population-densities indeed remained constant over time. For the first 48 hours of incubation, we measured the number of Colony Forming Units (CFUs) by taking out a small-volume aliquot from the liquid cultures at high temperatures and distributed droplets from a serial dilution of the aliquot across an agar pad (2% glucose with YEP) that we then incubated in 30 °C for several days until (no) colonies appeared. When there were few surviving cells per mL - especially for the last time points in each experiment - we determined, in parallel to the plating method, the number of CFUs by transferring an appropriate volume of the liquid cultures from the incubator to an Erlenmeyer flask and then diluting it with the same volume of fresh minimal media. We sealed this flask with a breathable film (Diversified Biotech: Breathe-Easy) and then left it still without stirring, on a benchtop at ∼24-30 °C - we checked that slightly lower temperatures (e.g., room temperatures) did not affect colony-forming abilities - which allowed any surviving cells to settle down to the bottom of the flask and form colonies. We counted the number of colonies at the bottom of the flask - this is the value that we plotted as the last time points in each experiment (Fig. 3 and Supplementary Figs. 7-8).

### Cell-transfer experiments

We incubated a 24-well brick that contained liquid cultures, each of which were in a deterministic-growth phase, at a desired temperature (e.g., 10,000 cells/mL at 39.2 °C). We incubated the brick containing these liquid cultures in the thermostatic incubators (Memmert ICP260) as described in the “Growth experiments” section. About 48 hours after the incubation, we took aliquots from the cultures that were growing in mid-log phase (as checked by flow cytometry) and then diluted each of them into fresh 5-mL minimal media that were in 24-well bricks so that these newly created populations were in the no-growth phase at the same temperature as the original population that they came from (400 cells/mL at 39.2 °C). We sealed the 24-well brick with a breathable film (Diversified Biotech: Breathe-Easy) and then incubated them at the same temperature as the original population. We performed the growth experiments with these new populations as described in the “Growth experiments” section.

### Medium-transfer experiments

Details are also in Supplementary Fig. 10. At a given temperature, we first grew populations in the deterministic-growth phase (e.g., initial population-density of 30,000 cells/mL at 39.2 °C). We used a flow cytometer to measure their growing population-densities at different times so that we knew in which part of deterministic growth they were in (e.g., mid-log phase). We then transferred each liquid culture to a 50-mL tube (Sarstedt) and centrifuged it so that the cells formed a pellet at the bottom of the tube. We then took the resulting supernatant, without the cell pellet, and flowed it through a filter paper with 200-nm-diameter pores (VWR: 150-mL Filter Upper Cup) to remove any residual cells from the supernatant. After filtering, we flowed an aliquot of the filtered media through a flow cytometer to verify that there were no cells left in the filtered media. We incubated fresh cells into these filtered media (instead of into fresh minimal media) and proceeded with a growth experiment at a desired temperature as described in the “Growth experiments” section.

### Measuring the depletion of extracellular nutrients

Details are also in Supplementary Fig. 11. We prepared various growth media by diluting the minimal media (SC media) by various amounts with water. These diluted SC-media were each supplemented with a 2% glucose. Next, we incubated fresh cells in these diluted SC-media at the desired temperature (e.g. 39.2 °C) as described in the “Growth experiments” section. We compared populations of cells that initially had 400 cells/mL (this corresponds to a no-growth phase, see Fig. 2d) with populations that initially had 10,000 cells/mL (this corresponds to a deterministic-growth phase, see Fig. 2d) in order to confirm that cells were still able to grow in these media. Similarly, we also varied the amounts of glucose that we supplemented to SC-media.

### Measuring concentration of extracellular glutathione

To quantify the concentration of extracellular glutathione, cells were removed from their liquid media by flowing the liquid cultures that contained cells through a 0.45−μm pore filter (VWR, cellulose-acetate membrane). To ensure and verify that there were no cells remaining in the filtered media, we flowed the filtered media through a flow cytometer. The flow cytometer indeed did not detect any cells in the filtered media. We measured concentrations of glutathione in the filtered media as described in the manufacturers’ protocol (38185 quantification kit for oxidized and reduced glutathione, 200 tests). We used “BMG Labtech Spectrostar Nano” to measure the optical absorbance at 415-nm. As a background subtraction (blank) for all absorbance measurements, we subtracted the absorbance that we obtained by applying the assay to fresh minimal medium which does not have any glutathione (the background absorbance could come from, for example, cysteine in the minimal media). We subsequently determined the concentrations of extracellular glutathione by using a calibration curve that we constructed by measuring the absorbance at 415-nm for known amounts of glutathione that we added by hand into a buffer provided by the manufacturer.

### RNA-seq

For each temperature that we studied, we collected cells in 50-mL tubes and spun them in a pre-cooled centrifuge. We then extracted RNA from each cell-pellet with RiboPure Yeast Kit (Ambion, Life Technologies) as described by its protocol. Next, we prepared the cDNA library with the 3’ mRNA-Seq library preparation kit (Quant-Seq, Lexogen) as described by its protocol. Afterwards, we loaded the cDNA library on an Illumina MiSeq with the MiSeq Reagent Kit c2 (Illumina) as described by its protocol. We analysed the resulting RNA-Seq data as previously described (*84*): We performed the read alignment with TopHat, read assembly with Cufflinks, and analyses of differential gene-expressions with Cuffdiff. We used the reference genome for *S. cerevisiae* from ensembl. We categorized the genes by the Gene Ontologies with AmiGO2 and manually checked them with the Saccharomyces Genome Database (SGD).

### Glutathione masking experiment

We incubated a brick of liquid cultures that were in the deterministic-growth phase (20.000 cells/mL) at 39.2 °C. After some time (e.g., 8.5 hours afterwards), we added 750μM of 1-Methyl-2-vinylpyridinium (M2VP, Sigma-Aldrich Cat. No. 69701), which is a thiol scavenging agent that rapidly masks reduced glutathione (*42*) and proceeded with the experiment as described in the “Growth experiments” section. Identical replicate cultures, that did not receive the M2VP, were used as a reference.

### Mutant yeasts

We constructed mutant strains that could not synthesize glutathione or could not import or export glutathione. Primers were designed with a 50-60bp sequence that was either homologous to the 50-60bp that is upstream of the desired gene’s start codon or downstream of the desired gene’s stop codon. These primers were used to amplify a selection marker by PCR, resulting in a PCR product that contained a selection marker and whose ends were homologous to the flanking regions of the gene to be knocked out. The wild-type strain (W303) was grown overnight in a 5-mL YPD in a rotator (40 rpm) at 30 °C, and subsequently transformed with the PCR fragment using standard methods of yeast cloning. The biosynthesis mutant (*GSH1Δ*-strain) was constructed by removing the *GSH1* gene from W303. The import mutant (*HGT1Δ*-strain) was constructed by removing the *HGT1* gene. The export mutant (*GEX1,2Δ*-*ADP1Δ*-strain) was constructed by removing, sequentially, the *GEX1* gene, then the *GEX2* gene, and then the *ADP1* gene. The resulting transformants were grown on YPD selection plates, and knockouts were verified by PCR.

### Measuring integrity of cell membrane

Cells of the *GEX1,2Δ*-*ADP1Δ*-strain were incubated in liquid media at 39.2 °C (3,000-10,000 cells/mL; corresponds to a random-growth phase). We took aliquots of these cultures and then stained them with 1 μg/mL of propidium iodide (Thermo Fisher Cat. No. P3566). We then flowed these stained aliquots through a flow cytometer. The flow cytometer measured the number of cells that were unstained by the propidium iodide (i.e., cells whose membranes were intact).

### Mathematical model

Derivations of equations, a detailed description of the mathematical model, and the parameter values used for simulations are in the Supplemental text.

## Supporting information

Supp. Figs and mathematical model

## Supplemental Information

- Supplementary Figures 1-17
- Supplementary Text: Detailed description of the mathematical model

## Acknowledgements

We thank Shalev Itzkovitz and Arjun Raj for insightful comments on our manuscript. We also thank the members of the Youk laboratory for helpful discussions and Mehran Mohebbi for help with initial experiments. H.Y. was supported by the European Research Council (ERC) Starting Grant (MultiCellSysBio, #677972), the Netherlands Organisation for Scientific Research (NWO) Vidi award (#680-47-544), the CIFAR Azrieli Global Scholars Program, and the EMBO Young Investigator Award.

## Author contributions

H.Y. initiated this research and performed the initial experiments. D.S.L.T. subsequently developed the project with guidance from H.Y. D.S.L.T. performed the experiments, developed the mathematical model, and analysed the data with advice from H.Y. D.S.L.T. and H.Y. discussed and checked all the data and wrote the manuscript.

## DATA AVAILABILITY

The authors declare that all data supporting the findings of this study are available within the paper and its supplementary information files. RNA-Seq data is available at NCBI GEO (accession #137151). The data that support the findings of this study are available from the corresponding author upon reasonable request.

