## Supplementary material for "Cells collectively reshape habitability of temperature by helping each other replicate": Supp. Figs and mathematical model

---

**This document contains:**

- **Supplementary Figures 1 - 17**
- **Supplementary Text (starts on Pg. 36): Detailed description of mathematical model**

#### Supplementary Figures

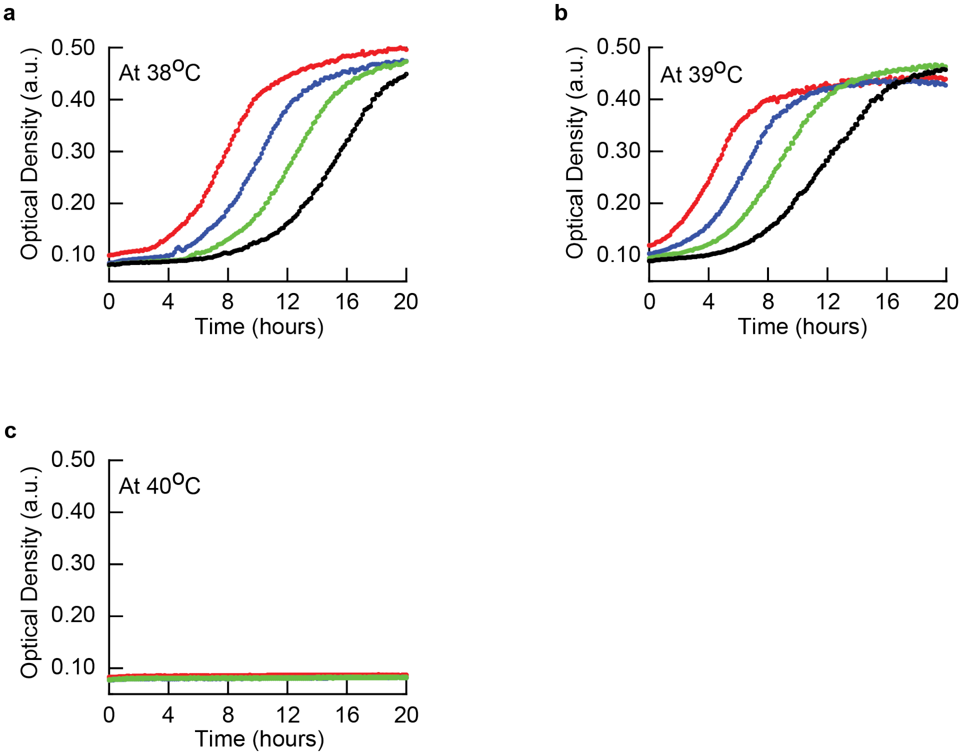


**Supplementary Fig. 1. Conventional view of temperature-dependent cell-growth for populations of wild-type yeast defines habitable and inhabitable temperatures (Related to Figure 1b). (a-c)** To obtain the conventional picture depicted in Figure 1b, we performed a laboratory-standard growth experiments in which we used a plate reader (BioTek Synergy HTX microplate plate reader, model S1LFA) to measure the Optical Density (OD) of liquid cultures of wild-type yeast cells over time (up to 20 hours shown here). OD represents the optical absorption of light at a wavelength of 600 nm and is directly proportional to the number of cells per volume, OD=0.10 corresponds to approximately $10^6$ cells/mL. The plate reader cannot detect sufficiently small ODs (i.e., OD < 0.08). We show here representative data for 38 $^{O}$C (**a**), 39 $^{O}$C (**b**), and 40 $^{O}$C (**c**). For each temperature, the different colors represent cell-populations with distinct starting ODs. To obtain these starting ODs for a given temperature, we diluted cells from a single liquid culture of cells that grew overnight at 30 $^{O}$C (also for growth experiments that appear in later figures). The starting ODs are approximately 0.10 (red), 0.05 (blue), 0.025 (green), and 0.0125 (black). Blue, green, and black curves start with ODs - obtained by serial dilutions of denser cultures - that are below the lowest OD that the plate reader can detect whereas the red populations start above it. All cultures at 38 $^{O}$C and 39 $^{O}$C reach their carrying capacities (i.e.,

ODs plateau over time in (a-b)). None of the populations grew at 40 $^{O}$C (i.e., OD remains flat over time in (c)).

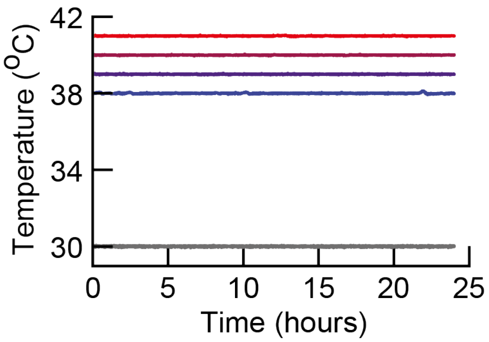


**Supplementary Fig. 2. Temperature remains stable during all our growth experiments (Related to Figure 2).** In all our growth experiments described in Figure 2 and subsequent figures (performed with liquid cultures of cells incubated in compressor-cooled, high-precision thermostatic incubators (Memmert ICPs)), the incubators stably maintained their target temperature throughout the course of our growth-experiments, with a typical standard deviation of 0.017 $^{O}$C over time (deviation measured over several days). As examples, shown here are temperatures recorded by the incubator's sensor, zoomed to 24 hours for five separate growth experiments: Starting from the top, the curves are for 41 $^{O}$C, 40 $^{O}$C, 39 $^{O}$C, 38 $^{O}$C and 30 $^{O}$C. We also verified and aligned the incubators' temperatures by using a different thermocouple device. Thus, we measured temperature values with two different thermocouple devices and the temperature remained stably constant over the course of each growth-experiment.

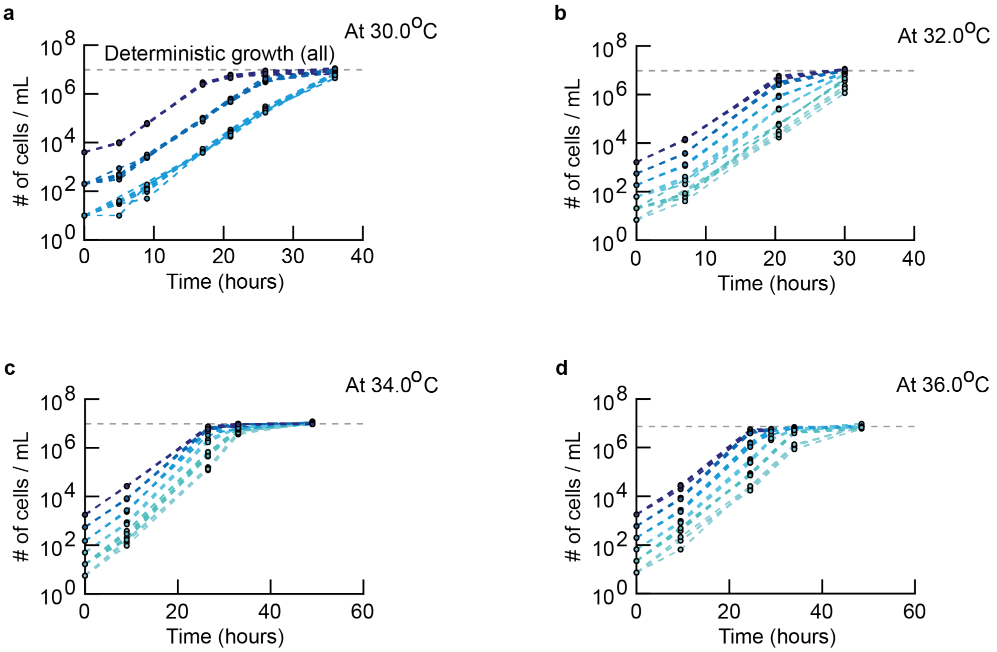


**Supplementary Fig. 3. If the temperature is at or below 36 °C, every population of wild-type cells deterministically grows regardless of the initial population-density (i.e., all populations grow to reach a carrying capacity) (Related to Figures 2a-d). (a-d)** Population density (number of cells/mL) measured over time with a flow cytometer. Each curve shows a population of wild-type cells that started with a desired initial density. The initial population-densities vary over ~1000-fold, from very dilute (~10 cells/mL) to less dilute (~1000 cells/mL). Horizontal, grey line at the top of each plot shows the carrying capacity that we estimated by averaging the density of each population after it eventually stops growing (we used only the populations with the highest initial densities for this estimate). Sample data shown for 30 °C (**a**), 32.0 °C (**b**), 34.0 °C (**c**), and 36.0 °C (**d**). Different colors represent different initial population-densities. For each color, there are eight (a) or four (b-d) replicate populations (biological replicates). For every initial population-density, every replicate population exponentially grew in an identical manner until they reached a carrying capacity. Here, "identical manner" means that all curves of the same color perfectly overlap within each panel (compare this with populations that exhibit random growths in Supplementary Fig. 4). In other words, every initial population-density, regardless of how low they were, led to a deterministic growth for temperatures at or below 36.0 °C. These results show that the no-growth and random-growth phases do not exist below 36 °C. Only deterministic growth exists for the wild-type cells at temperatures below 36 °C, consistent with the wild-type strain's phase diagram (Figure 2d).

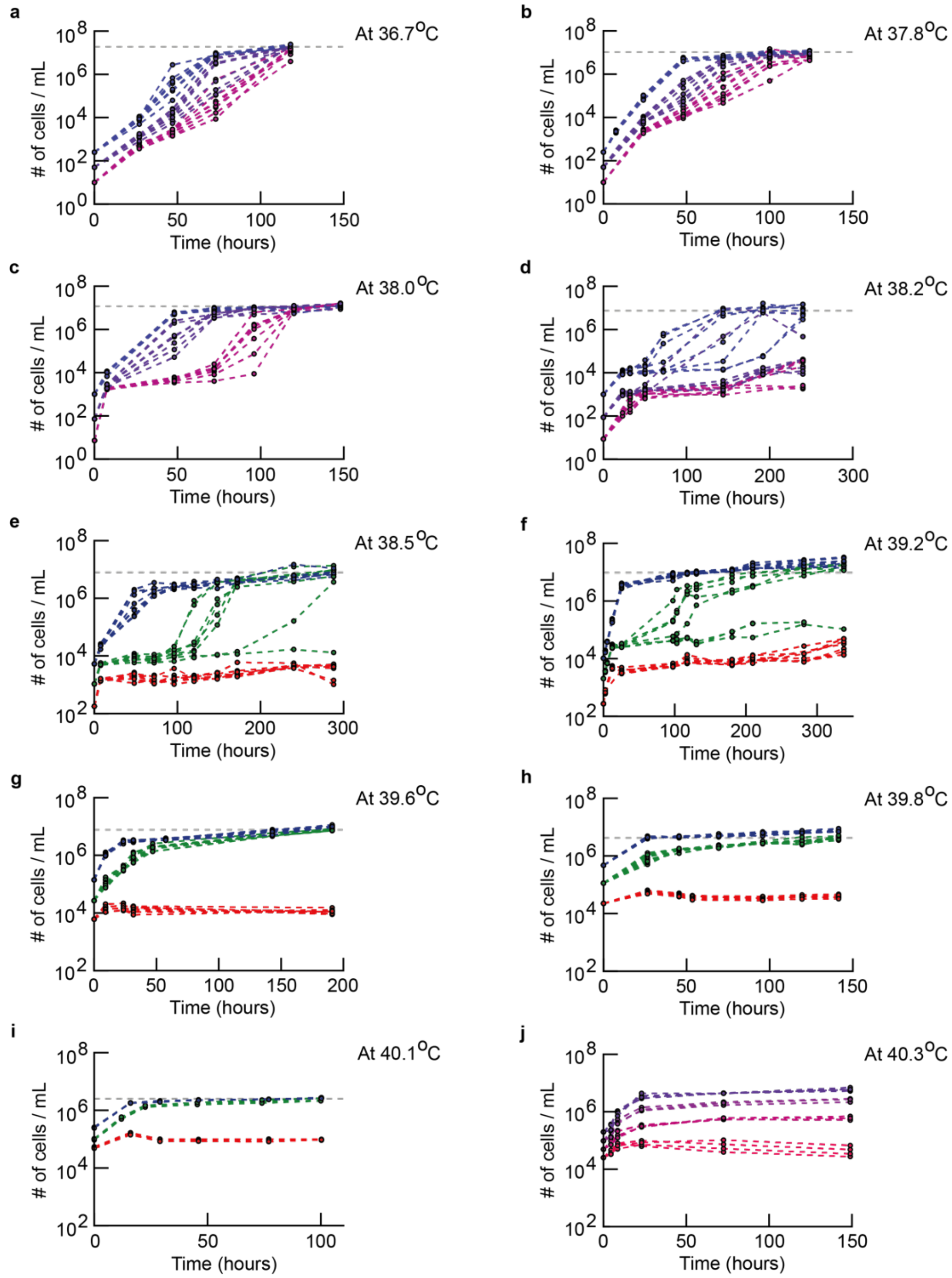


**Supplementary Fig. 4. If the temperature is above 36 °C, growth of wild-type cells depends on the initial population-density (Related to Figures 2a-e).** (**a-j**) Population density (number of cells/mL) measured over time with a flow cytometer for populations of the wild-type cells with differing initial densities. Grey line shows the carrying capacity that we estimated by averaging the final densities of the populations, which started with the lowest initial densities, that grew. Sample data shown for 36.7 °C (**a**), 37.8 °C (**b**), 38.0 C, (**c**), 38.2 °C (**d**), 38.5 °C (**e**), and 39.2 °C (**f**) - these are copied here from Figure 2a-b for completeness. Sample data shown for 39.6

$^{O}C$ (**g**), 39.8 $^{O}C$ (**h**), 40.1 $^{O}C$ (**i**), and 40.3 $^{O}C$ (**j**). Different colors represent different initial population-densities. Each color has multiple replicates. For temperatures below 40.0 $^{O}C$, there are eight replicate populations (biological replicates) for each color (i.e., they all started with the same population density). For temperatures above 40.0 $^{O}C$, there are at least three replicate populations for each color. To show multiple starting population-densities for populations having the same growth phase (e.g., random-growth phase), we used here a color scheme that is different from the one used in Figures 2a-c. Based on the growth experiments whose sample data are shown here, we constructed the phase diagram for the wild-type cells (Figure 2d). To construct the phase diagram (Figure 2d), we determined whether a given initial population-density belongs to a deterministic growth phase, or a random growth phase, or a no-growth phase from the growth-kinetics data such as the ones shown here in the following manner: An initial population-density belongs a deterministic growth phase in the phase diagram if every replicate population, all of which start with the same population density, exponentially grows over time in an identical manner (i.e., all curves of the same color overlap - collapse into a single curve - in the plots above). As an example, Supplementary Fig. 3 shows all initial population-densities leading to a deterministic growth at temperatures below 36 $^{O}C$. An initial population-density belongs to the no-growth phase in the phase diagram if none of the replicate populations grow after an initial, transient growth that typically lasts at most ~10 hours due to the effect of the cells having just been transferred from a 30 $^{O}C$ to their new temperature. As an example, in (g), the lowest initial population-density (red curves) belongs to the no-growth phase. An initial population-density belongs to a random-growth phase in the phase diagram (Figure 2d) if the curves of the same color - representing replicate populations - do not overlap or, in the starkest cases, when some replicate populations do grow while others do not. Here, the curves of the same color, representing replicate populations that grow, do not overlap due to the each population growing at distinct rates or starting to grow - after a stasis - at different times after the transient growths stop, causing these populations to reach a carrying capacity at vastly different times (i.e., different by tens to hundreds of hours) despite all replica populations having the same initial density. As an example, an intermediate initial population-density in (e) - represented by green curves - leads to random growths. Finally, we determined the phase boundaries in the phase diagram (Figure 2d) as follows: We drew the boundary that divides the deterministic-growth and random-growth phases by connecting the data points that represent the lowest measured initial-population-density that yielded a deterministic growth for each temperature. We drew the boundary that divides the random-growth and no-growth phases by connecting the data points that represent the highest measured initial-population-densities that yielded a no-growth phase for each temperature. Finally, we determined the temperature above which a population-level growth is no longer possible by determining the lowest temperature at which populations that start with different densities always reach different final densities when they stop growing (i.e., populations never grow to the carrying capacity whereby one or more essential nutrients has been depleted). As an example, at 40.3 $^{O}$C (j), populations with different initial densities never reach the same density when they stop growing.

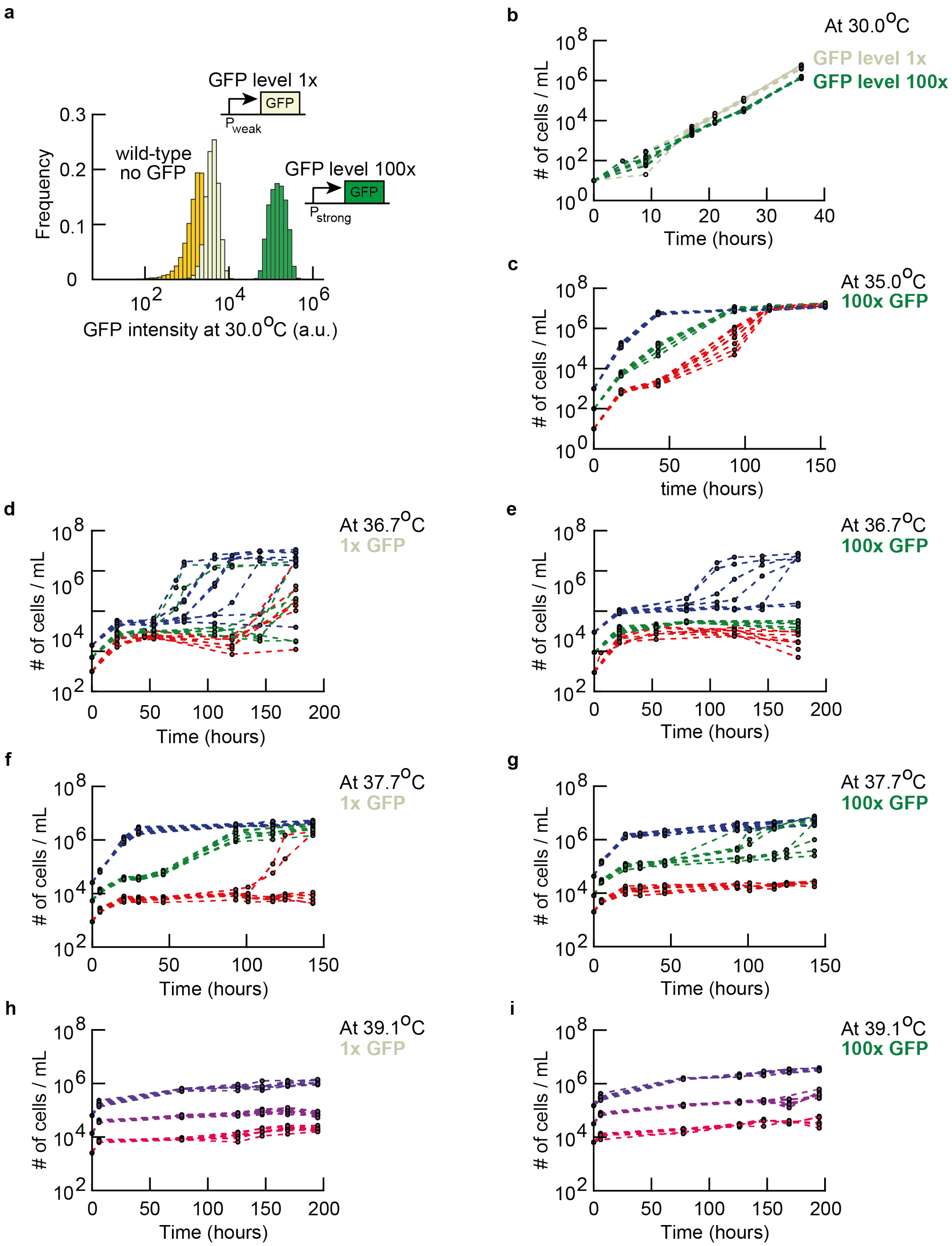


**Supplementary Fig. 5. Cost of expressing a spurious gene (*GFP*) alters how an initial population-density affects whether a population grows or not at high temperatures (Related to Figure 2f).** Characterization of the two yeast strains ("1x-GFP" and "100x-GFP" strains) that constitutively express GFP, which we used to construct the phase diagrams in Figure 2f. **(a)** A histogram of GFP-expression levels of the two engineered yeasts and the wild-type strain as measured by a flow cytometer. The 1x-GFP cells have a higher mean-fluorescence than the wild-type cells whereas the 100x-GFP cells have approximately 100-fold higher mean-fluorescence than the 1x-GFP cells. (**b-i**) As with the wild-type strain (Figure S3-4), we performed growth experiments in which we used a flow cytometer to measure the population density (number of cells/mL) for the 1x-GFP and 100x-GFP strains. Sample data shown for 30 $^{O}$C (**b,** 1x-GFP strain in grey and 100x-GFP strain in green), 35.0 $^{O}$C (**c,** 100x GFP strain), 36.7 $^{O}$C (**d,** 1x-GFP strain; **e,** 100x-GFP strain), 37.7 $^{O}$C (**f,** 1x-GFP strain; **g,** 100x-GFP strain), and 39.1 $^{O}$C (**h,** 1x-GFP strain; **i,** 100x-GFP strain). Each color shows multiple populations that all start with the same initial density. There are typically eight replicate populations (biological replicates) for each color. To show multiple starting population-densities for each of the growth phases (e.g., random-growth phase), we used here a color scheme that is different from the one that we used in Figures 2a-c. To distinguish deterministic, random, and no-growth phases for the 1x-GFP and 100x-GFP strains, we used criteria that are similar to the ones that used for the wild-type strain (Figure 2d) which we described in the caption for Supplementary Fig. 4. Here, for a given temperature, we classified an initial population-density as belonging to the deterministic-growth phase if at least six out eight replicate populations (biological replicates) that started with this density exponentially grew (note that for the wild-type cells, all eight out of eight replicate populations must have exponentially grown for the initial population-density to be classified as yielding a deterministic growth). Conversely, for a given temperature, we classified an initial population-density as belonging to the no-growth phase if six out of eight populations with the same initial population-density did not grow (for the wild-type strain, all eight populations had to not grow). These slight differences in the definitions of the phases between the wild-type and the GFP-expressing strains do not qualitatively change the main features of the phase diagrams (Figure 2f). In short, the main conclusion - that expressing more of a spurious gene (*GFP*) means that, at a given temperature, a population must start with a higher density of cells than a population of cells with a lower GFP-expression in order to achieve a deterministic growth - is unaffected by the slight difference in the definition of the phases.

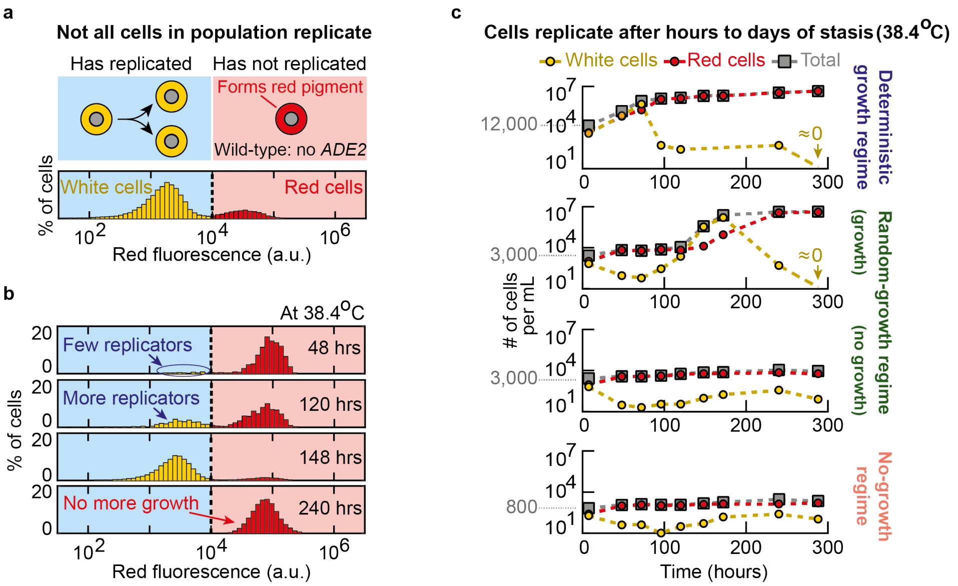


**Supplementary Fig. 6. Few cells stochastically and transiently replicate within populations that are in either a no-growth or a random-growth phase (Related to Figures 2a-d).** (**a**) Wild-type strain, with a defective *ADE2* gene, produces red pigments as by-products of the not-fully repressed adenine-biosynthesis. Red cells have not been dividing for some time - they appear red because they accumulated the red pigments rather than diluting them through cell divisions. Non-red ("white") cells have been dividing. Histogram shows percentages of red and white cells in a population determined by a flow cytometer's red-fluorescence detector that quantified redness of individual cells. (**b**) Percentage of white and red cells over time measured with the flow cytometer for a population of wild-type yeasts. Time shows hours of incubation in 38.4 °C. These histograms show example time courses for a population that grew at a high temperature. (**c**) Numbers of white and red cells in a population per mL, at various times for three different growth regimes indicated by the phase diagram (Figure 2d). Random-growth regime shows two replicate populations - one growing (second row) and one not growing (third row).

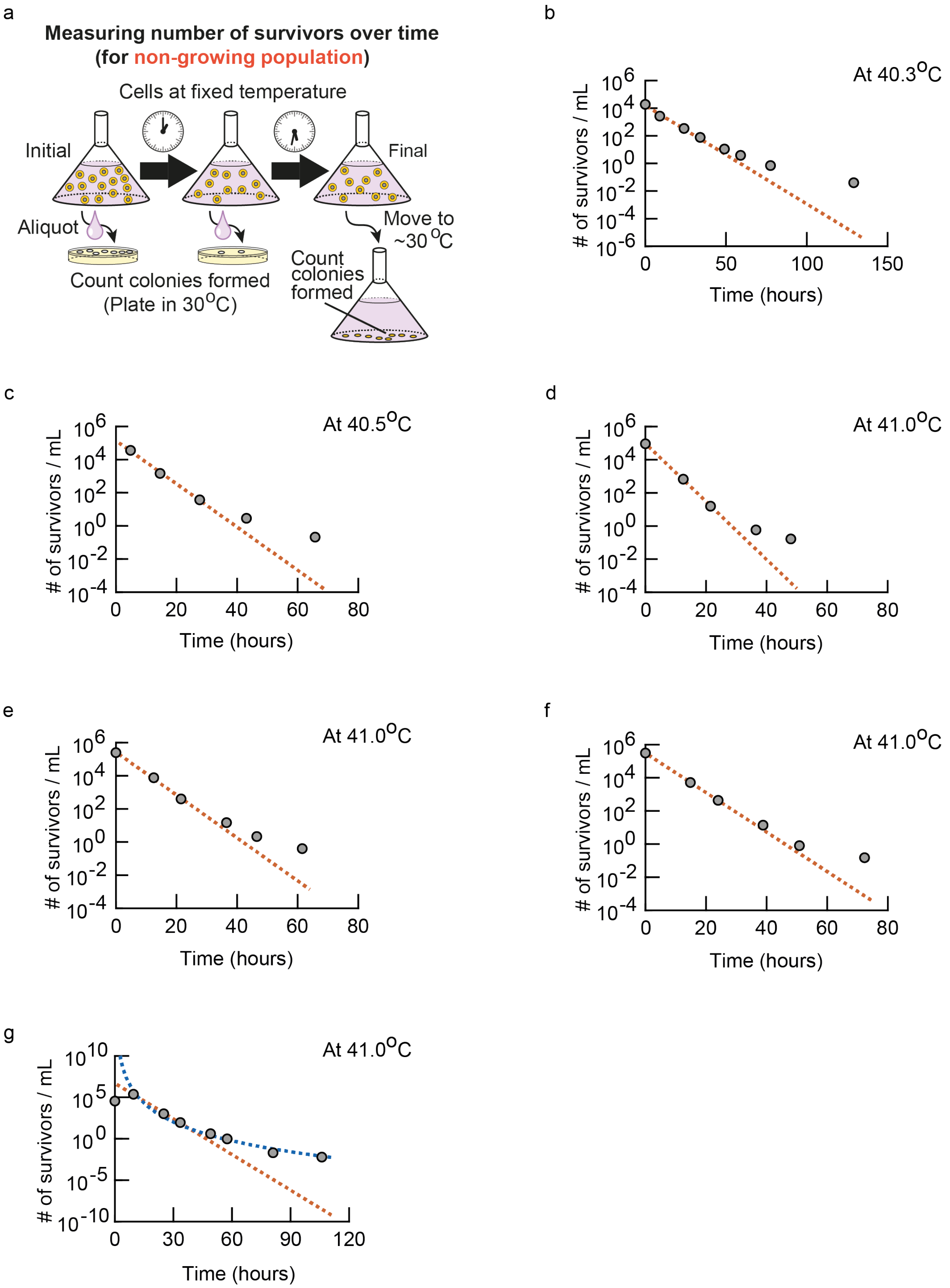


**Supplementary Fig. 7. Unconventional deaths at high temperatures: number of survivors at high temperatures decreases over time as a heavy-tailed function and deviates by orders of magnitude from the conventionally expected number of survivors (Related to Figures 3a-b).** (**a**) Schematic description of how we measured the number of surviving wild-type cells, over time, in a population that was incubated at a high temperature and could not grow (because it was in the no-growth phase according to the phase diagram (Figure 2d)). In short, at various times, we took out an aliquot from a liquid culture of non-growing population that was kept at a high temperature. We then serially diluted this aliquot by known amounts - this was to ensure that we could have countable numbers of colonies if there were any survivors in the aliquot - and spread the droplets containing the serially diluted cells on an agar pad which we incubated at 30 $^{O}$C. After a few days, we counted the number of colonies that formed (colony forming units) on the agar pad at 30 $^{O}$C. By counting the number of colony-forming units and knowing the dilutions and volumes of the aliquots that we took out at various times, we determined the "number of survivors/mL" that we plotted in (b-g) and Figures 3a-b. For the last several time points in (b-g) and Figures 3a-b, aside from counting the number of colony forming units, we double checked our results by using a complementary method to count the number of survivors per volume: taking an aliquot whose volume is only a fraction of the total volume of the liquid culture would have yielded very few colonies on the agar pad, since there was typically less than one survivor per mL in the liquid cultures. Thus, we took out an appropriate volume (typically tens of mL) from the flask that contained the entire liquid culture which was incubated at a high temperature, transferred it to an Erlenmeyer flask, and then left the flask at ~30 $^{O}$C for several days without shaking it so that all the surviving cells in that liquid that we took out settled down to the bottom of the flask and formed colonies. To measure the last time points in (b-g) and Figure 3a-b, we moved the flask that contained the entire remaining liquid culture (i.e., the entire population) from a high temperature to ~30 $^{O}$C to ensure that we counted all remaining survivors using the "settling-to-the-bottom" method. Both methods - directly counting the colonies formed on agar after spreading a serially diluted aliquot of the liquid culture and counting the colonies formed by surviving cells that settled down to the bottom of a flask - yielded the same results. (**b-g**) We used the method in (a) to measure the number of surviving wild-type cells per mL at 40.3 $^{O}$C (**b**), 40.5 $^{O}$C (**c**), and 41.0 $^{O}$C (**d-g**). In (b-f), brown dashed lines represent an exponentially decaying function that we fitted to the first three time points (i.e., data points for the first day of incubation at a high temperature). For (g), the brown dashed lines represent an exponentially decaying function that we fitted to the data points that lie within 10 - 50 hours. The blue dashed curve is a power-law function fitted to the same data points as the ones that we used to fit the exponentially decaying functions. In (b-g), we see that the data points vastly deviate from the brown dashed line (i.e., the final time point deviates by at least ~$10^4$ cells/mL). Thus, contrary to the conventional view of cell death at high temperatures (*4*), the number of survivors does not exponentially decrease over time (i.e., cell death is neither autonomous nor fixed by a single (exponential) rate constant). Instead, it decreases over time as a heavy-tailed function as seen here.

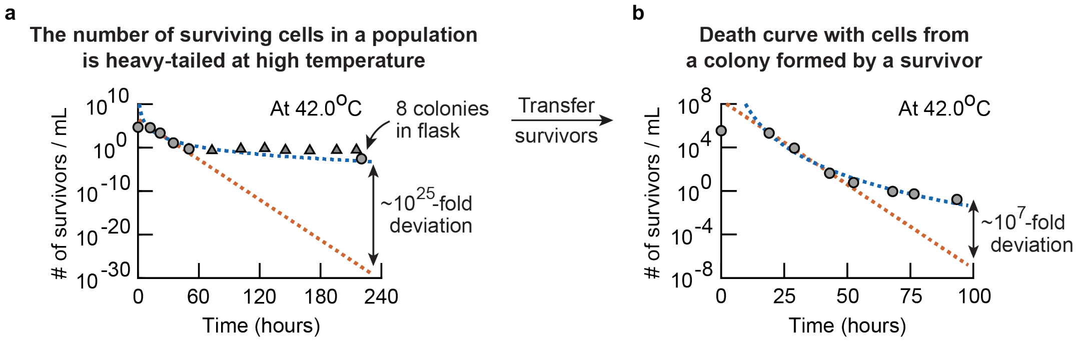


**Supplementary Fig. 8. Number of survivors decreasing over time as a heavy-tailed function is not due to heat-tolerant mutants or persistor-like cells existing within a population (Related to Figures 3a-b). (a-b)** We used the method described in Supplementary Fig. 7a to measure the number of survivors per unit volume (per mL) at different times for a population of wild-type cells kept in 42.0 $^{O}$C. At this temperature, the population is in the no-growth phase in the phase diagram (Figure 2d) so the number of viable cells cannot increase over time. The brown dashed lines represent an exponentially decaying function that we fitted to the data points that lie within 10 - 50 hours. The blue dashed curve is a power-law function that we fitted to the same data points as the exponentially decaying function. Circles show measured number of survivors/mL. (**a**) The number of surviving wild-type cells. Triangles are also from measurements and are overestimates. Specifically, by "overestimates", we mean that the aliquots taken from the liquid culture at 42 $^{O}$C did not yield any colonies on agar at 30 $^{O}$C, which, in turn, means that we should have taken a larger aliquot from the liquid culture in order to find at least one survivor if there were any survivors in the liquid culture. Hence, if there were any survivors in the liquid culture, there could not have been more than values represented by the triangles (i.e., the concentration of survivors must be more dilute than the value represented by the triangles). We directly observed eight colonies formed at the last timepoint (~220 hours - see the last circle). (**b**) We took one of these eight colonies - progenies of the survivors from the last timepoint in (a) - and used the cells from this colony to repeat the experiment shown in (a) (i.e., see how these cells survive at the same high temperature (42.0 $^{O}$C)). The number of survivors in this new experiment decreased over time as a heavy-tailed function, just as in (a). This result eliminates the possibility that the survivors of the high temperature (42 $^{O}$C) in the first experiment (a) are either heat-tolerant mutants or persistor-like cells that behave similarly to persistors of antibiotic treatments. Indeed, if the survivors of the first experiment (the last time point in (a)) were persistor-like cells or heat-tolerant mutants, then the number of survivors/mL in (b) should have decreased over time as a single, slowly decaying exponential function instead of as a heavy-tailed function. To see this, note that if the population in (a) initially contained a subpopulation of persistor-like cells or heat-tolerant mutants, then the cells of this subpopulation would have died at a slower rate than the rest of the population. This, in turn, would mean that the number of surviving cells/mL would decrease over time as a biphasic function - a sum of a slowly decaying exponential function (representing the subpopulation of heat-tolerant mutants or persistor-like cells) and a more rapidly decaying exponential function (all the other cells) - which would superficially appear to resemble a heavy-tailed function shown in (a). This is in fact why one can wonder whether the survivors in (a) are heat-tolerant mutants or persistors in the first place. But by design, the population in (b) is pure - it does not contain any subpopulation of cells that are different from the rest of the population. Thus, if the survivors at the last time point in (a) are heat-tolerant mutants or persistor-like cells, then the population in (b) must start as a pure population of heat-tolerant mutants or persistor-like cells. It then follows that the number of survivors/mL in (b) should decrease as a single, slowly decaying exponential function, which it does not. Rather, it decreases over time as a heavy-tailed function. Thus, this experiment establishes that the survivors of a high temperature are not heat-tolerant mutants or persistor-like cells.

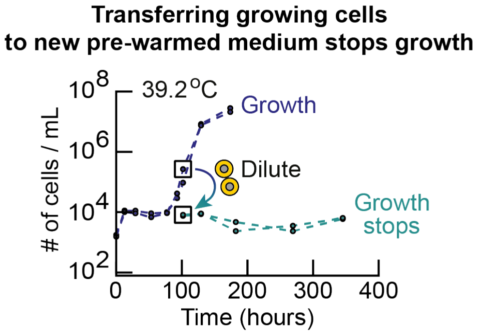


**Supplementary Fig. 9. Transferring cells growing at ~39 °C to a fresh medium that was pre-warmed to 39 °C before the transfer stops the cells' growth.** All experiments done at 39.2 °C. Two overlapping blue curves show two populations of wild-type cells that started with the same initial density. At this initial population-density, these two populations are in the random-growth phase at 39.2 °C. After ~100 hours of incubation at 39.2 °C, we took some of the cells in these populations, which were growing in mid-log phase (marked by the boxed blue data points), and transferred them into a fresh minimal medium that we pre-warmed to 39.2 °C. The transfer was such that the newly created population, in the fresh medium, started with the same number of cells/mL as the population density that the original populations had after their transient growth and before they started to grow (~10,000 cells/mL as shown). A population that starts with 10,000 cells/mL at this temperature (39.2 °C) would be in the deterministic-growth phase according to the phase diagram (Figure 2d). We incubated the newly created cultures at the same temperature as the original population (see "Growth experiments" in the Methods section) and measured their population densities over time (green curves show two replicate populations). Both populations did not grow at all (see flat green curves), indicating that cells that were growing at the high temperature did not continue to grow when transferred to fresh media at the same high temperature. The fact that these populations of transferred cells (green curves) did not grow does not contradict the phase diagram (Figure 2d) since, for constructing the phase diagram, we transferred cells from 30 °C to a new (higher) temperature whereas for the experiment described in this supplementary figure, we transferred cells from 39.2 °C to 39.2 °C.

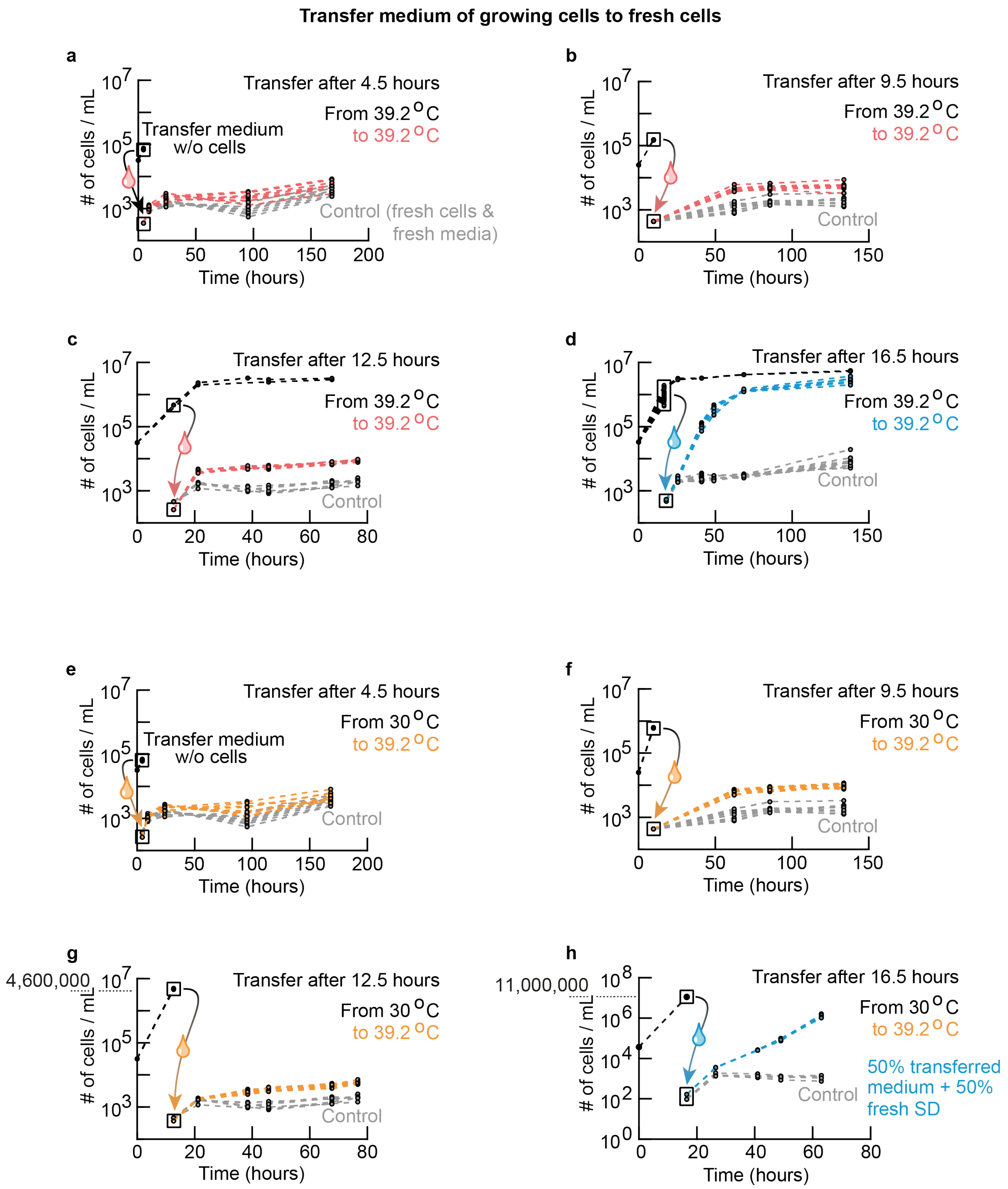


**Supplementary Fig. 10. Density-dependent growths of cells at high temperatures are due to the cells collectively altering their shared extracellular environment (Related to Figures 4b-c).** (**a-d**) All black curves all represent a population of wildtype cells that started with 30,000 cells/mL and was deterministically growing at 39.2 °C. We let such a population grew for 4.5 hours (**a**), 9.5 hours (**b**), 12.5 hours (**c**), or 16.5 hours (**d**) before we flowed the liquid cultures of these growing cells through filter papers that had 200-nm-diameter pores. This resulted in a complete, physical separation of the cells from their liquid growth media. We confirmed the complete separation by flowing the filtered media through a flow cytometer: we detected no cells in them at all. We then transplanted fresh cells into each of these filtered media. This created new populations, all starting with 400 cells/mL, which we also incubated at 39.2 $^{O}$C. We then measured the densities of these newly created populations over time (red curves for (a-c) and blue curves for (d)). As a control, we also incubated fresh cells in fresh media - rather than in one of the filtered media - so that the resulting population started with the same density - 400 cells/mL - as the other newly created populations. We then measured the control population's density over time (grey curves in (a-d)). For each color in every panel (a-d), there are at least six replicate populations shown. Red curves in (a-c) show no appreciable growth beyond transient growths while the blue curves in (d) show deterministic growths that reach the carrying capacity. These results (a-d) show that growing cells at a high temperature (e.g., 39.2 $^{O}$C) gradually alter their extracellular growth medium - for example, by potentially secreting a factor - in such a way that the medium can induce growth of populations that cannot grow by themselves (without inheriting the changed medium) because their initial densities are too low (e.g., 400 cells/mL). (**e-h**) We used the same protocol as in (a-d) except that this time, we grew the wildtype cells at 30.0 $^{O}$C instead of at 39.2 $^{O}$C before taking away their growth media for creating new populations, which we did incubate at 39.2 $^{O}$C. Specifically, we grew the wildtype cells - starting again at 30,000 cells/mL as in (a-d) - for 4.5 hours (**e**), 9.5 hours (**f**), 12.5 hours (**g**), or 16.5 hours (**h**) before taking away their liquid media and transplanting fresh wildtype cells into them by using the same filtration method as in (a-d). We incubated the newly created populations - each starting with 400 cells/mL - at 39.2 $^{O}$C as in (a-d). Orange curves (e-g) show these populations' densities over time at 39.2 $^{O}$C. Grey curves (e-g) show control populations which are identical to the control populations in (a-d). Beyond the initial, transient growths, none of the orange curves (e-g) show any sustained exponential growths (i.e., none reach the carrying capacity). The results in (e-g) show that, unlike the media taken from the cells that were growing at a high temperature (39.2 $^{O}$C), the media taken from cells growing at the conventional temperature (30 $^{O}$C) does not contain the right factors for inducing growth of populations at a high temperature (39.2 $^{O}$C). (**h**) Population that was incubated at 30 $^{O}$C was in a stationary phase after 16.5 hours of incubation, as shown here, after a log-phase growth that depleted essential nutrients which caused the population to undergo a diauxic shift. We took out this stationary population's growth medium and transplanted fresh cells into it, and then incubated this newly created population at 39.2 $^{O}$C. Blue curves show this population - all replicate populations - growing at 39.2 $^{O}$C and nearly reaching the carrying capacity. These results (e-h) show that some factors that yeasts secrete during a stationary phase at 30 $^{O}$C after a diauxic shift - and perhaps during a diauxic shift - can induce population growths at high temperatures (e.g., at 39.2 $^{O}$C).

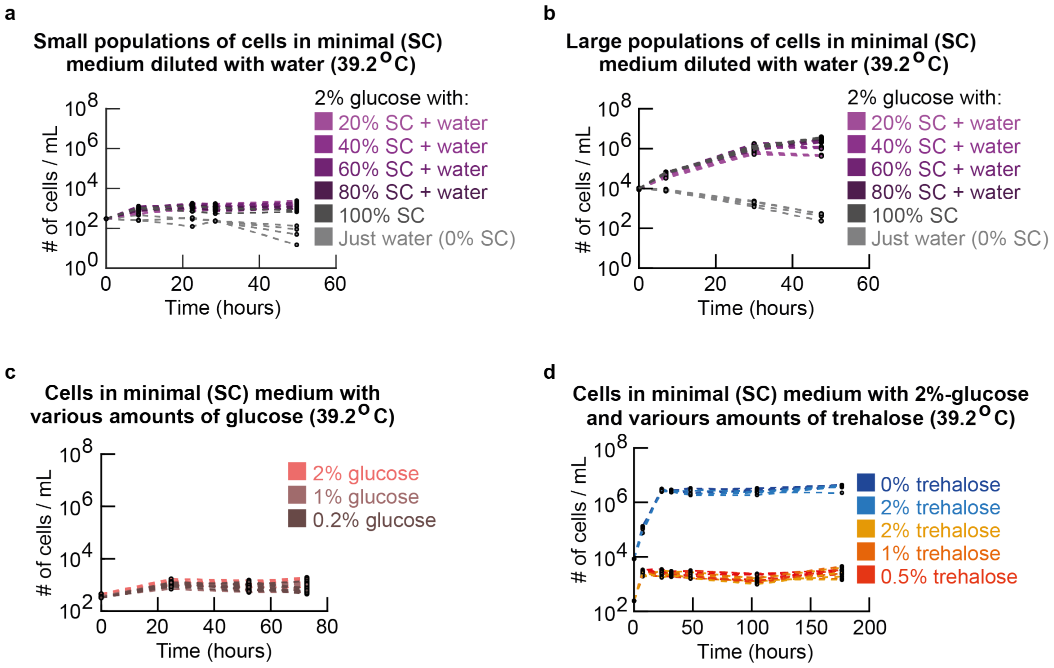


**Supplementary Fig. 11. Population-density dependent growths at high temperatures are not due to cells depleting essential nutrients from the extracellular medium (Related to Figure 4c). (a-b**) Minimal media (called "SC media") contains all essential amino acids and nitrogenous bases. Wildtype cells were incubated in SC media that we diluted with water by various amounts as indicated (note: "100% SC" means no dilution, "50% SC" means that we used water to dilute all contents of SC by half, and so on). We supplemented all media, regardless of by how much they were diluted, with a saturating concentration (2%) of glucose. Graphs show how population densities change for cells incubated in a 20%-SC medium (i.e., 20% SC + 80% water, supplemented with a 2% glucose), 40%-, 60%-, 80%-, and a 100%-SC medium. Different colors represent different dilutions of SC. Light grey curves show populations incubated in just water with a 2%-glucose (i.e., no SC). (**a**) All curves start at ~400 cells/mL. None of the populations grew. (**b**) All curves start at 10.000 cells/mL. All populations, except for those without any glucose (100% SC) or without any essential amino acids (0% SC), grow deterministically until they reach their respective carrying capacities. Thus, the 20% to 100% SC all contain sufficient nutrients for a population to grow. Together, (a) and (b) show that population growths are not caused by depletions of some extracellular components in the SC-medium (i.e., these results suggest that cells secrete some factors that induce population growths). (**c**) Complementary to (a), wildtype cells were incubated in minimal media (100% SC -media) with various concentrations of glucose. Shown here are cells incubated in SC + 2% glucose, SC + 1% glucose, and SC + 0.2% glucose. None of these populations grew. Thus, population-density dependent growths that we observed in our study are not due to glucose depletions. (a-c) together establish that it is not a depletion of any of the nutrients in the media that cause the observed population-density dependent behaviors. (**d**) Wildtype cells incubated in minimal media supplemented with various concentrations of trehalose at two different initial population densities (one that is too low for population-level growth (400 cells/mL) and one that is sufficiently high for population-level growth (10,000 cells/mL)). These data show that trehalose neither inhibits (at 10,000 cells/mL) nor induces (at 400 cells/mL) population growths. Trehalose is a common antioxidant. These results show that trehalose plays no role in aiding or preventing populations growths at high temperatures.

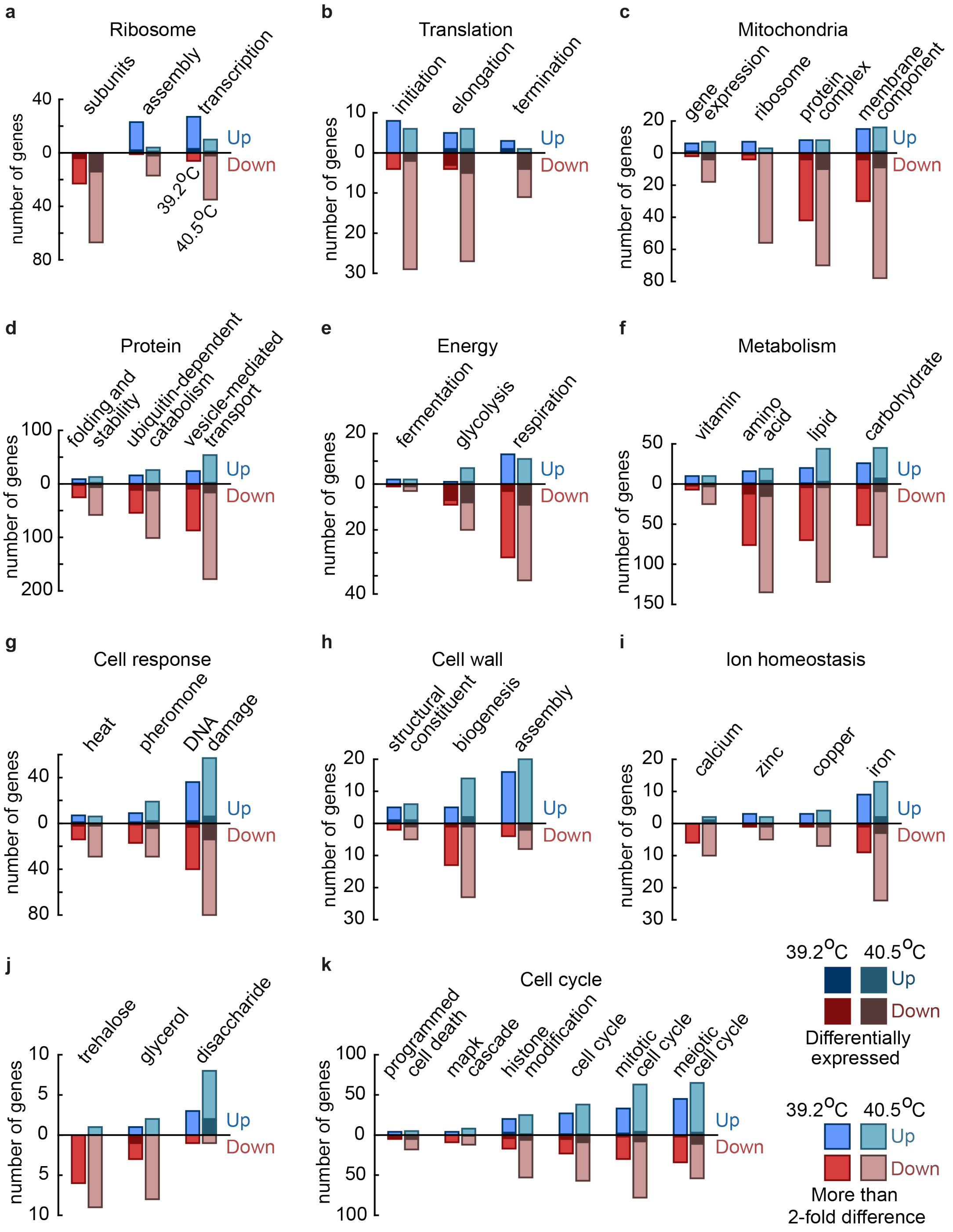


**Supplementary Fig. 12. Transcriptome (RNA-seq) analysis reveals effects of high temperatures on gene expression in deterministic-growth and no-growth phases (Related to Figures 4d-f).** Transcriptome (RNA-seq) analysis of wild-type yeast cells in mid-log phase growth at 39.2 $^{O}$C after 75 hours and 100 hours of incubation (initial population-density: 11,000 cells/mL) and in no-growth-phase at 40.5 $^{O}$C after 72 hours of incubation (initial population-density: 48,000 cells/mL). Genes were categorized based on the Gene Ontology (GO)

annotations. Shown here are the number of genes that are upregulated (blue) and downregulated (red) for the respective categories. Genes are classified as up- or down-regulated relative to their expression levels when they grow (always deterministically) at 30 $^{O}$C. For the analysis, we averaged the expression levels of three (at 39.2 $^{O}$C and 30 $^{O}$C) or two (at 40.5 $^{O}$C) biological replicates. The gene counts only include genes whose expression level differed by at least a 2-fold from their expression levels at 30 $^{O}$C (shown in light colors) and differentially expressed genes (shown in dark colors). (**a**) For genes associated with the ribosome: Notably, ribosomal protein subunits were downregulated. Ribosome assembly, polymerases I and III and transcription of rRNA were upregulated for deterministically growing yeasts at 39.2 $^{O}$C while they were downregulated for yeasts in the no-growth phase at 40.5 $^{O}$C. (**b-c**) For genes associated with translation (**b**) and the mitochondrial genes (**c**): Almost all differentially expressed genes were downregulated. The ribosomal genes of mitochondria were upregulated for deterministically growing cells at 39.2 $^{O}$C and downregulated for the no-growth-phase cells at 40.5 $^{O}$C compared to their expression levels at 30 $^{O}$C. (**d-f**) For genes associated with protein processing (**d**) and genes associated with the central carbon metabolism (**e**) and other metabolic activity (**f**), many of which are significantly differentially expressed: Most notably, genes of the glycolysis and respiration were downregulated at the high temperatures. (**g-h**) For cellular responses to heat and DNA damage (**g**) and genes associated with the cell wall (**h**): Strikingly, cell wall assembly was upregulated for both the deterministically growing cells and the no-growth-phase cells at high temperatures relative to their expression levels at 30 $^{O}$C. (**i-j**) For genes associated with ion homeostasis (**I**), and other carbohydrates (**j**): Most genes involved in the turnover of trehalose (a metabolite involved in thermotolerance) were downregulated compared to their expression levels at 30 $^{O}$C. Genes associated with metabolism of disaccharides such as maltose were upregulated, even though the minimal growth medium lacked disaccharides and our wild-type strain is unable to grow on maltose. (**k)** Genes associated with the cell cycle. (**a-k**) Overall summary: Our transcriptome analysis revealed that global gene-expression levels were predominantly downregulated for both deterministically growing cells (at 39.2 $^{O}$C) and no-growth-phase cells (at 40.5 $^{O}$C) compared to the expression levels at 30 $^{O}$C. Moreover, we found that this global downregulation was more pronounced for the no-growth-phase cells at 40.5 $^{O}$C than for the growing cells at 39.2 $^{O}$C. Interestingly, the central carbon metabolism was downregulated for the no-growth-phase and deterministically growing cells and the ribosomal subunits were also downregulated, even for the deterministically growing cells at 39.2 $^{O}$C. Furthermore, genes involved in the ribosomal transcription and assembly, the mitochondrial ribosome, and translation termination were upregulated for the deterministically growing cells at 39.2 $^{O}$C and downregulated for the no-growth-phase cells at 40.5 $^{O}$C.

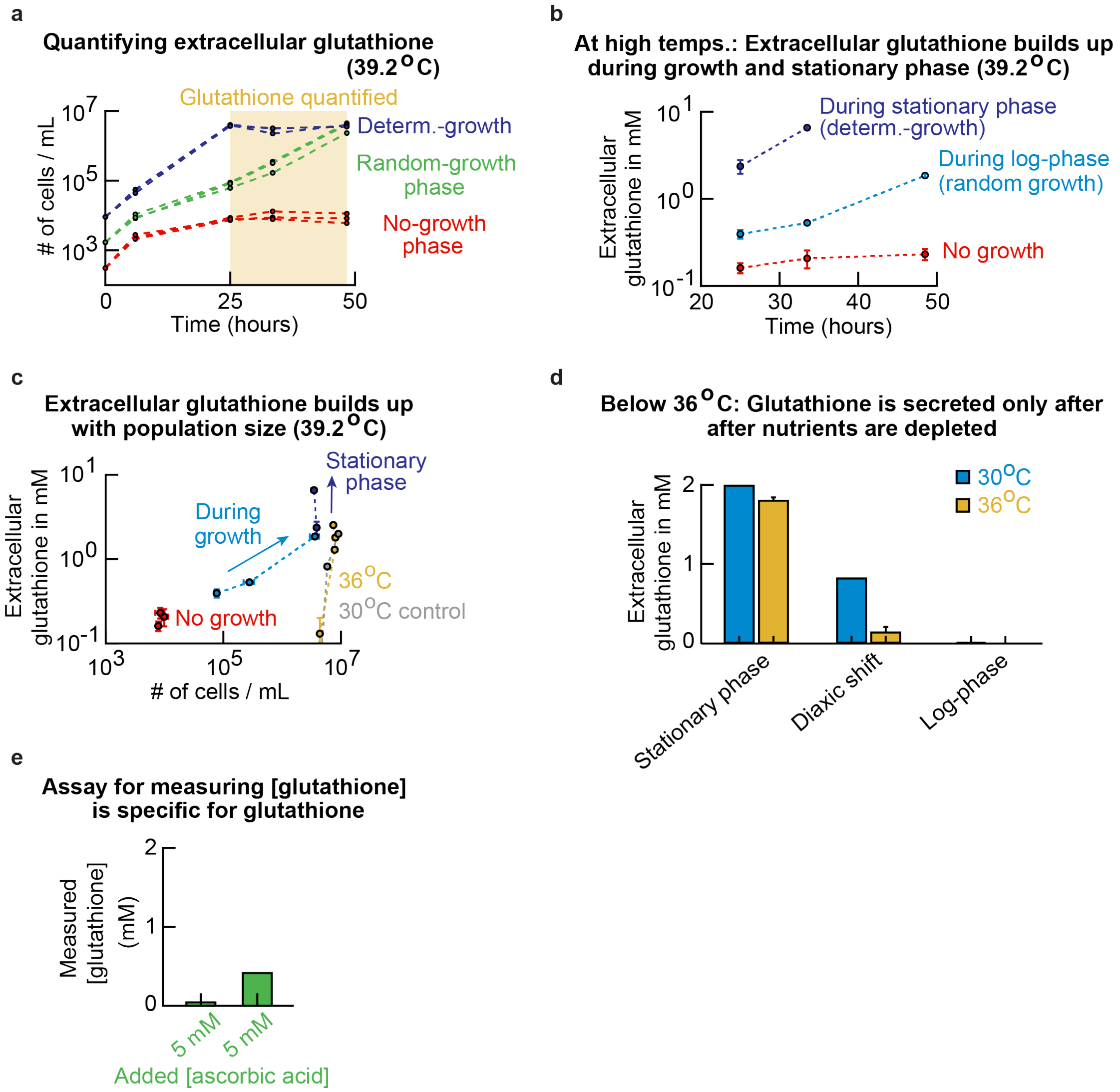


**Supplementary Fig. 13. At high temperatures, cells in the random-growth phase and deterministic-growth phase populations secrete glutathione, while the cells are in log-phase growth. These cells also secrete glutathione while they are in stationary phase (Related to Figure 4e).** (**a**) Populations with different starting densities at 39.2 °C ($n$ = 3). (**b-c**) For each of these populations, we measured their extracellular glutathione concentration after 25 hours, 33 hours, and 48 hours of incubation at 39.2 °C (i.e., for the time points in the yellow region in (a)) ($n$ = 3). (**b**) To quantify the extracellular glutathione concentration, we separated the cells from their medium by using a filter that removes the cells (VWR: filter with 0.45-μm pores and a cellulose-acetate membrane). To ensure and verify that there were no cells left behind in the filtered media, we flowed the filtered media through a flow cytometer. The flow cytometer did not detect any cells in the filtered medium. We measured the glutathione concentration in the filtered medium that we took from each population shown in (a) with a commercial assay kit (see Methods). The extracellular glutathione concentration remained constant over time at a very low level for the no-growth-phase populations (red curve in (b)). The extracellular glutathione concentration increased over time during the log-phase growths (light blue curve in (b)). The extracellular glutathione concentration kept increasing over time after a population had stopped growing because it reached a stationary phase (due to reaching a carrying capacity) (dark blue curve in (b)). To check whether most of the extracellular glutathione was in the oxidized or the reduced form, we also determined the concentration of oxidized glutathione in the filtered media taken from a population that was incubated for 48 hours at 39.2 $^{O}$C. For populations growing in log-phase (light blue in (b)), we found that most of the extracellular glutathione was in the reduced form (77% $\pm$ 3% (s.e.m., $n$ = 3), approximately 3:1 ratio of reduced-to-oxidized form). For the non-growing populations in the no-growth phase (red in (b)), we found that most of the extracellular glutathione was in the oxidized form (25% $\pm$ 24%, approximately 1:3 ratio of reduced-to-oxidized form). Hence, growing populations maintain an extracellular environment with more reduced glutathione than oxidized glutathione (note that the reduced form of glutathione, not the oxidized form, is able to remove reactive oxygen species through redox reactions). (**c**) By combining the results of (a) and (b), we determined the extracellular glutathione concentration as a function of the population density. This plot shows that the no-growth-phase populations maintain, over time, a nearly constant population density as well as a nearly constant extracellular glutathione concentration. While a population is growing in log-phase, the extracellular glutathione concentration keeps increasing while the population density is increasing over time. The glutathione concentration continues to increase after the population has reached the carrying capacity and stops growing (i.e., during stationary phase). Also shown is the concentration of the extracellular glutathione for populations at 30 $^{O}$C (grey: glutathione only detectable after the population enters a stationary phase - no secretion of glutathione during log-phase growth). Also shown is the concentration of the extracellular glutathione for populations at 36 $^{O}$C (orange: same conclusion as in 30 $^{O}$C). (**d**) As a control, we measured the concentration of the extracellular glutathione for populations incubated at 30 $^{O}$C and 36 $^{O}$C. We did not measure any extracellular glutathione for populations that were growing at log-phase at these temperatures (unlike in the case of higher temperatures such as 39.2 $^{O}$C - see (c)). But as soon as these cells depleted glucose, they started to secrete glutathione, resulting in the glutathione accumulating in the extracellular medium over time while the cells were in stationary phase at 30 $^{O}$C and 36 $^{O}$C. This observation matches the fact that the media that we transferred from a population that was in stationary phase at 30 $^{O}$C (16 hours after incubation in 30 $^{O}$C) induced population growth at 39.2 $^{O}$C (Figure S10h). Moreover, these results show that cells only secrete glutathione during log-phase growth at temperatures above 36 $^{O}$C, consistent with our observation that the wildtype cells' growths depend on the initial population-density only for temperatures above 36 $^{O}$C (Supplementary Figs. 3-4). (**e**) Testing the specificity of the commercial glutathione assay kit used in (b-d). We added either a low or beyond-saturating (physiologically unrealistic) concentration of ascorbic acid into minimal media and then subjected these ascorbic-acid containing media to the glutathione assay kit. Note that ascorbic acid is also an antioxidant but one that the budding yeast does not produce. For physiological concentrations of ascorbic acid (e.g., 5 μM shown), the glutathione assay kit did not show any readings (i.e., it did not falsely report that glutathione was present). It only reported a false signal (i.e., presence of glutathione) for the non-physiological ascorbic-acid concentration of 5 mM - the kit falsely reported ~0.4 μM of non-existent glutathione. Note that 0.4 μM is much lower than the >1 μM glutathione that we observed in the filtered media of cultures grown at the high temperatures. Thus, for our purpose, we can say that the glutathione assay kit is specific for detecting glutathione with negligible false positive readings.

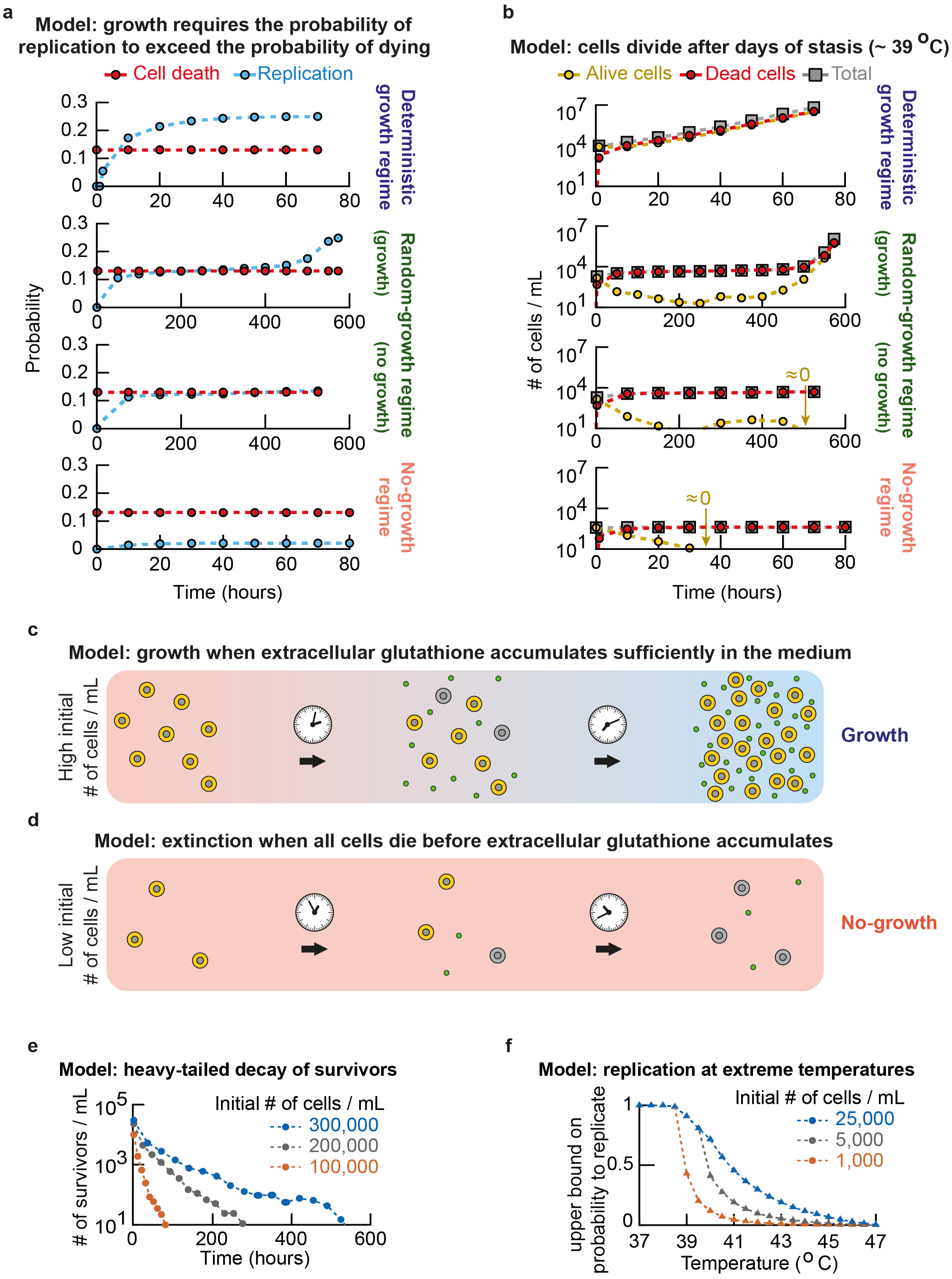


**Supplementary Fig. 14. Schematics of the mathematical model that recapitulates all the main features of our data (Related to Figure 5).** (**a**) The probability of replicating and the probability of dying as a function of time for the deterministic growth (top row), random growth

(2nd row), random no-growth (3rd row) and no-growth phases (4th row). Here, we used the same, fixed set of parameters as the ones that we used to reproduce all the other main experimental observations in Figure 5. By assumption, the probability of dying per unit time is fixed for a specific temperature as illustrated in Figure 5b. For all populations, the probability of replication is initially zero and increases over time depending on the number of alive cells in the population. With alive cells secreting the extracellular factor, the probability of replicating quickly exceeds the probability of dying for high initial population-densities in the deterministic growth phase after a few units of time (top row). Eventually the probability of replicating plateaus at the maximally allowed value. For initial population-densities in the random growth phase (2nd and 3rd rows), the number of alive cells initially decreases over time as the probability of replicating approaches – but is smaller than – the probability of dying. Simultaneously, this continuously decreasing pool of alive cells secretes the extracellular factor, resulting in the probability of replicating continuously increasing - albeit with decreasing rate - over time. After some point in time (around 300 hours in the graphs shown here), the probability of replicating is very close to the probability of dying, and very few alive cells are left in the population. Here, the probability of replicating either exceeds the probability of dying - leading to net growth of the population - or the probability of replicating comes close to but remains smaller than the probability of dying - leading to a population extinction. For initial population-densities in the no-growth phase (bottom row), the probability of replicating remains well below the probability of dying. Here, the population goes extinct before the alive cells can accumulate sufficient concentration of the extracellular factor to increase the probability of replicating. (**b**) Besides all other features (shown in Figure 5), the model also reproduces the number of red cells and white cells, including the sustained population of few white cells in the random growth and no-growth phase, that we observed in Supplementary Fig. 5. Shown here are the number of alive (yellow points) and dead cells (red points) over time that our model produced for the same populations as in (a). For initial population-densities in the deterministic-growth phase (top row), the population grows exponentially as the probability of replicating quickly exceeds the probability of dying. For initial population-densities in the random-growth phase (2nd and 3rd rows), the number of alive cells over time reflects the fine balance between the probability of replicating and probability of dying. Here the population is ultra-sensitive to the state transitions of the very few alive cells (at ~300 hours). Based on whether these alive cells stay alive without replicating, replicate, or die in the next time steps, the population can eventually either expand and grow exponentially or go extinct in subsequent times. Finally, for initial population-densities in the no-growth regime (bottom row), the number of alive cells decreases exponentially as the probability of replicating remains close to zero and does not approach the probability of dying, leading to a net death until the population goes extinct. (**c-d**) Schematic summary that outlines the main features of the model. All cell populations eventually either grow exponentially (**c**) or go extinct (**d**). Whether a population grows exponentially or goes extinct is determined by the balance between a decreasing number of alive cells that continuously accumulate the extracellular factor, which in turn determines the probability of replicating for those alive cells. There is a net decrease per unit time in the number of alive cells as long as the probability of replicating is smaller than the probability of dying. If a sufficient amount of the extracellular factor accumulates, then the probability of replicating at some time exceeds the probability of dying, leading to net - exponential - growth if there are any alive cells remaining (c). As long as a sufficient amount of extracellular factor does not accumulate before all cells die, the probability of replicating remains below the probability of dying, leading to a decrease in the number of alive cells and eventually a population extinction (d). (**e**) A competition between two elements - a constant probability of dying and an initially lower probability of replicating that keeps approaching the probability of dying, evermore closing the gap between the two values - results in the population whose approach to extinction continuously slows down over time, leading to the number of survivors decreasing over time as a heavy-tailed (power-law-like) function (see Supplemental Notes). (**f**) Finally, a consequence of our model, which recapitulates all the main features of the experimental data, is that cells can replicate, in fact, at extremely high temperatures (e.g. 45 $^{O}$C) for which only the no-growth phase exists in the population-level phase diagram. But this occurs with a vanishingly low probability,

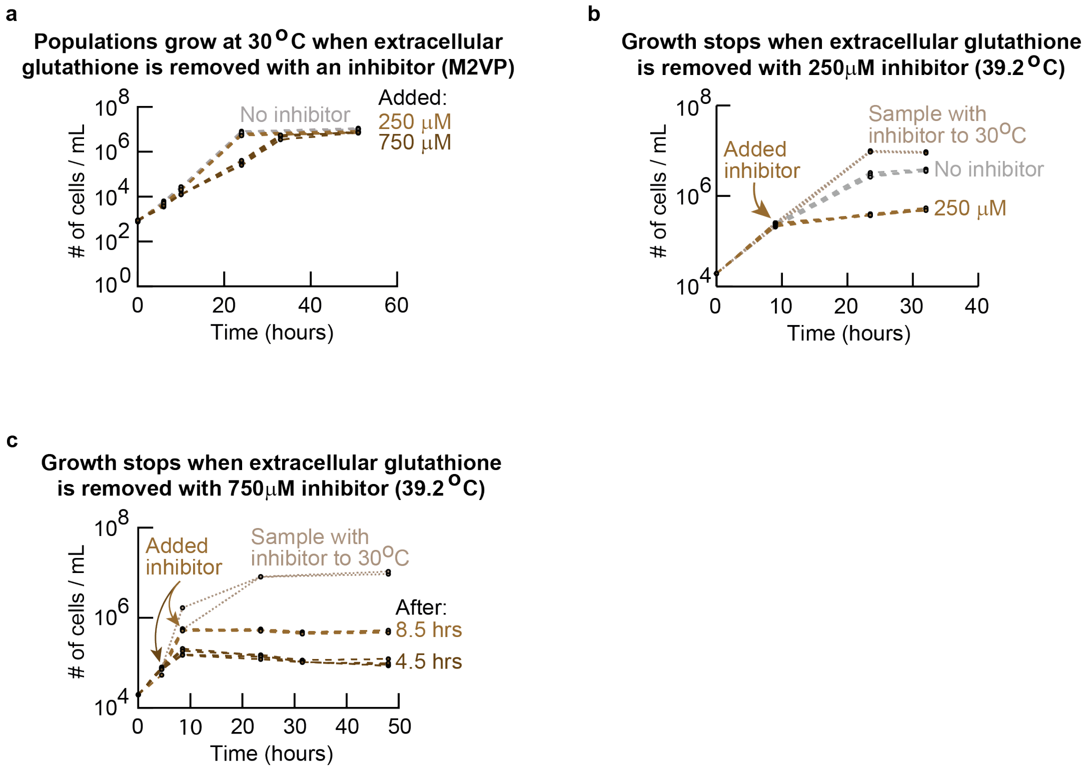


**Supplementary Fig. 15. Chemically masking reduced glutathione in the extracellular environment stops population growths at high temperatures (Related to Figure 6a).** (**a**) We used a thiol scavenging agent, 1-methyl-2-venylpyridinium (M2VP), to rapidly scavenge and mask all reduced glutathione (see Methods). Populations at 30 $^{O}$C were incubated with the masking reagent at 0 µM, 250 µM, and 750 µM. These populations exponentially grew at 30 $^{O}$C. These results show that the masking reagent (M2VP) does not interfere with intracellular processes and only scavenges extracellular glutathione (note that log-phase cells do not secrete glutathione at 30 $^{O}$C). (**b-c**) At 39.2 $^{O}$C. Deterministically growing populations at 39.2 $^{O}$C were subjected to 250 µM of the masking reagent after ~10 hours of incubation (**b**) or 750 µM of the masking reagent after 4.5 hours or 7.5 hours of incubation (**c**). All populations stopped growing after the masking reagent was added. These results show that removing extracellular, reduced glutathione stops all growths at a high temperature. Aliquots of these populations, from 39.2 $^{O}$C, were transferred to 30 $^{O}$C after the masking reagent was added (dotted light brown curves). These populations continued to exponentially grow to the carrying capacity in 30 $^{O}$C. For (a-c), all colors show at least four replicate populations.

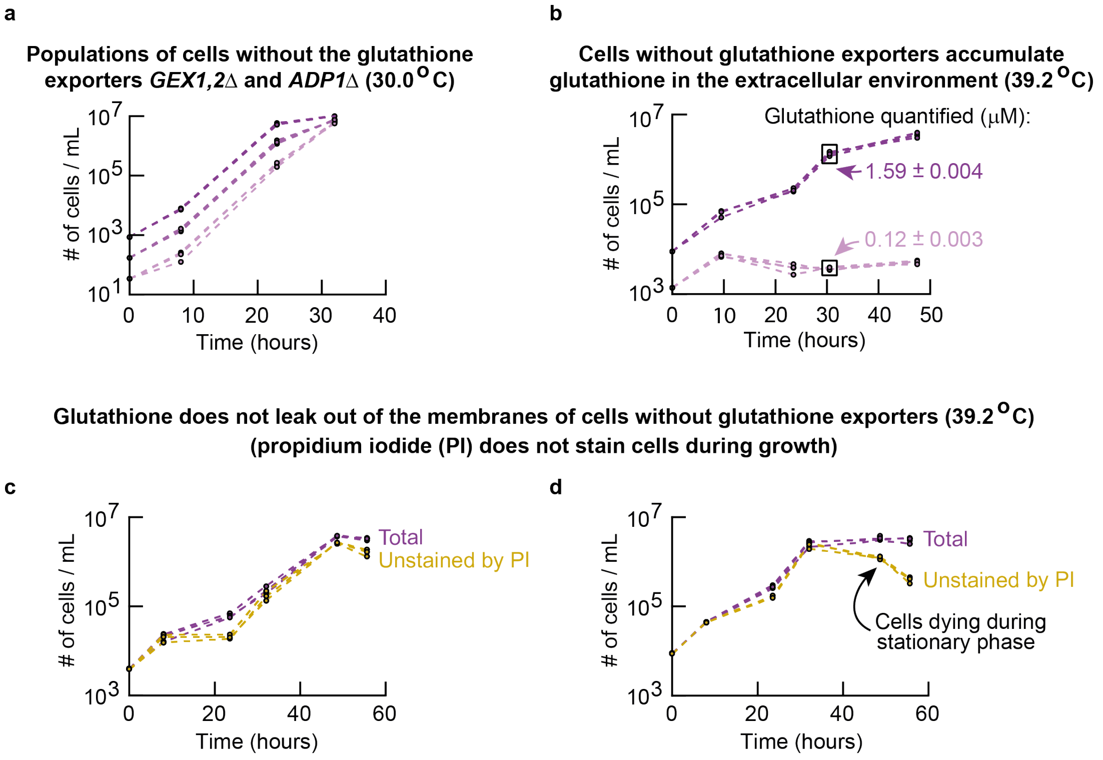


**Supplementary Fig. 16. Mutant yeasts lacking several glutathione exporters can still secrete glutathione while growing at high temperatures but not because their membranes are damaged, which would cause leakages of glutathione.** We constructed a mutant strain by knocking out, in the wildtype strain, all three genes that encode three of the known, major glutathione exporters: *GEX1*, *GEX2* and *ADP1* (see "Mutant yeasts" in the Methods section). (**a**) At 30 $^{O}$C. Population density (number of cells/mL) measured over time for populations of the mutant strain that lacks all three exporter genes, starting with different initial population-densities. (**b**) Populations of the mutant strain with different starting densities at 39.2 $^{O}$C ($n$ = 3). For each population, we measured the extracellular glutathione concentration after 30 hours of incubation at 39.2 $^{O}$C (indicated in the plot). These measurements show that growing populations of the mutant strain still secrete glutathione into the extracellular media. (**c-d**) Population density measured over time for the mutant strain with different starting densities at 39.2 $^{O}$C (purple curves, $n$ = 3). For each population and for every timepoint, we took an aliquot of the liquid culture and incubated it with 1 μg/mL of propidium iodide for 20 minutes at room temperature. We then flowed these aliquots through a flow cytometer to measure the number of cells that were not stained by the propidium iodide (yellow curves). Cells that were stained by the propidium iodide have lost their membrane integrity - this is what the commonly used propidium-iodide assay determines; it flows into the cells and stains their DNA if and only if their membranes are damaged. Conversely, cells that were not stained by the propidium iodide had intact membranes. The fact that almost all the cells in the populations were unstained by the propidium iodide (i.e., the purple and yellow curves nearly perfectly overlap in (c-d)) shows that glutathione does not simply leak out of the mutant cells - their membrane integrity was maintained during their growths at the high temperature. In other words, the mutant strain exports glutathione through other export mechanisms besides those mediated by the three genes that we knocked out.

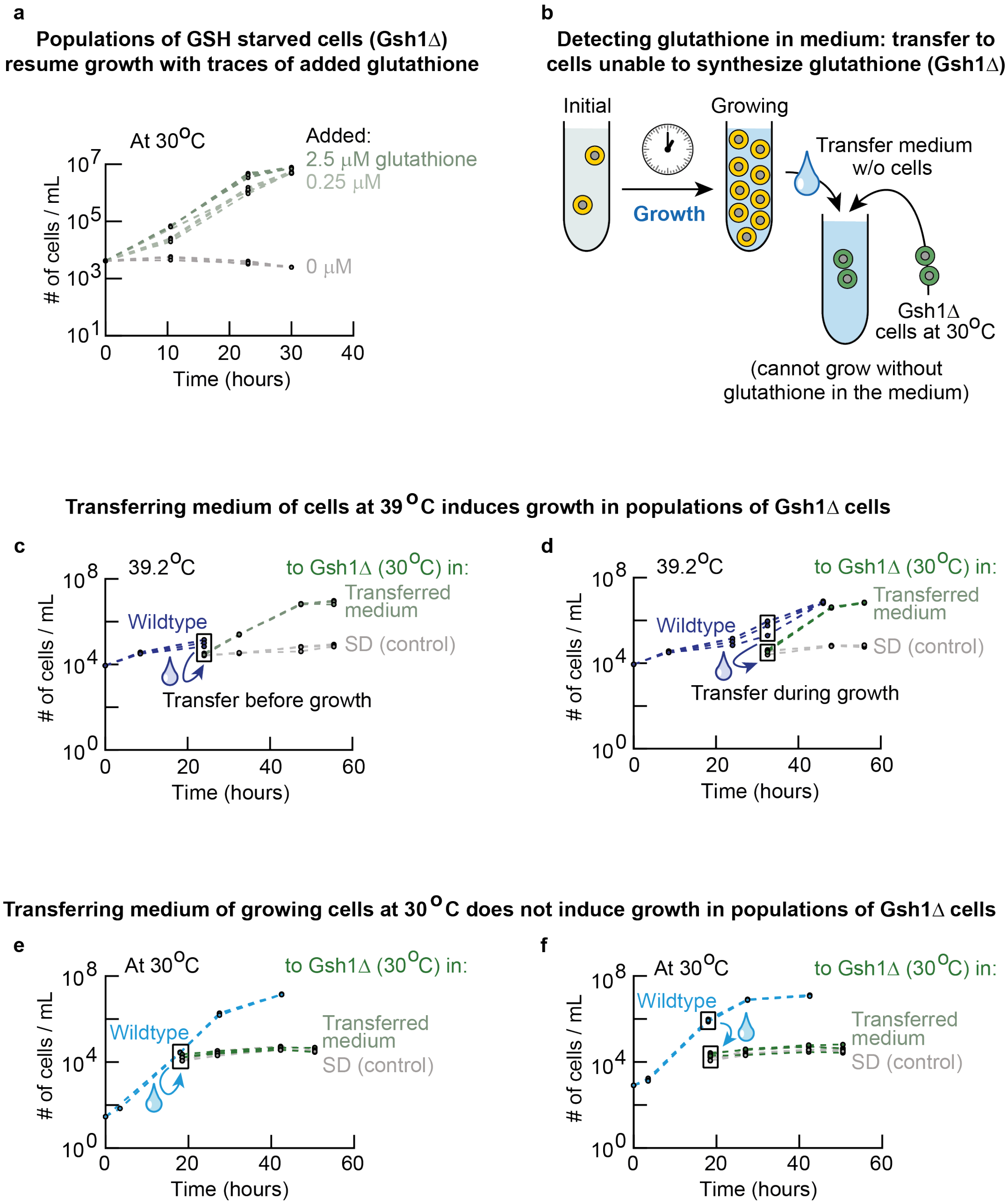


**Supplementary Fig. 17. Using a mutant strain that cannot synthesize glutathione (GSH1Δ-strain) to demonstrate, once again, that wildtype yeasts secrete glutathione at high temperatures (Related to Figure 4e and Supplementary Fig. 13).** (**a**) We constructed a mutant strain that could not synthesize glutathione by knocking out, in the wildtype strain, the *GSH1* gene which is essential for glutathione biosynthesis (see "Mutant yeasts" in the Methods section). Glutathione has an essential intracellular role in budding yeast that is separate from heat-shock responses. So, the GSH1Δ-mutant can only grow in media that we supplement with glutathione. To check this, we starved the GSH1Δ-mutant cells in the usual minimal media overnight without any glutathione. We then incubated these mutant cells with 0 μM, 0.25 μM or 2.5 μM of glutathione that we added to the media. These cells did not grow in medium without any glutathione (0μM) but they grew in media with (very small amounts of) glutathione (i.e., when there were more than 0.25μM of extracellular glutathione). (**b**) Complementary to and as a double check of the results in Supplementary Fig. 13, we show here that, at high temperatures, cells in both random-growth phase and deterministic-growth phase indeed secrete glutathione as we found earlier (Supplementary Fig. 13). To show this, we used the GSH1Δ-mutant strain. To do so, we separated the growth media from cells grown at a fixed temperature by flowing the liquid cultures through 0.45μm-pore filters as previously described. We confirmed that there were no cells left behind in the filtered media by flowing the filtered media through a flow cytometer (we did not detect any cells in the flow cytometer). We then transplanted GSH1Δ-mutant cells that were starved of glutathione into these filtered media at 30 $^{O}$C. Subsequently, we then measured the resulting population densities over time in 30.0 $^{O}$C. (**c-d**) Growth media taken from wildtype cells that were deterministically growing at 39.2 $^{O}$C by filtration. We filtered the medium from the wildtype cells either right before the wildtype population started to grow (**c**) or while it was growing (**d**). We then gave these filtered media to GSH1Δ-cells and incubated them at 30.0 $^{O}$C (green curves in (c-d)). These mutants grew to reach the carrying capacity in both (c) and (d). As a control, in fresh media without any glutathione, we incubated populations of GSH1Δ-cells at the same starting density as the green curves (the ones that received the wildtype cells' filtered media) (grey curves in (c) and (d)). These control populations did not grow. (**e-f**) Same experiments as in (c-d) but now having grown wildtype cells at 30.0 $^{O}$C before taking away their media into which we incubated GSH1Δ-cells at 30.0 $^{O}$C. For all colors in each panel, there are at least three replicate populations. None of the green curves - representing the GSH1Δ-cells transplanted into the filtered media - show any appreciable growths. This result complements Supplementary Fig. 13: these results support our earlier finding that wildtype cells at high temperatures (above 36 $^{O}$C) secrete glutathione at micromolar concentrations.

### I. Model Summary

This section summarizes the most important analytical results on the model for yeast growth. Please refer to the subsequent sections below for further mathematical argumentation and all derivations. The simplest stochastic model for yeast growth at high temperature is that, per unit time, cells replicate and cells die with fixed probabilities. In such a linear model, the average behavior of the population is independent of population size, and either all grow or go extinct. As the population density also dictates whether a population grows or not, any such linear model is unable to reproduce the behavior of yeasts we observed experimentally.
The presense or emergence of cells with a heritable trait, such as persister-like cells or heat-tolerant mutants that can replicate at high temperature and pass this property on to their offspring, also cannot explain our data. In our experiments, we use $c = 8$ replicate populations per condition, and the largest inital population size is $k = 25$-fold larger than the smallest one. An upper bound for the probability to observe the outcome of our experiments if the mechanism were a heritable trait is then given by (see section III),

$$P_{c,k} \leq k^c \cdot \left(\frac{1}{k+1}\right)^{c+\frac{c}{k}} = 0.26. \tag{1}$$

We were able to produce these results many times (see Figures 2a-c and Figures S4-5), such that a model with a mechanism based on heritable traits cannot explain our experimental observations. Hence, we need an extended non-linear stochastic model to reproduce the data. To this end we use experimental observations (Figures 4a-g). Our data suggests that cells secrete gluathione that allows for cell growth when glutathione accumulates sufficiently (Figures 4a-g and Figures. S9-13). We therefore extend the simplest model, by assuming that the probability of replicating of cells depends on the concentration of extracellular glutathione that cells secrete at a constant rate.

The full model is as follows. Let $A_t$ be the population-size of alive cells at time $t$ with initial population-size $A_0$. Per unit time, any cell dies with probability $p_d(T)$ linearly increasing with the temperature $T$. Moreover, cells replicate with probability $p_a(t)$, where $p_a(t)$ is given by the maximum probability of replicating $\mu$ that is scaled by a Hill function depending on the concentration of the extracellular glutathione $m_t$ and constant $k(T)$. Finally, the extracellular glutathione accumulates by secretion of alive cells with rate $r_m(T)$. We describe the total population size at time $t$ with $N_t$, and let $N_{birth}(t)$ and $N_{death}(t)$ be the number of births and deaths of cells at time $t$. Then the stochastic model describing yeast growth at high temperature is given by,

$$\begin{aligned} N_{birth}(t) &\sim \text{Binom}\big(A_t, p_a(t)\big), \\ N_{death}(t) &\sim \text{Binom}\big(A_t, p_d(T)\big), \\ A_{t+1} &= A_t + N_{birth}(t) - N_{death}(t), \\ p_a(t) &= \mu \cdot \frac{m_t}{k(T) + m_t}, \\ m_{t+1} &= m_t + r_m(T) A_t, \end{aligned} \tag{2}$$

where $A_0$ is the initial population-size, and the initial probability of replicating is given by $p_a(0) = 0$. The total population size changes according to,

$$N_{t+1} = N_t + N_{birth}(t). \tag{3}$$

This model reproduces all the main features we observed experimentally (Figure 5 and Figure S14). All simulations were run using one single set of parameters, choosing the temperature $T$ and initial population-size $A_0$ appropriately. The parameters used to fit the model to our experimental data are the maximum probability of replicating $\mu = 0.25$ (approximating the maximum growth rate of our wild-type yeast), $K = k(T)/r_m(T) = 30,000$ (chosen such that order of magnitude of the phase boundary matches the boundary we found experimentally, Figure 2d) and the probability of dying depending on temperature, $p_d(T) = \mu \cdot \frac{T - T_{min}}{T_{max} - T_{min}}$ with $T_{min} = 37.9$°C and $T_{max} = 40.2$°C (chosen such that the endpoints of the phase boundary match the boundary we found experimentally, Figure 2d).

An deterministic approximation of the stochastic model allows us to derive an analytical expression for the phase boundary between the deterministic growth phase and the no-growth phase (Section VI). The analytical expression for the phase boundary is given by (simplified form of 72),

$$A_0 \propto \frac{K \cdot p_d^2(T)}{\mu - p_d(T)}. \tag{4}$$

Hence, the initial population-size required for growth diverges as the probability of dying approaches the probability of replicating in the model. Finally, the deterministic approximation is used to show that - in the no-growth regime where the population does not grow - the decrease of the population of alive cells is not appropriately described by exponential decay (Section VII). Instead, the instantaneous rate at which the number of alive cells in the population changes continuously decreases. This is the result of the probability of replicating approaching the probability of dying. Therefore the decay of the number of alive cells in the population follows a heavy-tailed function, as we also find in our experiments (Figures 3a-b and Figures S7-8).

#### II. Simple model description

First, we consider the simplest stochastic model for yeast growth at high temperature. To this end, we assume that all cells are identical and independent of each other (i.i.d.). Let $A_t$ be a random variable representing be the number of alive cells at time $t$. Per unit time, cells replicate with probability $p_a$ and cells die with a probability $p_d(T)$ that is monotonically increasing with temperature $T$. Then $\{A_t\}_{t \geq 0}$ is a discrete-time Markov process describing yeast growth at high temperatures. Let $N_t$ be the total population size at time $t$ and describe the number of births and deaths with $N_{birth}(t)$ and $N_{death}(t)$ respectively. Then the simple model is described by,

$$N_{birth}(t) \sim \text{Binom}\big(A_t, p_a\big), \tag{5}$$

$$N_{death}(t) \sim \text{Binom}\big(A_t, p_d(T)\big), \tag{6}$$

$$A_{t+1} = A_t + N_{birth}(t) - N_{death}(t). \tag{7}$$

with the total population size changing according to,

$$N_{t+1} = N_t + N_{birth}(t). \tag{8}$$

We can approximate the stochastic model as follows. As both cell replication and death follow a Binomial distribution with parameters $p_a$ and $p_d(T)$ respectively, we have,

$$\mathbb{E}[A_{t+1}] = A_t + p_a A_t - p_d(T) A_t. \tag{9}$$

By approximation for large $A_t$, we then obtain the following linear differential equation describing the system,

$$A_{t+1} - A_t \approx \Big(p_a - p_d(T)\Big) \cdot A_t \tag{10}$$

$$\frac{dA}{dt} \approx \Big(p_a - p_d(T)\Big) \cdot A. \tag{11}$$

It follows that, for a sufficiently large initial population of replicating cells $A_0$, the number of alive cells in the population can be modeled by,

$$A(t) = A_0 \cdot \exp\Big((p_a - p_d(T))t\Big). \tag{12}$$

#### III. Necessity of a non-linear model

In this section we prove that we need a non-linear model to describe yeast growth at high temperature. To see this, suppose we have a linear model. The parameters of this model are fixed (i.e. there is no emergence of heritable traits such as persister-like cells or mutants), and all cells are autonomous (independent). In our experiments, we find that, at 39°C, populations with initially $400$ cells/mL never grow, while populations with initially $10,000$ cells/mL always grow (Figure 2b). Now suppose we simulate populations with initially $400$ cells/mL with this linear model. We can run a simulation of this population many times (say 25x), all with the same result (no growth). As cells are autonomous, we can combine these twenty-five simulations into one single simulation without changing the outcome. Then we simulate a population with initially $25\text{x}400 = 10,000$ cells/mL, and we would obtain the same result (no growth). This is because the cells are autonomous: the cells do not care what size their population is. This contradict our experimental results, as we find that the population of initially $10,000$ cells/mL always grows (at 39°C). This arguments shows that no linear model with fixed parameters can reproduce our experimental data.

This can also be seen from the linear model from Section II. The behavior of the model, as described by 12, is completely independent of the initial population-size of replicating cells $A_0$. When $p_a > p_d(T)$, the population will grow exponentially. In contrast, growth is impossible if $p_a < p_d(T)$ and then the population goes extinct. Hence, yeast growth ceases at the temperature where the probability of dying $p_d$ exceeds the probability of replicating $p_a$ for populations of any size. Thus, the linear model from Section II cannot explain our experimental data.

What the arguments above do not consider, is the emergence of cells in a population with a heritable trait. Next, we suppose that the ability to grow at a high temperature is a heritable trait (e.g., persisters or heat-tolerant mutants). These are mechanisms in which one special cell with this heritable trait (e.g. having its probability of replicating larger than its probability of dying, $p_d(T) < p_a$) emerges in the population and is responsible for giving rise to a whole lineage of growing cells. For convenience, we will refer to cells that can give rise to a whole lineage of growing cells as "persisters" here. To see how such a mechanism cannot explain our data, note that the persisters must exist in sufficiently low abundances in a population so that populations that start with low densities never grow. At the same time, the persisters must occur frequently enough such that populations that start with high enough densities always grow. As soon as a population has a persister, it will grow until it reaches a carrying capacity due to the persister yielding a growing population. The following proof shows that, regardless of the frequency at which the persisters initially are present or later emerge in a population, the probability that such a mechanism reproduces our data is too small to be consistent with our data.

Experimentally we observe populations of cells of some initial size $N_0$ that never grow (Figures 2a-c and Figures S4-5). Moreover, in our experiments we use a $k = 25$-fold difference in initial population size between the largest and smallest populations. Thus, the largest populations initially have $k \cdot N_0$ and always grow. Consider populations of cells with intially $N_0$ and $k \cdot N_0$ cells. We assume that cells are identical and independent of all other cells (i.i.d.), and that these cells are unable to grow at high temperature ($p_d(T) > p_a$). Let $p_g > 0$ be the probability that a cell is or will become a persister that will replicate at high temperature, and passes on this ability to replicate to its offspring ($p_d(T) < p_a$). Then the probability that the culture with an initial population of size $N_0$ will never exponentially grow is the probability that none of the cells becomes a persister, given by,

$$P_{\text{no growth}}(N_0) = (1 - p_g)^{N_0}. \tag{13}$$

Moreover, the probability that the culture with initial population-size $k \cdot N_0$ will eventually exponentially grow is the probability that some cell becomes a persister, given by,

$$P_{\text{growth}}(k \cdot N_0) = 1 - (1 - p_g)^{k \cdot N_0}. \tag{14}$$

Hence, the probability to observe $c$ cultures with initial population-size $N_0$ never grow exponentially, and simultaneously $c$ cultures with initial population-size $k \cdot N_0$ to all grow exponentially is given by,

$$\begin{aligned} P_c(N_0) &= \Big(P_{\text{no growth}}(N_0)\Big)^c \cdot \Big(P_{\text{growth}}(k \cdot N_0)\Big)^c & (15)\\ &= \Big((1-p_g)^{N_0}\Big)^c \cdot \Big(1 - (1-p_g)^{k \cdot N_0}\Big)^c. & (16) \end{aligned}$$

Here $P_c(N_0)$ gives the probability to observe the outcome we observe in our experiments: all cultures with low initial population-size do not grow, while all cultures with high initial population-size do grow exponentially. To maximize the probability of observing our experimental outcome, we therefore want to maximze $P_c(N_0)$ for the only free variable $p_g$. To simplify notation, let $x := (1 - p_g)^{N_0}$. Then,

$$\begin{aligned} P_c(N_0) &= x^c \cdot (1 - x^k)^c & (17)\\ &= \big(x - x^{k+1}\big)^c. & (18) \end{aligned}$$

Taking the derivative to maximize $P_c(N_0)$,

$$\begin{aligned} \frac{dP_c}{dp_g} &= \frac{dP_c}{dx} \cdot \frac{dx}{dp_g} & (19)\\ &= c\big(x - x^{k+1}\big)^{c-1} \cdot \big(1 - (k+1)x^k\big) & (20)\\ &\quad \cdot N_0(1-p_g)^{N_0-1} \cdot -1 & \end{aligned}$$

Notice that $\frac{dP_c}{dp_g}$ is zero for the trivial solutions $p_g = 0$ and $p_g = 1$. The nontrivial solution of $\frac{dP_c}{dp_g} = 0$ is described by,

$$1 - (k+1)x^k = 0, \tag{21}$$

which yields the following solution that maximizes $P_c(N_0)$,

$$x = \Big(\frac{1}{k+1}\Big)^{\frac{1}{k}}. \tag{22}$$

Therefore, the probability $P_c(N_0)$ that describes the outcome we observe in our experiments is maximized for,

$$(1-p_g)^{N_0} = \left(\frac{1}{k+1}\right)^{\frac{1}{k}}, \tag{23}$$

hence the probability $p_g$ - the probability of being a persister cell - that maximizes the probability of observing our experimental outcome is given by,

$$p_g = 1 - \left(\frac{1}{k+1}\right)^{\frac{1}{kN_0}}. \tag{24}$$

Finally, by substititing 24 into 15 the actual probability $P_c(N_0)$ that describes the outcome we observe in our experiments is bounded by,

$$P_c(N_0) \leq \left(\frac{1}{k+1}\right)^{\frac{c}{k}} \cdot \left(1 - \frac{1}{k+1}\right)^{c} \tag{25}$$

$$= k^c \cdot \left(\frac{1}{k+1}\right)^{c+\frac{c}{k}}. \tag{26}$$

This upper bound for the outcome we observe in our experiments only depends on the number of replicate populations $c$ per condition and the dilution factor $k$ between these conditions. As described above, we use $c = 8$ replicate populations per condition and a $k = 25$-fold difference between the largest and smallest initial population-sizes in our experiments. Then an upper bound for the probability to observe the outcomes of our experiments is given by substituting $c = 8$ and $k = 25$ into 25, leading to $P_c(N_0) \leq 0.26$. As we consistently make these experimental observations (Figures 2a-c and Figures S4-5), the presense or emergence of persister cells cannot explain our observations.

#### IV. Non-linear model definition

We conclude that a simple, linear model 12 is insufficient to describe the behavior of our yeast cells at high temperature. Moreover, our data suggests that cells secrete glutathione that accumulates extracellularly and allows for cell growth when a sufficient concentration has been reached (Figures 4a-g and Figures S9-13). As our cells are genetically identical, we can safely assume that cells are not independent (not autonomous). We therefore extend the simplest model with secretion of glutathione and an effective probability of replicating that depends on the extracellular concentration of glutathione.

Similar to the simple model 12, let $A_t$ be the population size of alive cells at time $t$. Per unit time, any cell dies with probability $p_d(T)$ depending on the temperature $T$. In contrast with the simplest model, we now assume that the probability of replicating is not constant, based on our observation that population-sizes can remain constant while still containing alive cells (random phase, Figures 2a-b). Hence, assume that cells replicate with probability $p_a(t)$, where $p_a(t)$ is some maximum probability of replicating $\mu$ scaled by a Hill function (Michaelis-Menten) depending on the concentration of extracellular glutathione $m_t$ and constant $k(T)$. Finally, the extracellular glutathione accumulates by constant secretion of alive cells with secretion rate $r_m(T)$. Again describe the total population size at time $t$ with $N_t$ and let $N_{birth}(t)$ and $N_{death}(t)$ be the number of births and deaths of cells at time $t$. Then the full stochastic model is described by,

$$
\begin{aligned}
N_{birth}(t) &\sim \text{Binom}\big(A_t, p_a(t)\big),\\
N_{death}(t) &\sim \text{Binom}\big(A_t, p_d(T)\big),\\
A_{t+1} &= A_t + N_{birth}(t) - N_{death}(t),\\
p_a(t) &= \mu \cdot \frac{m_t}{k(T) + m_t},\\
m_{t+1} &= m_t + r_m(T)A_t,
\end{aligned}
\tag{27}
$$

with the total population size changing according to,

$$N_{t+1} = N_t + N_{birth}(t). \tag{28}$$

The behavior of this model is completely different than the simplest model (Section II). Here, the probability of replicating $p_a(t)$ increases (monotonically) over time as function of the number of alive cells. Hence, as $p_a(0) = 0$, there is no guarantee that any population of cells will grow exponentially, unless the cells accumulate sufficient extracellular glutathione $m_t$ such that $p_a(\tau) > p_d(T)$ for some time $\tau > 0$. This model 27 is studied in more detail with simulations (Figure 5 and Figure S14) and analytically with an approximation in the following sections.

#### V. Deterministic approximation

We analytically study the model 27 next, for which we use a deterministic approximation to gain insight into some key features of the model. In our model both cell replication and death follow a Binomial distribution, such that the number of alive cells at the next time step can be estimated by,

$$\mathbb{E}[A_{t+1}] = A_t + p_a(t)A_t - p_d(T)A_t. \tag{29}$$

By approximation, we obtain the following nonlinear system of equations:

$$A_{t+1} = A_t + p_a(t)A_t - p_d(T)A_t, \tag{30}$$

$$p_a(t) = \mu \cdot \frac{m_t}{k(T) + m_t}, \tag{31}$$

$$m_{t+1} = m_t + r_m(T)A_t. \tag{32}$$

First, we rewrite this system into a more convenient form. We rescale the extracellular glutathione as $M_t = m_t/r_m(T)$ and $K(T) = k(T)/r_m(T)$. Here, $K(T)$ now represents the constant relative to the production rate. Thus we obtain the following simplified determinist approximation of the stochastic model 27 describing growth at high temperature,

$$A_{t+1} = A_t \cdot \Big(1 + p_a(t) - p_d(T)\Big) \tag{33}$$

$$p_a(t) = \mu \cdot \frac{M_t}{K(T) + M_t} \tag{34}$$

$$M_{t+1} = M_t + A_t. \tag{35}$$

##### A. Interpretation

The relative change of the number of alive cells in the population is determined by the factor $1+p_a(t)-p_d(T)$ which depends on time (probability of replicating) and temperature (probability of dying). Here the number of alive cells on average increases when $p_a(t) > p_d(T)$ and on average decreases when $p_a(t) < p_d(T)$. Notice that $p_a(t) \leq \mu$ for all $t > 0$ by choice of the Hill function. Moreover, as $p_a(0) = 0$, the population of alive cells in the population initially decreases exponentially (approximately with a factor $1 - p_d(T)$ per unit time). For the maximum probability of replicating $\mu$ and probability of dying $p_d$ we can destinguish three cases:

- $p_d < \mu$: In the limit of the concentration of the extracellular glutathione that accumulates ($M_t \to \infty$) we have $p_a(t) \to \mu$, such that $p_a(t) > p_d$ for some $t > 0$ and therefore the population can grow exponentially.
- $p_d \approx \mu$: Here the probability of dying is very close to the maxium probability of replicating for a cell. Hence, only large populations of alive cells can sustain the population as $p_a(t) \to p_d$ only when the concentration of extracellular glutathione increases ($M_t \to \infty$). The population cannot grow exponentially.
- $p_d > \mu$: The probability of dying is always higher than the maxium probability of replicating, and the population of alive cells on average decreases, and is guaranteed to go extinct.

##### B. Probability of dying depends linearly on temperature

The qualitative behavior of the model is fixed for a given probability of dying. Without loss of generality, a sensible assumption is to let the probability of dying for a cell increase monotonically with temperature - for two different temperatures, the probability of dying for the higher temperature is at least the probability of dying for the lower temperature. Since we know that all populations grow at $T = 37.9$°C and all populations do not grow at $T = 40.2$°C (Figure 2d), the simplest assumption is to linearly increase the probability of dying between these values such that $p_d(40.2) = \mu$. Any non-linear, but still montonically increasing probability of dying as function of temperature yields the same qualitative behavior of the model but displays these behaviors at different temperatures.

##### C. Glutathione secretion rate is constant

Instead of being a constant, one can set the glutathione secretion rate $r_m(T)$ to be dependent on, for example, the population size $A_t$ or glutathione concentration $m_t$. This changes the treshold $K(T)$, which in turn merely shifts Hill function describing the probability of replicating $p_a(t)$ as function of glutathione concentration 33. Thus, choosing a non-constant secretion rate or treshold concentration of glutathione modifies the probability of replicating in the model. However, the qualitative behavior of the model does not change upon a different, sensible, choice for the glutathione secretion rate $r_m(T)$ (or treshold $K(T)$). For example, we could choose $r_m(T)$ to be linearly dependent on the population size $A_t$ - larger populations secrete more glutathione. Reversely, we could set the secretion rate $r_m(T)$, for example, inversely proportional to glutathione concentration $m_t$ - low glutathione concentrations trigger a higher secretion rate than high glutathione concentrations. These choices for non-constant parameters of the model increase or decrease the sensitivity of populations to the initial population size, but do not change the qualitative growth behavior of the model (i.e. the existence of no-growth, random growth and deterministic growth). We therefore choose the simplest assumptions for our model by having these parameters ($r_m(T)$ and $K(T)$) constant.

#### VI. DESCRIPTION OF THE PHASE BOUNDARY

The goal of this section is to derive an description of the phase boundary of our model that we observe in simulations (Figure 5). In contrast to the simple model 12, the non-linear model 27 allows for a population of cells with $p_a(0) < p_d(T)$ that can still exponentially grow for some $t > 0$ due to the accumulation of extracellular glutathione. Although the model 27 is stochastic, we can use the deterministic approximation 33 to gain some insight into the shape of the phase boundary and how the behavior of the model depends on the variables of the model. To this end, notice that any cell population eventually either exponentially grows or goes extinct. Therefore, without loss of generality, assume that there exists some $\epsilon > 0$ such that a cell population will (on average) grow exponentially when $p_a(t) > \epsilon p_d$ for some $t > 0$. Equivalently, a cell population will go extinct if $p_a(t) \leq \epsilon p_d$ for all $t > 0$. Note that $\epsilon = 1$ would suffice. For now we ignore the dependence of cell populations on temperature, and write $K = K(T)$ and $p_d = p_d(T)$. The approach here is as follows: First we derive upper and lower bound on the number of alive cells in the population, followed by bounds for the concentration of the extracellular glutathione. Finally, all bounds are used to derive an approximate description of the phase boundary in the simulated phase diagram (Figure 5).

##### A. Bounds on number of alive cells

First, we determine a lower and upper bound on the population size of alive cells when the population is not growing exponentially ($1 + p_a(t) - p_d(T) < 1$ for 33). Notice that, by recursive substitution of 33,

$$A_{t+1} = A_0 \cdot \prod_{s=0}^{t} \Big(1 + p_a(s) - p_d\Big), \tag{36}$$

where $A_0$ is the initial population-size of alive cells at time $t = 0$. Moreover, the probability of replicating is bounded by $p_a(s) \geq 0$, such that,

$$\begin{aligned} A_t &= A_0 \cdot \prod_{s=0}^{t-1} \Big(1 + p_a(s) - p_d\Big) && (37)\\ &\geq A_0 \cdot \prod_{s=0}^{t-1} \Big(1 - p_d\Big) && (38)\\ &= A_0\Big(1 - p_d\Big)^t. && (39)\end{aligned}$$

Next, suppose that the cell population will go extinct. Then $p_a(s) < \epsilon p_d$ for all $s > 0$ by assumption, and,

$$\begin{aligned} A_t &= A_0 \cdot \prod_{s=0}^{t-1} \Big(1 + p_a(s) - p_d\Big) && (40)\\ &< A_0 \cdot \prod_{s=0}^{t-1} \Big(1 - p_d(1 - \epsilon)\Big) && (41)\\ &= A_0\Big(1 - p_d(1 - \epsilon)\Big)^t && (42)\end{aligned}$$

Hence, when a cell population will go extinct, then the number of alive cells in the population at time $t$ is bounded by,

$$A_0\Big(1 - p_d\Big)^t \leq A_t < A_0\Big(1 - p_d(1 - \epsilon)\Big)^t. \tag{43}$$

##### B. Bounds on concentration of extracellular glutathione

Next, we determine a lower and upper bound on the concentration of the extracellular glutathione when the population of cells is not exponentially growing, similarly to the bound of the number of alive cells. Recursive substitution of $M_t$ and using 36 yields,

$$M_{t+1} = \sum_{s=0}^{t} A_s \tag{44}$$

$$= A_0 + \sum_{s=1}^{t} A_s \tag{45}$$

$$= A_0 + A_0 \sum_{s=1}^{t} \prod_{k=0}^{s-1} \Big(1 + p_a(k) - p_d\Big) \tag{46}$$

Moreover, we can bound the probability of replicating by $p_a(k) \geq 0$, such that,

$$M_{t+1} \geq A_0 + A_0 \sum_{s=1}^{t} \prod_{k=0}^{s-1} \Big(1 - p_d\Big) \tag{47}$$

$$= A_0 + A_0 \sum_{s=1}^{t} \Big(1 - p_d\Big)^s \tag{48}$$

$$= A_0 \sum_{s=0}^{t} \Big(1 - p_d\Big)^s. \tag{49}$$

Equation 49 represents the first $t+1$ terms of a geometric series that converges as its ratio satisfies $|1-p_d(T)| < 1$. Hence,

$$M_{t+1} \geq A_0 \cdot \frac{1 - \big(1 - p_d\big)^{t+1}}{p_d}. \tag{50}$$

Next, we seek an upper bound on the concentration of the extracellular glutathione. To this end, suppose that $p_a(k) < \epsilon p_d$ for all $k > 0$ such that the population is will go extinct by assumption. Then, starting from 46, and substituting $p_a(k) < \epsilon p_d$ and simplifying as in 49,

$$M_{t+1} = A_0 + A_0 \sum_{s=1}^{t} \prod_{k=0}^{s-1} \Big(1 + p_a(k) - p_d\Big) \tag{51}$$

$$< A_0 + A_0 \sum_{s=1}^{t} \prod_{k=0}^{s-1} \Big(1 - p_d(1-\epsilon)\Big) \tag{52}$$

$$= A_0 \sum_{s=0}^{t} \Big(1 - p_d(1-\epsilon)\Big)^s. \tag{53}$$

Again substituting the known sum of a geometric series we obtain,

$$M_{t+1} < A_0 \cdot \frac{1 - \big(1 - p_d(1-\epsilon)\big)^{t+1}}{p_d(1-\epsilon)}. \tag{54}$$

Hence, when the cell population goes extinct, 50 and 54 yield the following bounds for the concentration of the extracellular glutathione $M_t$ at time $t$,

$$A_0 \cdot \frac{1 - \big(1 - p_d\big)^{t}}{p_d} \leq M_t < A_0 \cdot \frac{1 - \big(1 - p_d(1-\epsilon)\big)^{t}}{p_d(1-\epsilon)}. \tag{55}$$

##### C. Growth versus extinction regime

The bounds 43 and 55 provide us with estimates of the number of alive cells in the population and the concentration of the extracellular glutathione when knowing that the population will go extinct. These bounds are usefull, as they provide the worst-case estimate for the accumulation of extracellular glutathione and population-size of alive cells. Using these bounds we seek a contradiction next. Assuming the worst-case scenario (extinction), we seek the initial population-size of alive cells $A_0$ for which the probability of replicating still exceeds the probability of dying before extinction. Hence the population cannot (on average) go extinct as - even in the worst-case - the population will accumulate sufficient extracellular glutathione to grow.

More specifically, we first determine a lower bound for the probability of replicating $p_a(t)$ at the time the cell population is not yet extinct as function of $A_0$. Here, the cell population is not yet extinct when 43,

$$A_t \geq A_0(1-p_d)^t = 1. \tag{56}$$

Let $\tau$ be the time of extinction. Then, by solving 56, the time of extinction is lower bounded by,

$$\tau > \frac{\log(1/A_0)}{\log(1-p_d)}. \tag{57}$$

Substitution of 57 in the lower bound for the concentration of the extracellular glutathione yields 55,

$$M_\tau \geq A_0 \cdot \frac{1-\left(1-p_d\right)^\tau}{p_d} \tag{58}$$

$$> \frac{A_0 - 1}{p_d}. \tag{59}$$

Hence the worst-case concentration of the extracellular glutathione right before extinction is lower bounded by 59. Finally, substitution of 59 into the probability of replicating in our model 33 gives, as $p_a(t)$ is monotonically increasing in $M_t$,

$$p_a(\tau) > \mu \cdot \frac{A_0 - 1}{K p_d + A_0 - 1}. \tag{60}$$

Recall that we assume that the cell population will exponentially grow when $p_a(t) > \epsilon p_d$ for some $t > 0$. Therefore the cell population will at some point grow exponentially when 60,

$$p_a(\tau) > \mu \cdot \frac{A_0 - 1}{K p_d + A_0 - 1} > \epsilon p_d, \tag{61}$$

Solving 61 for $A_0$ yields the following lower bound on the initial population-size of alive cells required to be able to grow exponentially,

$$A_0 - 1 > K \cdot \frac{\epsilon p_d^2}{\mu - \epsilon p_d}. \tag{62}$$

The above equation was derived from a contradiction: we assumed that a population goes extinct, and determine what minimum concentration of extracellularly accumulated glutathione it can produce. This concentration yields a lower bound of the probability of replicating at the time of extinction. Finally we derived a lower bound on the population size 62 for which this probability of replicating exceeds the probability of death, and the population will grow. Thus the population cannot go extinct.

Next, again using 43 and 55, we determine an upper bound for $A_0$ for which the cell population will go extinct. Recall that the cell population will go extinct when $p_a(t) \leq \epsilon p_d$ for all $t > 0$. Specifically, when $\tau$ is the time of extinction, we require $p_a(\tau) \leq \epsilon p_d$ as $p_a(t)$ is monotonically increasing. The number of alive cells when the population will go extinct is bounded from above by 43,

$$A_t < A_0\Big(1 - p_d(1-\epsilon)\Big)^t. \tag{63}$$

Then the cell population is extinct when $t$ solves,

$$A_0\Big(1 - p_d(1-\epsilon)\Big)^t = 1. \tag{64}$$

Let $\tau$ be the time of extinction. Then, by solving 64 we obtain an upper bound for the time of extinction,

$$\tau < \frac{\log(1/A_0)}{\log\big(1 - p_d(1-\epsilon)\big)}. \tag{65}$$

Substitution into the upper bound for the concentration of the extracellular glutathione 55 when the population goes extinct yields,

$$M_\tau < A_0 \cdot \frac{1 - \big(1 - p_d(1-\epsilon)\big)^\tau}{p_d(1-\epsilon)} \tag{66}$$

$$< \frac{A_0 - 1}{p_d(1-\epsilon)}. \tag{67}$$

The bound 67 gives an upper bound on the amount of extracellular glutathione a given population can accumulate at the time of extinction. Finally, substituting 67 into the probability of replicating yields,

$$p_a(\tau) < \mu \cdot \frac{A_0 - 1}{Kp_d(1-\epsilon) + A_0 - 1}. \tag{68}$$

Recall that we assume that the cell population will go extinct when $p_a(\tau) \leq \epsilon p_d$. Therefore the cell population indeed goes extinct if, by substitution into 68,

$$p_a(\tau) < \mu \cdot \frac{A_0 - 1}{Kp_d(1-\epsilon) + A_0 - 1} < \epsilon p_d. \tag{69}$$

Solving 69 for $A_0$ yields the following upper bound on the initial population-size of alive cells that guarantees that the population goes extinct,

$$A_0 - 1 < K \cdot \frac{\epsilon(1-\epsilon)p_d^2}{\mu - \epsilon p_d}. \tag{70}$$

In summary, we now found a boundaries for the initial population-size of alive cells that guarantees extinction 70 and for which the population is able to grow exponentially 62. For any initial population-size of alive cells in between, growth and extinction are unpredictable. Hence the random phase in our phase diagram (Figure 5) is described by 62, 70,

$$K \cdot \frac{\epsilon(1-\epsilon)p_d^2}{\mu - \epsilon p_d} < A_0 - 1 < K \cdot \frac{\epsilon p_d^2}{\mu - \epsilon p_d}. \tag{71}$$

The interpretation of these bounds is as follows. Suppose that we want cell populations to eventually grow when $p_a(t) > \epsilon p_d$ for some $t > 0$ and to go extinct when $p_a(t) \leq \epsilon p_d$ for all $t > 0$. Then, the initial population-size of alive cells that describes the phase boundary scales according to 71,

$$A_0 \propto 1 + K(T) \cdot \frac{p_d^2(T)}{\mu - p_d(T)} \tag{72}$$

Moreover, for the cell populations that go extinct, a lower bound for the extinction time is given by 57.

#### VII. Heavy-tailed decay to extinction

In this final section, we consider the extinction of cell populations. More specifically, we study the instantaneous rate of death of the number of alive cells in the population. A common assumption is that the number of alive cells follows some exponential decay over time. This is indeed the case when the probability of replicating $p_a(t)$ is constant, as then $1 + p_a(t) - p_d(T)$ is a constant over time 33. In contrast, in our model the probability of replicating $p_a(t)$ is monotonically increasing as a result of extracellular glutathione accumulating over time.

##### A. The instantaneous decay rate

First, we derive the instantaneous rate at which the number of alive cells in the population decreases. To this end, notice that the number of alive cells in the population is approximated by 33,

$$A_{t+1} = \Big(1 + p_a(t) - p_d(T)\Big) \cdot A_t. \tag{73}$$

We will only be interested in the rate of decay of the number of alive cells at time very close to some time $t^* > 0$. We therefore assume that $p_a(t)$ at time $t^*$ is temperarily a constant, $p_a(t) = p_a(t^*) := p_{a,t^*}$ for $t$ close to $t^*$, and thus independent of time. For insight, we further approximate 73 with the following linear differential equation,

$$\frac{dA}{dt} \approx \frac{A_{t+1} - A_t}{t + 1 - t} \tag{74}$$

$$= A_{t+1} - A_t \tag{75}$$

$$= -\Big(p_d(T) - p_a(t^*)\Big) \cdot A_t. \tag{76}$$

Solving this differential equation yields,

$$A_t = A_{t^*} \exp\Big( - (p_d(T) - p_a(t^*))(t - t^*)\Big), \tag{77}$$

where $A_{t^*}$ is the population of alive cells at our chosen time $t^*$. For $p_d(T) - p_a(t^*) > 0$, the above equation shows that the population at time $t^*$ indeed exponentially decays with instantaneous rate $p_d(T) - p_a(t^*)$. Thus, at each moment in time $t^*$, the number of alive cells in the population decreases exponentially with some characteristic (instantaneous) rate $p_d(T) - p_a(t^*)$. Crucially however, this instantaneous rate is monotonically decreasing as we change $t^*$: the population dying continuously slows down over time. This is because the probability of replicating increases towards the probability of dying. To see how the probability of replicating increases towards the probability of dying, reconsider 77. We note that the rate at which the number of alive cells in the population decreases, given by $p_d(T) - p_a(t^*)$, increases from,

$$p_d - p_a(0) = p_d, \tag{78}$$

at time $t = 0$ to (using the lower bound for the probability of replicating 61),

$$p_d - p_a(\tau) \leq p_d - \mu \cdot \frac{A_0 - 1}{Kp_d + A_0 - 1} \to p_d - \mu, \tag{79}$$

at the time of extinction of a large enough population ($A_0 \to \infty$). Thus, the rate at which the number of alive cells in a large population decreases, changes from initially $p_d$ to $p_d - \mu$ when the population goes extinct. This shows that the rate of decay of the number of alive cells in the population is not exponential, and can even halt in cases where the maximum probability of replicating is equal to the probability of dying ($\mu = p_d$), as then, on average and subject to fluctuations, the number of births matches the number of deaths in the population. Thus, our model explains how secretion of glutathion leads to a heavy-tailed decay of the number of alive cells in the population. We study this decay of the population of alive cells in more detail next.

##### B. The decay is not exponentially bounded

Next we will show that the decay of the number of alive cells in the population cannot be bounded with an exponentially decaying function if sufficiently large populations can grow. To this end, we consider the decay of the population of alive cells $A_t$. Our model states that 33,

$$A_{t+1} = A_t \cdot \Big(1 + p_a(t) - p_d\Big), \tag{80}$$

such that the decay of the number of alive cells is given by recursively substituting 80 into itself 40,

$$A_t = A_0 \cdot \prod_{s=0}^{t-1} \Big(1 + p_a(s) - p_d\Big). \tag{81}$$

Now suppose that the number of alive cells $A_t$ in the population decays exponentially or faster. We can then find an upper bound for the number of alive cells in the population with some exponentially decaying function. Let $0 < \alpha < 1$ be some constant and assume that the number of alive cells in the population decays at least exponentially,

$$A_t \leq A_0 \cdot \alpha^t. \tag{82}$$

For $A_t$ to be exponentially bounded, both equations 81 and 82 require that, for all $t > 0$,

$$A_0 \cdot \prod_{s=0}^{t-1} \Big(1 + p_a(s) - p_d\Big) \leq A_0 \cdot \alpha^t. \tag{83}$$

Taking the logarithm and eliminating common terms yields the following condition for the decay of the population of alive cells to be exponentially bounded. For some constant $0 < \alpha < 1$, we need,

$$\frac{1}{t} \sum_{s=0}^{t-1} \log\big(1 + p_a(s) - p_d\big) \leq \log(\alpha), \qquad \text{for all } t > 0. \tag{84}$$

In words, for the decay of $A(t)$ to be exponentially bounded, we need $\log(1 + p_a(t) - p_d)$ to be on average remain smaller than $\log(\alpha)$ for some fixed $1 > \alpha > 0$. Recall the probability of replicating $p_a(t)$ is monotonically increasing in time. Therefore we have $p_a(t) \geq p_a(\tau)$ for any $t \geq \tau$ and some $\tau > 0$. We can now split the function $p_a(t)$ into two time regimes for which we have a lower bound of the value $p_a(t)$: we can use the bound $p_a(t) \geq 0$ for $t < \tau$ and the bound $p_a(t) \geq p_a(\tau)$ for $t \geq \tau$. Hence, we can further bound the left hand side of condition 84 as,

$$\frac{1}{t} \sum_{s=0}^{t-1} \log\big(1 + p_a(s) - p_d\big) \tag{85}$$

$$\geq \frac{1}{t} \sum_{s=0}^{\tau} \log(1 - p_d) + \frac{1}{t} \sum_{s=\tau}^{t-1} \log(1 + p_a(\tau) - p_d) \tag{86}$$

$$= \frac{\tau}{t} \log(1 - p_d) + (1 - \frac{\tau}{t}) \log(1 + p_a(\tau) - p_d). \tag{87}$$

Substitution of the lower bound 87 into 84 yields the following condition for the decay of the population of alive cells to be bounded by an exponentially decaying function,

$$\frac{\tau}{t}\log(1-p_d) + (1-\frac{\tau}{t})\log(1+p_a(\tau)-p_d) \leq \log(\alpha), \quad (88)$$

for all $t > \tau$. For $0 < \alpha < 1$, we can distinguish the following cases:

- $p_d < \mu$: These parameters allow for growth in our model (experimentally, for $T < 40.2°$C). We can choose any $A_0$ such that $p_a(\tau) \geq p_d$ eventually for some $\tau > 0$. Then $1+p_a(\tau)-p_d > 1$ and $\log(1+p_a(\tau)-p_d) > 0$. Then, by substituting into 88,

$$\frac{\tau}{t}\log(1-p_d) < \frac{\tau}{t}\log(1-p_d) + (1-\frac{\tau}{t})\log(1+p_a(\tau)-p_d). \quad (89)$$

  We notice that $\frac{\tau}{t}\log(1-p_d) \to 0$ as $t \to \infty$ for $\tau$ fixed. Thus, for large enough populations and after sufficient time, the condition 88 yields $0 \leq \log(\alpha)$ which cannot be satisfied for $0 < \alpha < 1$. Thus, the decay of the number of alive cells in the population cannot be bounded by an exponentially decaying function when $p_d < \mu$.
- $p_d > \mu$: The probability of dying is higher than the maxium probability of replicating. We cannot obtain $p_a(t) \geq p_d$ and the population is guaranteed to go extinct. At time of extinction $\tau > 0$ we have $p_a(\tau) < p_d$ and such that there exists some $1 > \alpha > 0$ for which the decay of the population of alive cells is exponentially bounded (we can choose $\alpha = 1 + p_a(\tau) - p_d$ in 83). However, the decay rate of the number of alice cells in the population monotonically decreases over time up to the point of extinction, starting with instantaneous rate $p_d(T)$ at time $t = 0$ decreasing to $p_d(T) - p_a(\tau)$ at time of extinction $\tau$.

In summary, the number of alive cells in the population does not decrease exponentially over time. Instead, the decay of the number of alive cells is heavy-tailed, as a result of the probability of replicating approaching the probability of dying. When populations can grow ($p_d < \mu$), this heavy-tailed decay cannot be bounded by an exponentially decaying function. Hence, the change of the number of alive cells in the population cannot be appropriately modeled by an exponential function. Experimentally, we find that the decay of the population of alive cells is indeed heavy-tailed (Figure 3a-b and Figure S7-8), and appropriately modelled by a power-law function.
